# Prostate cancer associated fibroblasts have distinct morphomechanical features that are associated with patient outcome

**DOI:** 10.1101/2025.03.21.644569

**Authors:** Antje Garside, Angela Jacobi, Shivakumar Keerthikumar, Vaibhav Mahajan, Michelle Richards, Birunthi Niranjan, Linda Teng, Nicholas Choo, Gail P Risbridger, Mitchell G Lawrence, Anna V. Taubenberger

**Affiliations:** Center for Molecular and Cellular Bioengineering (CMCB), BIOTEC, TUD, Dresden University of Technology, Dresden, Germany; Leibniz-Institute of Polymer Research Dresden (IPF), Max Bergmann Center of Biomaterials, Dresden, Germany; Melbourne Urological Research Alliance, Biomedicine Discovery Institute, Monash University, Clayton, Victoria, Australia; Peter MacCallum Cancer Centre, Melbourne, Victoria, Australia; Sir Peter MacCallum Department of Oncology, The University of Melbourne, Victoria, Australia; Cabrini Institute, Cabrini Health, Malvern, Victoria, Australia

## Abstract

Tumour development and progression reshape the physical properties of the surrounding tumour microenvironment (TME) including its biomechanical traits. This is driven by a prominent cell type in the TME, cancer associated fibroblasts (CAFs), which increases tissue stiffness via extracellular matrix deposition and remodelling. Currently, it is unclear whether there are also physical changes to CAFs at the cellular level and, if so, how they relate to patient outcome. Here we show that CAFs have distinct morphological and biomechanical features from normal fibroblasts. We examined matched, patient-derived CAFs and non-malignant prostate fibroblasts (NPFs) from 35 patients with primary prostate cancer. Morphologically, CAFs had more aligned stress fibres, and larger and more elongated nuclei, based on quantitative image analysis of confocal microscopy images. In addition, single-cell mechanical measurements using real-time deformability cytometry showed that CAFs are larger and stiffer than NPFs. These changes were consistent across patients and validated with atomic force microscopy. A combined morphomechanical score encompassing these features was significantly associated with patient outcome. In transcriptomic analyses, the score was correlated with microtubule dynamics and a myofibroblast phenotype. Importantly, we also demonstrated that morphomechanical features of prostate fibroblasts are modified by approved treatments for prostate cancer, such as docetaxel, and other small molecular inhibitors, such as axitinib. In summary, changes in cellular morphomechanical properties are a consistent feature of CAFs and associated with patient outcome. Moreover, cellular morphomechanical properties can be therapeutically targeted, potentially providing a new strategy for manipulating the TME to control cancer progression.

## Introduction

Prostate cancer is one of the most common malignancies worldwide with approximately 1.4 million new cases each year^1^. Most patients in high-income countries present with localized disease. Patients with intermediate to high-risk disease receive curative treatments, such as radiotherapy or radical prostatectomy to ablate or remove the tumour^2^. Unfortunately, some patients subsequently develop disease recurrence and progress to advanced prostate cancer. Despite multiple systemic treatments, advanced prostate cancer is ultimately incurable, so it remains a leading cause of cancer-related death. The variability in patient outcomes demonstrates the need to better understand the factors that contribute to disease progression.

In addition to cancer epithelium^3,4,5,6^, the tumour microenvironment (TME) facilitates tumour growth, progression, and therapy resistance^7^. The TME comprises the extracellular matrix (ECM, e.g., collagen, fibronectin, hyaluronan) and various cell types, including immune cells, endothelial cells, and cancer associated fibroblasts (CAFs)^8,9^. CAFs are one of the most abundant cell types within the TME. They lack genetic alterations, but are epigenetically, phenotypically and functionally altered compared with normal fibroblasts^9–13^. There are also different subtypes of CAFs, such as inflammatory CAFs (iCAFs) and myofibroblast-like CAFs (myCAFs) with distinct characteristics^14,15^.

The crosstalk between CAFs and epithelial cells promotes cancer cell transformation, invasion, angiogenesis and immune evasion^10,16,17,18^. One way that CAFs facilitate cancer progression is through excessive secretion and remodelling of ECM, a characteristic feature of reactive stroma^9,19^. *In vitro*, CAFs deposit more aligned ECM^5,20–22^, which promotes more directional migration of epithelial cells^21,23,24^. CAFs are also more contractile, as shown in 3D collagen gels for mammary^22^ and colon CAFs^25^. Higher contractility is associated with increased actomyosin contractility, regulated by Rho GTPases and its effector Rho kinase (ROCK). In addition, Rab21 modulates the levels of active integrins on the cell surface of CAFs, which is needed for force transmission^26^. Increased levels of actomyosin activity are consistent with our previous report that prostatic CAFs have higher cell cortical stiffness compared to patient-matched normal prostatic fibroblasts (NPFs)^27^.

CAF-induced matrix deposition and remodelling modifies the physical properties of the TME, which becomes evident across multiple scales. The properties of bulk tissue are determined by various factors, including ECM stiffness, solid stress, interstitial fluid pressure, and cellular contributions^28^ ^29^. Like other solid tumours, prostate cancer foci are stiffer at the bulk tissue level than normal tissue, allowing these harder masses to be detected during physical examinations, e.g. through digital rectal examination or elastography^30^. Indeed, ECM anisotropy and tissue stiffness have been examined as potential biomarkers using prostate cancer biopsies^31^. Yet in contrast to these properties of bulk tumour tissue, at the single cell level, cancer cells are often more compliant than healthy cells^32^, including for prostate cancer^33^. This difference between the stiffness of bulk tissue and individual cells seems counterintuitive. However, most studies have focused on the mechanical properties of isolated epithelial cancer cells or established cell lines, which only represents one cell type among the complex ecosystem of a tumour.

Therefore, we asked here whether CAFs have a distinctive cellular mechanical phenotype compared to NPFs, and whether it is associated with tumour grade and patient outcome. Using high-throughput cell mechanical analysis (real-time deformability cytometry (RT-DC))^34^, atomic force microscopy (AFM) and imaging, we show that CAFs have a consistently different mechanical and morphological phenotype compared to NPFs^23^. These morphomechanical features are associated with patient outcome, correlated with transcriptional pathways for cytoskeletal remodelling, and can be modulated with pharmacological inhibitors. Therefore, CAFs have distinctive biophysical traits that could be therapeutically targeted.

## Results

### Establishing a cohort of patient-matched CAFs and NPFs

To investigate the phenotype of fibroblasts in the prostate cancer TME, we established a cohort of patient-matched pairs of CAFs and NPFs from 35 patients undergoing a radical prostatectomy (**Figure 1**). We sampled tumour tissue and a contralateral region of non-malignant tissue from each prostate, and digested them to establish primary *in vitro* cultures of CAFs and NPFs (**Figure 1a**). We examined the pathology of each piece of tissue to confirm that they were sampled correctly (**Figure 1b**, **Supplementary Figure 1**). This cohort is typical of patients having surgery for clinically significant prostate cancer (**Supplementary Table 1 & 2**). All patients had grade group 2 prostate cancer or higher (GG2-3: 69%; GG4/5: 29%). In addition, all patients with evaluable data had high-risk prostate cancer based on the D’Amico classification (i.e. ≥4), which combines PSA levels, Gleason score and tumour stage to assess the five-year risk of treatment failure. Accordingly, 29% of patients have since had biochemical or clinical recurrence of prostate cancer. As expected, there was significantly shorter time to biochemical or clinical relapse for patients with higher grade (GG4 or GG5) tumours (**Figure 1c,d**). We used this cohort to comprehensively compare the morphological, biomechanical, and transcriptional features of low passage CAFs and NPFs (median passage 3, range 2-9) (**Supplementary Table 3**).

**Figure 1.**
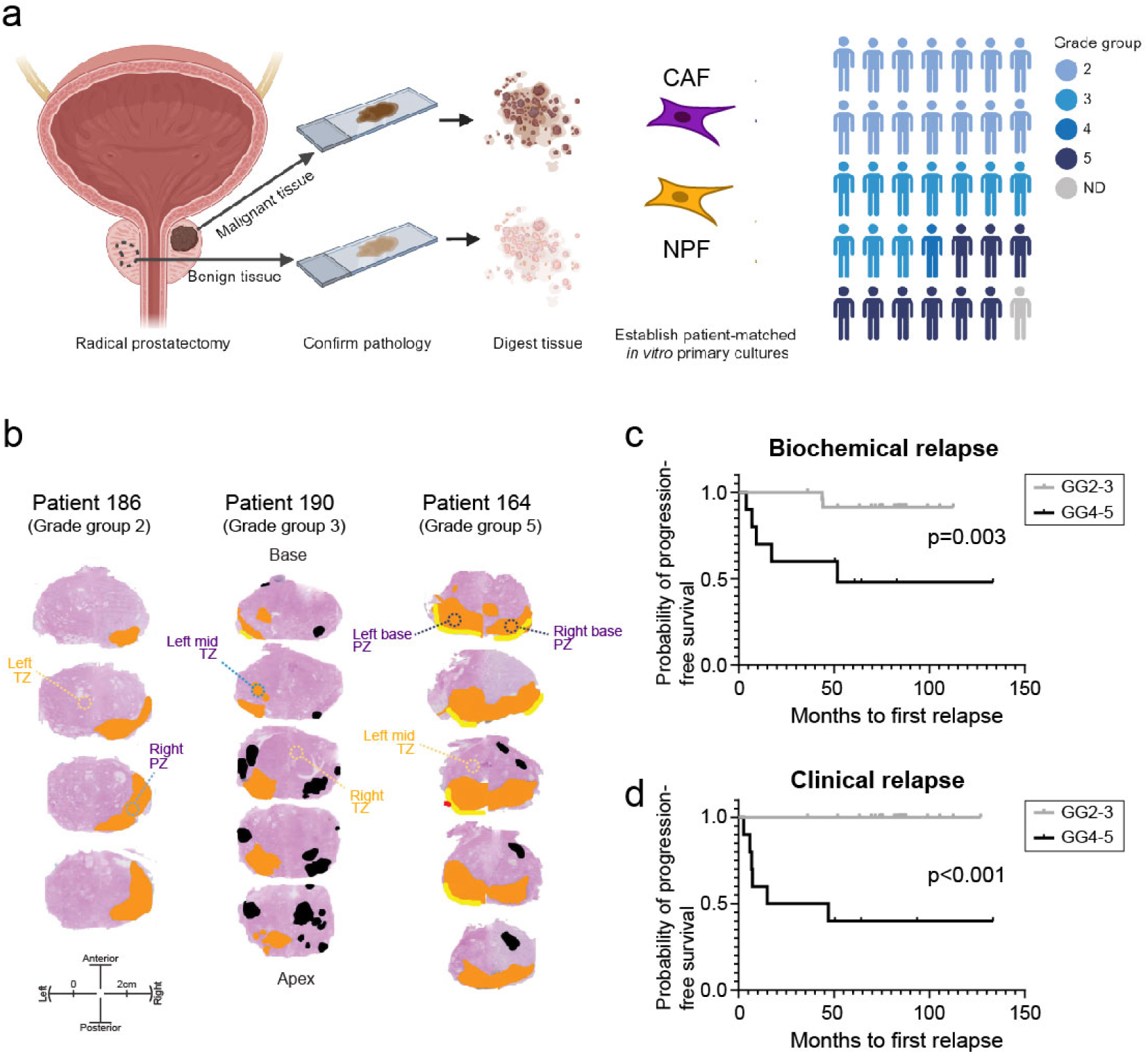
Patient cohort and isolation of CAFs and NPFs. (**a**) Schematic showing isolation of CAFs and NPFs from radical prostatectomy specimens. In total, 35 pairs of patient-matched CAFs and NPFs pairs were analysed, encompassing different grade groups (GG) (14x GG2, 10x GG3, 1x GG4, 9x GG 5, 1 undefined). (**b**) Pathology maps of three representative patients of different grade groups. The index tumour is shown in orange, other tumour foci in black, extra-prostatic extension in yellow and positive margin in red. The regions that were sampled for cancer (purple) and benign tissue (yellow) for each patient are shown (peripheral zone – PZ; transition zone – TZ) (**c**,**d**) Kaplan Meier plots showing time to biochemical (prostate specific antigen, PSA) and clinical relapse for the patient cohort by grade group (note that grade groups GG2 & 3, and GG4 & 5 were pooled, respectively).

### CAFs and NPFs have distinct morphologies

To compare the morphology of CAFs and NPFs, we firstly examined nuclei and the F-actin cytoskeleton in confocal microscopy images of DAPI and phalloidin-TRITC stained cells in confluent 2D cultures (**Figure 2a,b & Supplementary Figure 2**). Across cells from different grade groups of prostate cancer, we noted that CAFs had more unidirectionally aligned actin fibres and elongated nuclei compared to patient-matched NPFs. Quantitative image analysis showed that across the whole cohort, the angles of actin fibres in CAFs typically had narrower distributions compared with NPFs, as indicated by the significantly lower median Gaussian widths (**Figure 2a-f**). This confirmed our previous findings of CAFs having more congruent cell bodies, F-actin and matrix alignment^27,20^. Based on the ratio of F-actin angle distributions, the more uniform cytoskeletal alignment of CAFs versus NPFs is a common feature across patients; the ratio was less than 1 for 27 of 35 patient pairs (**Figure 2g**).

**Figure 2.**
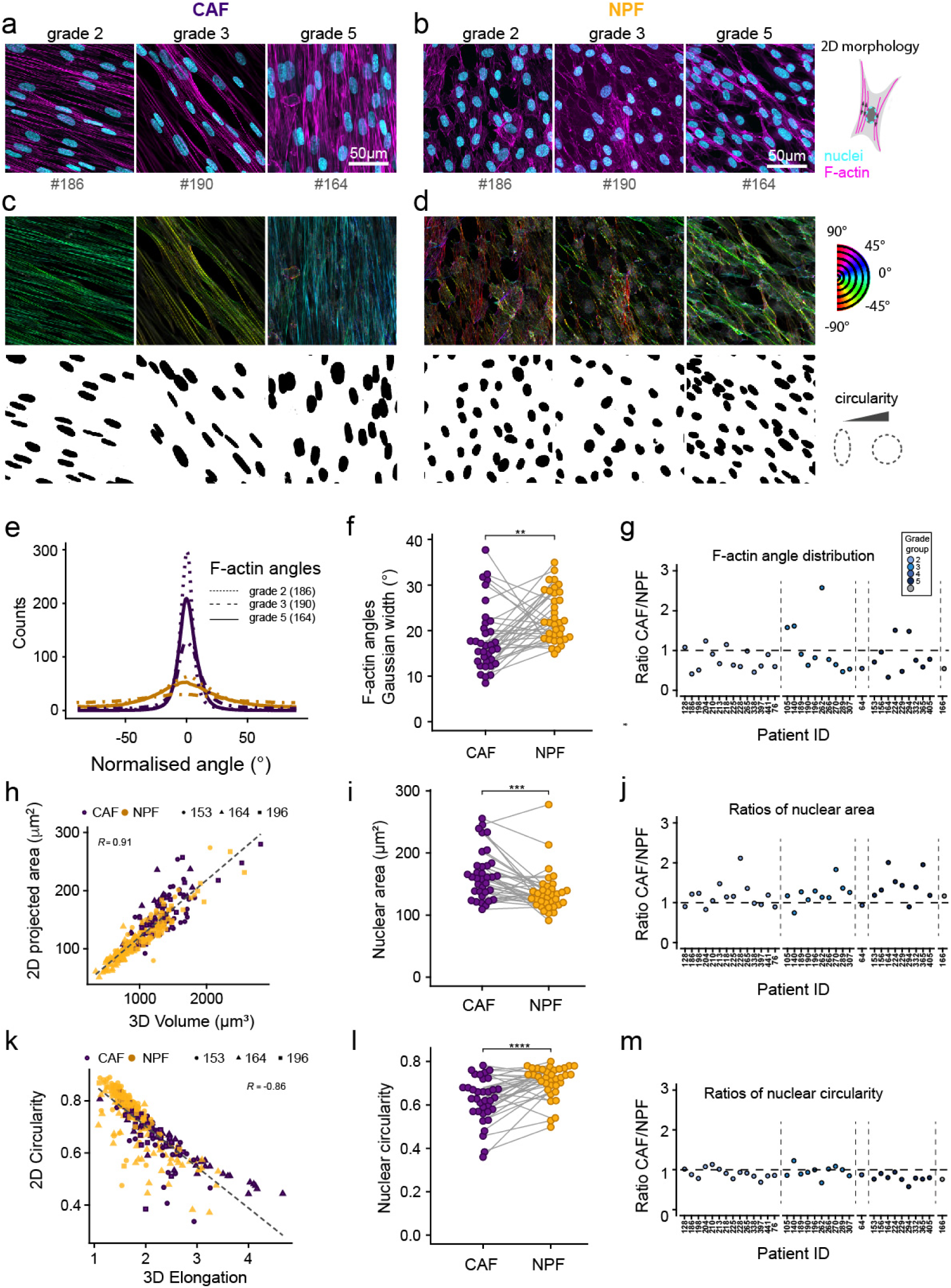
Quantitative morphometric analysis of CAF and NPF cultures. (**a, b**) Representative confocal microscopy images of F-actin (phalloidin-TRITC) and nuclear (DAPI) staining for patient-matched (**a**) CAFs and (**b**) NPFs originating from grade group 2, 3 and 5 tumours. (**c**, **d**) Analysis of F-actin angle distributions and segmented nuclei (**c**: CAF, **d**: NPF) in FIJI for the cultures shown in **a** and **b**. A more uniform colour range indicates higher levels of F-actin fibril alignment. (**e**) Representative histograms showing representative F-actin angle distributions for the patients shown in **a-d**. (**f**) Gaussian width calculated from F-actin distributions (as in **e**) for the whole cohort (n=35 patients). Each patient sample is shown with a separate line with CAFs in purple and NPFs in gold. (**g**) Ratios of Gaussian width measured for CAFs versus matched NPFs. Dot colours indicate tumour grades. (**h**) Correlation of 2D projected nuclear areas and volumes for CAF/NPFs from three representative patients (n=303 nuclei). Spearman correlation coefficients are shown. (**i**) Median projected nuclear area for 35 patient CAF/NPF pairs. Each dot represents the median of an individual patient sample. Lines connect matched CAF/NPF pairs. (**j**) Ratios of projected nuclear areas measured for CAFs and respective NPFs. (**k**) Correlation of 2D circularity and 3D elongation in CAF/NPFs from three representative patients (n=303 nuclei). Spearman correlation coefficients are shown. (**l**) Median nuclear circularity for 35 patient CAF/NPF pairs. Each dot represents the median of an individual patient sample. Lines connect matched CAF/NPF pairs. (**m**) Ratios of nuclear circularity measured for CAFs and respective NPFs. Dot colours indicate tumour grades (**g, j, m**): Results of a Wilcoxon Signed-Rank test are shown. * p<0.05. ** p<0.01, ***p<0.001). See summary of statistical results in **Supplementary Table 4.**

We then characterised nuclear sizes and shapes (**Figure 2a-d, h-m**). Quantitative analysis across all patients revealed significantly greater nuclear projected areas in CAFs (**Figure 2i**), with higher ratios in matched CAFs versus NPFs for 27 of 35 patients (**Figure 2j**). This was not only due to the typically more oblate/flattened nuclear morphology in CAFs, but directly related to increased nuclear volumes as supported by volumetric nuclear analysis using the deep-learning based method Cellpose (**Figure 2h, Supplementary Figures 3 & 4**). Nuclei were on average also significantly more elongated in CAFs, as shown by 2D projections for all patients (**Figure 2l**) and 3D analysis on a subset (**Figure 2k, Supplementary Figure 3 & 4)**. Specifically, CAFs from 26 of 35 patients displayed a less circular nuclear shape compared to NPFs (**Figure 2m**). The variation in morphology within cultures was similar for CAFs and NPFs, with no significant difference in the standard deviations of nuclear area, circularity or angle distributions per patient (**Supplementary Figure 5).** There was also no obvious association between the three morphological parameters and tumour grade (colour coded grade groups in **Figure 2g,j,m**). Altogether, these observations show that CAFs are morphologically distinct from matched NPFs for most patients.

### CAFs are stiffer and larger than NPFs

Given the morphological differences between CAFs and NPFs, we hypothesised that their biomechanical properties also differed. We investigated this using RT-DC to measure the morphorheological features of thousands of individual cells in suspension (**Figure 3a, Supplementary Figure 6 and Movie 1 & 2**). Cells were passed at a defined flow rate (0.32ul/s) through a microfluidic chip mounted on an inverted light microscope. As cells transited through a 30 µm constriction in the microchannel, shear stress deformed their shape. By imaging the cells with a high-speed camera, we determined the deformation and cross-sectional area of each cell in real-time (**Figure 3b**).

**Figure 3.**
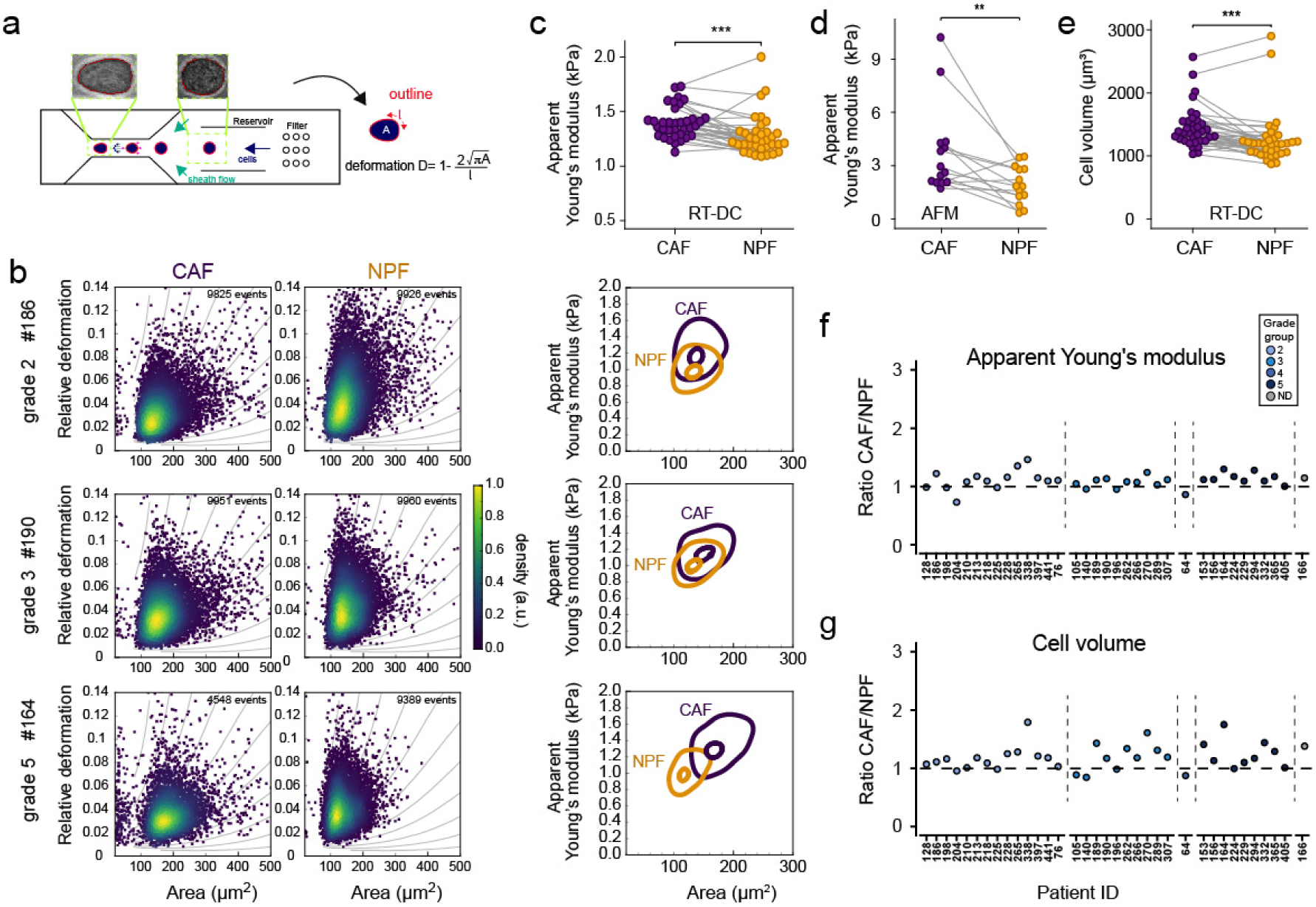
Quantitative biomechanical characterisation of NPFs and CAFs. (**a**) Schematic showing the deformation of cells in the RT-DC microfluidic channel under shear stress. Representative images of a cell before and after entering the channel constriction are shown. From the automated edge detection of cells, projected area and deformation are obtained in real-time. (**b**) Left: Scatter plots of deformation versus area for three representative patients with grade group 2, 3 or 5. Each dot represents the measurement of one cell. Colours indicate the density of overlaid points. Right: Contour plots highlighting 0.5 and 1 density threshold values calculated from the kernel density estimation calculated in Shape-Out 2. (**c**) Median apparent Young’s moduli for CAFs and NPFs from 35 patients. Matched CAF/NPF pairs are connected by lines. (**d)** Apparent Young’s moduli from AFM indentation tests for CAF/NPF pairs in adherent culture (n=14). Ratios of apparent Young’s moduli (CAF:NPF) derived from RT-DC for each patient. Colours indicate tumour grades. (**e**) Median volumes for 35 CAF/NPF pairs. Lines connect matched cultures from each patient (**f,g**) Ratios of median apparent Young’s moduli (**f**) and volumes (**g**) (CAF:NPF) for each patient measured by RT-DC. (**c-e**) Results of a Wilcoxon Signed-Rank test are shown. * p<0.05. ** p<0.01, ***p<0.001. See summary of statistical results in **Supplementary Table 4**.

We found that CAFs had lower deformations and larger sizes compared to NPFs, as evident in RT-DC scatter and contour plots (**Figure 3b**). The lower deformability of CAFs indicates that they are stiffer than NPFs. Accordingly, when we used deformation and cell size to calculate apparent Young’s moduli, a measure of cell stiffness^35,36^, apparent Young’s moduli were significantly greater across the cohort for CAFs versus NPFs (CAF - 1.39kPa versus NPF - 1.28kPa, medians) (**Figure 3c**). Moreover, ratios of apparent Young’s moduli showed that CAFs were consistently stiffer than NPFs across most patients (28 of 35 patients; **Figure 3f**). The difference in stiffness was also reproducible across technical repeats of CAFs and NPFs at different passage numbers (**Supplementary Figure 7**). Since RT-DC measures the properties of cells in suspension, we also used AFM to probe adherent cells in culture for 14 patients. In line with the RT-DC results, CAFs had significantly greater apparent Young’s moduli compared to NPFs (medians 2.78kPa versus 1.84kPa **Figure 3d**), indicating increased cortical stiffness.

CAFs were also significantly larger than NPFs (**Figure 3e**), based on reconstructed cell volumes from the cross-sectional areas measured by RT-DC (medians: CAF - 171µm^2^ versus NPF - 158 µm^2^). The ratios of cell volume confirmed that CAFs were larger than NPFs in 28 of 35 patients (**Figure 3g**). This concords with the larger nuclear volumes of CAFs (**Figure 2f**), especially when assuming comparable ratios of nuclear-to-cytoplasmic volume. There appeared to be greater variability in deformation and size within CAF cultures (**Figure 3b and Supplementary Figure 8**). Accordingly, the standard deviations of apparent Young’s moduli and volumes were greater for CAFs compared to NPFs, demonstrating more variation in these biomechanical features for CAF cultures. There was no apparent association between the relative biomechanical properties of CAFs versus NPFs in each patient with tumour grade (**Figures 3f,g and Supplementary Figure 9**). To rule out possible influences of large changes in cell volumes on apparent Young’s moduli that may not be correctly accounted for with the model used in the Shape-Out software, we compared apparent Young’s moduli for similar volume ranges, which showed consistent differences between CAFs and NPFs (**Supplementary Figure 10**). In summary, CAFs are stiffer and larger than NPFs, both in suspension and in adherent culture.

### A combined morphomechanical score is associated with patient outcome

Since there were consistent differences in the morphology and biomechanical properties of CAFs versus NPFs, we suspected that these features may be correlated with each other, despite intrinsic differences in methodologies and cell states assessed (microscopy of 2D cultures or RT-DC in suspension). Indeed, median cell volume, apparent Youngs’ modulus, nuclear area, nuclear circularity and F-actin Gaussian width were all significantly correlated with each other albeit to varying degrees (**Supplementary Figure 11**). The strongest correlation was between cell volumes and apparent Young’s moduli (Spearman correlation coefficient R=0.8). Given the associations between morphological and biomechanical features, we examined whether combining them by principal component analysis could distinguish CAFs and NPFs. The first two principal components (PC1 and PC2, together explaining about 74% of the variance) efficiently separated CAF and NPFs, with an overlap of 46.4% of the 95% confidence ellipses (**Figure 4a**). Moreover, based on significant differences between PC1 values, this set of features differentiated CAFs and NPFs from both moderate and high grade group cases, with greater separation among high grade group and high D’Amico classification samples (**Figure 4b**).

**Figure 4.**
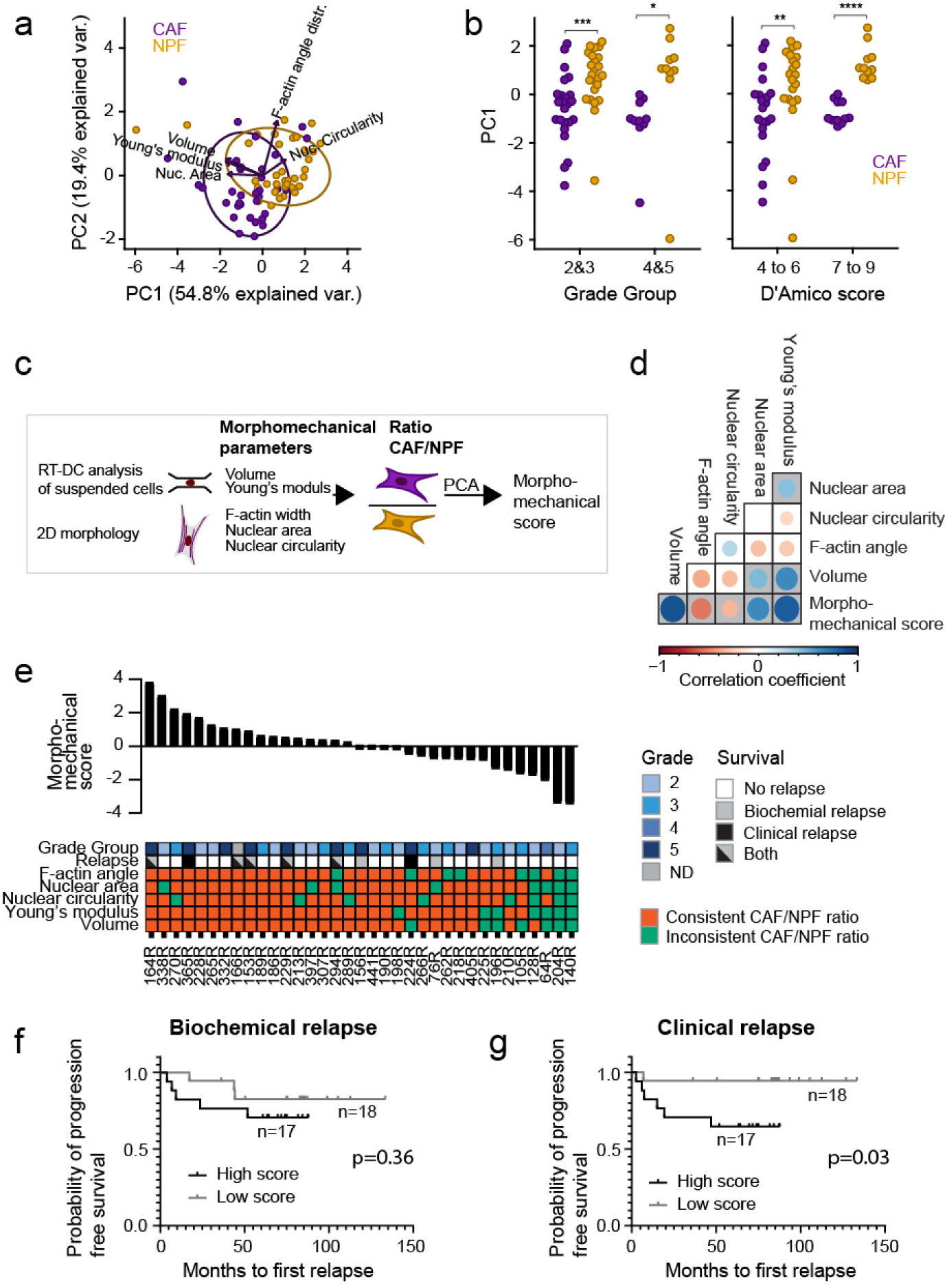
Integration of morphological and mechanical parameters. **(a)** Principal components 1 and 2 calculated from the five morphomechanical parameters volume, apparent Young’s modulus (both RT-DC), and nuclear area and circularity and F-actin angle distribution (2D morphology). Dots represent NPF/CAFs from individual patients (n=35). 95% confidence ellipses are shown. **(b)** PC1 values for CAFs and NPFs grouped by prostate cancer grade group and D’Amico classification. Results of a Wilcoxon Signed-rank test are shown. **(c)** Schematic showing the calculation of the morphomechanical score from ratios of patient-matched medians of the five morphometric and biomechanical parameters. Note that different to (a) ratios of patient-matched CAF/NPF pairs were used to obtain a single score per patient. **(d)** Spearman correlation between individual morphometric and biomechanical parameters and morphomechanical score. The size and colour of the dots represent the correlation coefficients. Significantly correlated parameters have grey backgrounds (*p*<0.05). (**e**) Ranked morphomechanical scores for each patient in the cohort. The heatmap shows Grade group and relapse of each patient and whether the ratios of CAF:NPF morphological and mechanical parameters were consistent with the rest of the cohort as per Fig 2g, j, m and Fig 3f, g. (**f**,**g**) Kaplan-Meier plots for (**f**) biochemical (PSA rise) and (**g**) clinical relapse for patients with morphomechanical scores of below (grey) and above 0 (black). See summary of statistical results in **Supplementary Table 4.**

To enable more in-depth analyses of whether cell features were associated with grade group and patient outcome, we calculated a ‘morphomechanical score’ as a single parameter for each patient. For this, we performed principal component analysis using the ratios of the five morphological and biomechanical parameters in CAFs versus NPFs for each patient (**Figure 4c**). As expected, the ratios of all five parameters were significantly correlated with the morphomechanical score, with the strongest association for volume and Young’s modulus (**Figure 4d)**. Of note, patients with higher morphomechanical scores had greater and more consistent differences between the features of CAFs and NPFs compared to patients with lower scores (**Figure 4e**). Also, we noted among patients with higher morphomechanical scores a larger proportion of high-grade group patients and clinical relapse cases (**Figure 4e**). Indeed, Kaplan-Meier analysis revealed that patients with positive morphomechanical scores had significantly shorter time to clinical relapse (but not PSA relapse) (**Figure 4f,g**). Some individual parameters showed the same trend with patient follow-up (**Supplementary Figure 12**). Overall, this shows that the combined measurements of RT-DC and morphometric analysis distinguishes between CAFs and NPFs, and that patients with greater differences have poorer relapse-free survival.

### Transcriptomic profiles are correlated with morphomechanical features

We examined whether the morphometric and biomechanical features of CAFs and NPFs are associated with specific gene expression differences using RNA sequencing data (**Figure 5a**). First, we performed a principal component analysis which separated most CAFs and NPFs based on the top 5000 highly variable genes (**Figure 5b**). This confirms the consistent transcriptional differences between cells across patients. Some samples were outliers, such as CAF204R, which was notably also an outlier based on F-actin distribution, nuclear area, nuclear circularity, and apparent Young’s modulus (**Figure 2 & 3**), and had one of the lowest morphomechanical scores. Next, we calculated the relative expression of each gene (gene ratio) between CAFs and NPFs from each patient and compared it to respective morphomechanical scores (**Figure 5a**). We identified 49 genes correlated with the morphomechanical score (**Supplementary Table 5**). Hierarchical clustering of these genes separated CAFs from NPFs, again with outliers such as CAF204R (**Figure 5c**). The top correlated genes included *NAV3* (involved in microtubules regulation), *CTHRC1* (Wnt pathway and collagen matrix deposition), *SLC14A1* (volume control), *MYOCD* (regulating expression of serum response factor (SRF)), *ARHGAP28* (RhoA GTPase), *IGFBP5* (lipid metabolism), *MAPK10* (ErbB signaling), among others (**Figure 5c**). Many of the 49 genes were also correlated with individual morphological and biomechanical features (**Figure 5d**). Indeed, 34 genes were significantly correlated with volume and/or apparent Young’s modulus and they also separated CAFs from NPFs in hierarchical clustering (**Supplementary Figure 13**). This validates that these two biomechanical features are robust at differentiating CAFs and NPFs and contribute to the morphomechanical score.

**Figure 5.**
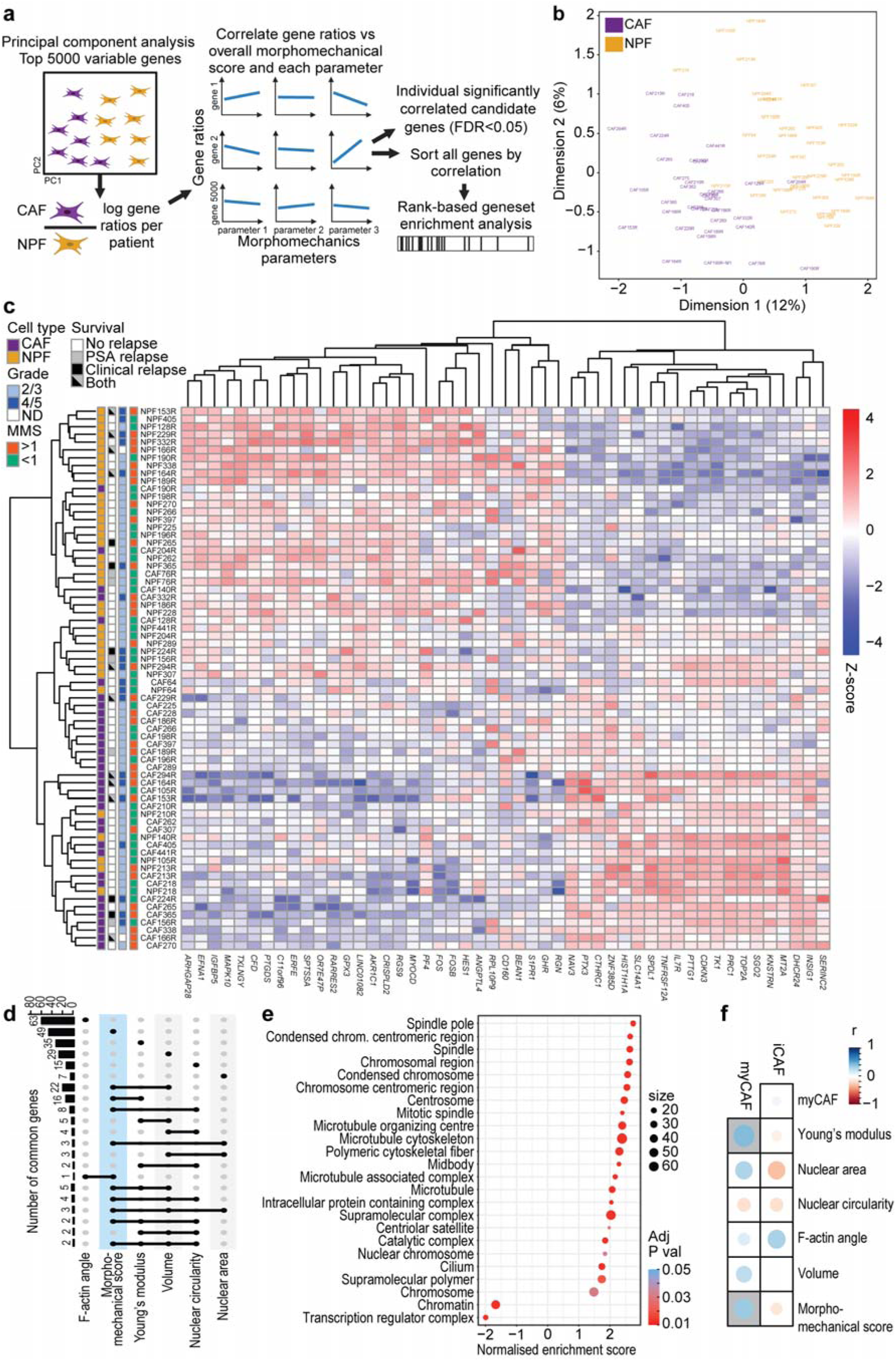
The morphomechanical score is correlated with transcriptomic differences between CAF and NPFs. **(a**) Schematic showing the strategy for comparing transcriptomic data to the morphomechanical score. Based on principal component analysis, 5000 highly variable genes between CAFs and NPFs were identified. The ratios of log transformed read counts (CPM) for each gene in CAFs versus NPFs from each patient were compared to the morphomechanical scores. Significantly related genes (FDR<0.05) were ranked by Spearman’s correlation and used for gene set enrichment analysis. (**b**) Principal components PC1 and PC2 for the 70 CAF and NPF samples from 35 patients. (**c**) Heatmap of complete linkage hierarchical clustering of 49 genes correlated with the morphomechanical score. Colours denote scaled gene expression levels (Z-scores) across the heatmap. (**d**) Upset plot showing the number of genes with a significant correlation with each morphological and biomechanical parameter and common genes that are correlated with multiple parameters. (**e**) Normalised enrichment scores for cellular component pathways with significant differences in GSEA of genes ranked by their correlation with morphomechanical scores. (**f**) The correlation between iCAF and myCAF signatures with morphomechanical score and individual features. The size and colour of each circle represent the Spearman correlation coefficient. Grey backgrounds indicate significant correlations (*P*<0.05). See summary of statistical results in **Supplementary Table 4.**

To identify pathways that differ between CAFs and NPFs, we ranked genes based on their correlation with morphomechanical scores and performed Gene Set Enrichment Analyses (GSEA) (**Supplementary Table 6**). The common themes among the top-ranked cellular component gene sets were enrichment of microtubules, spindle and chromosome related signatures (**Figure 5e**). This suggests that differences in cytoskeleton and nuclear architecture are correlated with the distinct morphological and biomechanical features of CAFs and NPFs. CAFs can be divided into different subtypes, including inflammatory (iCAF) and myofibroblastic (myCAFs) cells^37^, so we examined whether the relative enrichment of transcriptional signatures for these phenotypes between CAFs and NPFs for each patient was associated with morphomechanical parameters (**Figure 5f & Supplementary Table 7**). There were no significant correlations with the iCAF signature. However, the enrichment of the myCAF signature was significantly positively correlated with morphomechanical scores (r=0.36, *P*=0.038) and the apparent Young’s moduli (r=0.43, *P*=0.004). This association with cell stiffness is consistent with myCAFs having a more contractile phenotype^37^.

### Morphometric and biomechanical features of prostate fibroblasts respond to TGF-**β** signalling

Having observed a correlation of the morphomechanical score with myCAF signatures, we explored whether known inducers of the myCAF phenotype would affect morphometric and biomechanical features of CAFs and NPFs. Therefore, we modulated the activity of the TGF-β pathway, which regulates CAF-like phenotypes in fibroblasts^9,38^. TGF-β is a known inducer of a CAF and myofibroblast phenotype^39,40^ and a recent report has shown increased Young’s moduli of pancreatic CAFs after TGF-β treatment as measured with AFM^41^. We treated CAFs and NPFs from three patients for 48 hrs with recombinant TGF-β1 to activate the pathway or A83-01, an ALK4/5/7 inhibitor, to block the pathway. TGF-β1-treated CAFs, but not NPFs, had enlarged nuclei in 2D cultures (**Figure 6a-c**), while nuclear circularity and F-actin angle distribution did not change (**Figure 6d-e**). We also inspected αSMA staining in CAF and NPF cultures (**Figure 6a,b**) but only found single cells with a strong signal, with considerably varying levels across different patients (**Supplementary Figure 14**). Quantitative co-localisation analysis with F-actin revealed only minor changes with TGF-β pathway manipulation (**Supplementary Figure 14**). Cells were stiffer after TGF-β1 treatment based on changes in the apparent Young’s moduli measured in adherent cells (AFM) (**Figure 6f**) and in suspension (RT-DC) (**Figure 6g,h**). Conversely, blocking by A83-01 decreased cell stiffness particularly in CAFs and less consistently in NPFs. TGF-β1 treatment also modulated cell volumes (**Figure 6i**), with more pronounced changes induced in CAFs. Altogether, these results demonstrate that the morphological and biomechanical features of prostate fibroblasts in baseline culture can be further modulated by targeting the TGF-β pathway.

**Figure 6.**
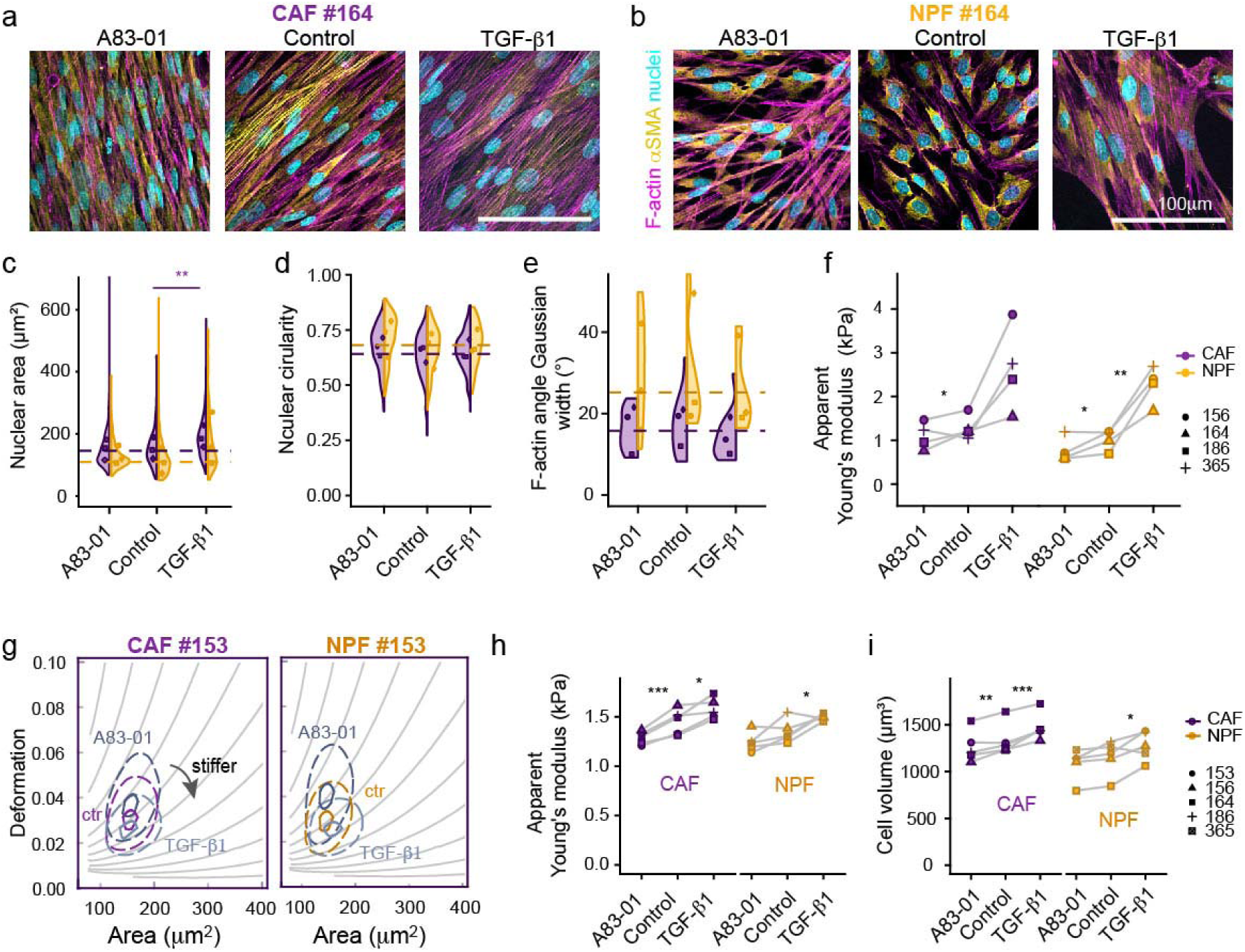
Modulating TGF-β signalling changes the morphological and biomechanical features of CAFs and NPFs. **(a,b)** Representative confocal microscopy images of CAFs (**a**) and NPFs (**b**) stained for F-actin, αSMA and nuclei after 48hrs treatment with 10ng/ml TGF-β1 or the TGF-β inhibitor A83-01 (and vehicle controls). (**c-e**) Half-violin plots showing distributions and medians per patient after quantitative analysis of nuclear area (**c**), nuclear circularity (**d**) and F-actin angle distribution (Gaussian width, **e**) for three representative patients. (**f**) Median apparent Young’s moduli obtained by AFM indentation tests on adherent CAF/NPFs for three patient pairs in dependence of TGF-β1 and A83-01 treatment (compared to vehicle controls). (**g**) Representative contour plots from RT-DC analysis of suspended CAFs and NPFs, showing vehicle controls and cells treated with TGF-β1 or A83-01. (**h,i**) Median apparent Young’s moduli (**h**) and volumes (**i**) obtained by RT-DC for three patient pairs. (**c-f,h,i**) Results from a linear mixed model statistical analysis are shown (* *P*<0.05. ** *P*<0.01, \*\*\**P*<0.001). Different donors (n=4-5) are highlighted by symbols/patient numbers. See summary of statistical results in **Supplementary Table 4.**

### The morphomechanical phenotype of CAFs and NPFs is sensitive to drugs targeting microtubules and PDGF/VEGF receptor signalling

We next tested the effect of a panel of cytoskeletal-targeting drugs, known anti-cancer drugs and statins (previously also associated with CAF contractility) on the morphological and mechanical properties of CAFs and NPFs. While acute treatments (30-45min) in suspension were used to test cytoskeletal contributions to cortical stiffness, longer treatments (24-72hrs) were used to mimic clinical treatment protocols. We confirmed that the doses of each inhibitor were not toxic at these timepoints (**Figure 7a**, **Supplementary Figure 15**). Specifically, we treated cells with compounds targeting microtubule turnover (docetaxel, nocodazole), Rho-associated kinase (ROCK) (Y27632), Rac/Cdc42 (MBQ-167), and Wnt pathway (CCT251545). In addition, we tested clinical drugs, e.g. docetaxel, the PDFR and VEGF receptor tyrosine kinase inhibitor Axitinib, recently in clinical trials for advanced prostate cancer; simvastatin (blocking cholesterol production) and enzalutamide, inhibitor of androgen receptor signalling (standard of care for advanced prostate cancer). After drug treatment, we used microscopy to examine changes in nuclear features (**Figure 7b,c**) and RT-DC to examine stiffness and volume (**Figure 7d,e**). Axitinib had the greatest effect on the cell mechanical properties and morphology of CAFs and NPFs. CAFs and NPFs were significantly stiffer and larger after Axitinib treatment, as measured by RT-DC. In addition, cells had a more spread-out morphology and more pronounced stress fibrils (**Figure 7a**). Notably, disrupting microtubule dynamics with docetaxel had varying effects, depending on the time and setting of treatment (adherent vs suspended). Short-term docetaxel treatment in suspension had little effect on the mechanical phenotype, whereas 24 hrs treatment of adherent cells induced morphological changes and cell stiffening (**Figure 7c-f, Supplementary Figure 16**). In contrast, nocodazole reduced cellular stiffness, in keeping with its ability to destabilise microtubules^42^. ROCK inhibition with Y27632 had distinct, but reproducible, effects on CAFs and NPFs from different patients. While Y27632 treatment in suspension prior RT-DC had mixed effects in different patients, in AFM CAFs and NPFs became consistently more compliant after 30 min treatment (**Supplementary Figure 17**). The rac inhibitor MBQ-167 reduced the apparent Young’s moduli of NPFs but had no significant effect on CAFs. For other drugs, such as the Wnt inhibitor as well as simvastatin, there were changes in morphology but not the mechanical phenotype (**Figure 7c,d**). Overall, these results demonstrate that the morphological and mechanical features of prostate fibroblasts are modified by tool compounds and commonly used therapies for prostate cancer.

**Figure 7.**
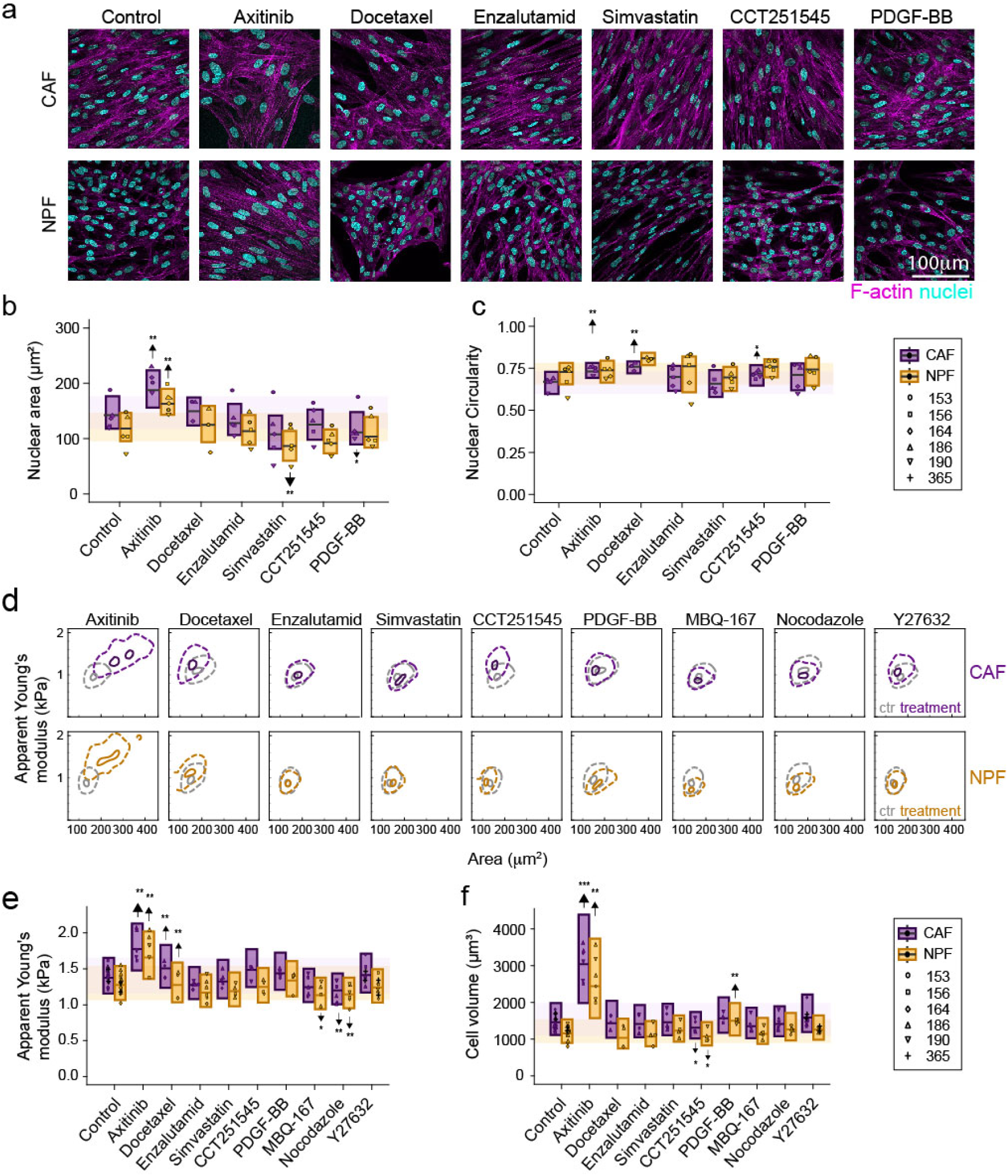
Effects of clinical drugs and inhibitors targeting cytoskeleton or specific pathways on morphological and mechanical features. **(a)** Representative images of CAFs and NPFs treated with different drugs. F-actin (magenta) and nuclei (DAPI, cyan) are shown. Cells were treated with compounds targeting microtubule turnover (5 nM docetaxel, 24 hrs adherent), Rac/Cdc42 (0.5 µM MBQ-167, 2 hrs adherent); Wnt pathway (CCT251545, 72 hrs adherent); PDFR and VEGF receptor tyrosine kinases 1,2,3 (1 µM axitinib, 72 hrs adherent; recently in clinical trials for advanced prostate cancer); cholesterol production (1 µM simvastatin, 24hr adherent); and the androgen receptor (10 µM, enzalutamide, 72hrs adherent, standard of care for advanced prostate cancer). Cells remained viable under these conditions (see also **Supplementary Figures 15**). **(b,c)** Boxplots and overlaid dot plots showing effects of above-mentioned drugs on 2D nuclear area (**b**) and circularity (**c**). Boxplots represent whole data sets, dot plots medians per patient. (**d**) Representative contour plots from RT-DC analysis showing apparent Young’s moduli versus area for controls and drug-treated CAFs (above) and NPFs (below) (**e,f**) Box plots and overlaid dot plots showing apparent Young’s moduli (**e**) and volumes (**f**) measured from RT-DC experiments after drug treatment as outlined in (**a**). Boxplots represent whole data sets, dot plots medians per patient. In addition to morphological analysis (**b**,**c**), the acute effect of cytoskeletal drugs on cells on suspended cells was tested, namely, 5 µM nocodazole and docetaxel (45 min) and 10 µM of the ROCK-inhibitor Y27632 (30 min). (**b-e**) Results from statistical analysis using linear mixed models are given, comparing drug-treated samples to respective vehicle controls. See summary of statistical results in **Supplementary Table 4**.

## Discussion

It is well known that there are changes in the biomechanical features of prostate cancer at the tumour scale. Indeed, the greater stiffness of tumour foci compared to benign tissue is the basis of digital rectal examinations being used to palpate for the potential presence of cancer in patients. In contrast, little is known about the biophysical traits of prostate cancer at the cellular scale, especially within the stroma. Using a cohort of matched CAFs and NPFs from patients undergoing surgery for primary prostate cancer, we demonstrated consistent differences in morphology and biomechanics. Overall, CAFs are larger and stiffer, with large, elongated nuclei and more aligned actin filaments. Patients with greater differences in these features, as captured by the morphomechanical score, have poorer outcome. Therefore, the biomechanical properties of the TME appear to evolve at both the macro- and micro-scale in prostate cancer.

Beyond the molecular differences between CAFs and normal fibroblasts, recent studies have described their distinct biophysical attributes, including cell shape, motility, contractility and mechanics, albeit only for a limited number of unmatched patients and in different tumour entities^26,41,43^. Here, we focused on the morphological and mechanical characteristics of prostatic CAFs. Our robust cohort had long clinical follow up and patient-matched cells providing controls for each patient. We used early passage cultures (median passage 3) where possible, since both mechanical and morphological features of cells changed with passage number (particular beyond passage 5), consistent with a previous study of murine fibroblasts^44^.

A notable feature of CAFs is their larger nuclear and cellular volumes compared to NPFs. This is consistent with our pilot study^27^ and another study on pancreatic CAFs^41^. The reasons for this marked cell volume increase of CAFs versus NPFs is still speculative. It is unlikely that these changes in volume are due to differences in cell cycle progression, since we also see them in confluent cultures^27^. In addition, the distributions of cell volumes do not overlap as would be expected for shifting proportions of larger G2 and smaller G1 cells^45^. More likely explanations for the increased volume of CAFs include differences in growth factor signalling and cell growth rate, cytoskeletal regulation, or osmotic regulation^45,46^. This could be explored in future studies using transcriptomic datasets that might instruct more mechanistic explorations ^47^.

CAFs also exhibit greater cell spreading and more aligned F-actin cytoskeletal arrangements compared to NPFs. This concords with previous studies of CAF from prostate, colon and breast^23,27,22^. The increased spreading of CAFs was also consistent with their more elongated nuclei, likely due to forces exerted by the cytoskeleton on the nuclear structure^48^. More elongated nuclei are associated with increased mechano-transduction and translocation of transcriptional regulators such as YAP^49^. This fits with previous reports showing YAP activation in breast CAFs^22^.

The differences in nuclear and concomitant cell elongation and F-actin distribution indicate that CAFs are more contractile than NPFs. Accordingly, CAFs were stiffer than NPFs, which is indicative of higher cortical tension, in particular when measured by AFM on adherent cells as also previously shown^27^. In contrast, a previous AFM study of pancreatic CAFs reported similar or less stiffness compared to normal fibroblasts^41^, which may be related to the different tissue origin of CAFs or, more likely, to the use of non-matched samples. Nevertheless, another indication that prostatic CAFs were more contractile than matched NPFs was the correlation between the morphomechanical score and transcriptional signature for myCAFs. Indeed, in ovarian cancer models, myCAFs are more contractile and contribute to increased tissue stiffness^50^. It is notable that these mechanical and morphological differences between prostate CAFs and NPFs are maintained for several passages after removal from their native environment. This may be a manifestation of the consistent epigenomic differences between CAFs and NPFs^12,13^. It could also be due to a self-reinforcing feed-forward loop that maintains a contractile phenotype, as described for CAFs from squamous cell carcinoma^22^. There, ROCK inhibition of CAFs reduced contractility and causing a long-lasting reversion of the CAF phenotype. Altogether, this suggests that greater contractility is a consistent and persistent feature of CAFs, and that it is potentially targetable.

Despite intrinsic differences between AFM and RT-DC, both techniques revealed differences between the mechanical properties of CAFs and NPFs. While AFM on adherent cells is largely influenced by the F-actin cortex, RT-DC probes cells in suspension and at shorter time scales (millisecond vs second range), where stress fibrils are naturally absent and the cortex is remodelled during cell detachment^51^. Of note, while AFM indentation tests show a significant decrease in apparent Youngs moduli for both CAFs and NPFs when treated with ROCK inhibitor, such effect was not seen by RT-DC. This might be explained by different effects of myosin inhibition in suspended versus adherent state-as previously reported^52^.

As a single cell technique, RT-DC can also reveal mechanical heterogeneity of CAFs and NPFs. Indeed, distinct subpopulations became evident after drug treatment, including with Axitinib. Since different subtypes have been proposed as prognostic markers, single cell mechanical phenotyping with RT-DC could present a simple and label-free method to detect mechanical subtypes, for example by combining it with machine learning algorithms to analyse the optical images obtained for each cell. This might be beneficial to better understand the nature of CAF heterogeneity.

The morphological and mechanical attributes of CAFs and NPFs should be inherently linked to their gene expression profiles. Indeed, targeting the cytoskeleton also altered the morphological and mechanical features of CAFs and NPFs, which was also consistent with our transcriptomic analyses: the morphomechanical score was highly correlated with enrichment of microtubules, spindle and chromosome related signatures. The Rac inhibitor MBQ-167, nocodazole and docetaxel all changed the mechanical phenotype of both CAFs and NPFs. This is notable, because docetaxel is used for patients with advanced prostate cancer and high tumour burden. The increased stiffness of CAFs and NPFs treated with docetaxel is likely due to microtubule stabilisation, since taxanes binds to tubulin subunits and cause a more rigid microtubule structure^53^. Taxanes also increase the stiffness of oesophageal squamous carcinoma cells^54^ and ovarian cancer cells^55^, although not all studies agree with this trend^56^. The most striking change in the mechanics of CAFs and NPFs was induced by Axitinib, a PDGFR/VEGFR inhibitor that has been used in clinical trials for renal^57^ and prostate cancer^58^. This may also be due to cytoskeletal remodelling, since it is regulated by PDGF^59^. Besides a clear stiffening effect, axitinib was recently shown to induce a proto-myofibroblast-like phenotype in pericytes with increased αSMA levels^60^, which is in line with our mechanical data.

A previous study has shown a role for statins including simvastatin in blocking collagen gel contraction by head and neck cancer-derived CAFs^26^. Interestingly, we did not detect an effect of simvastatin mechanical cell properties by RT-DC, albeit an induced change in NPF nuclear shapes in 2D cultures. Since the effect of simvastatin on the cells’ mechanical phenotype was previously attributed to its inhibitory effect on RhoGTPase prenylation, the lack of mechanical alterations in our study may be again attributed to probing cells in suspended rather than adherent state.

We also showed that the morphomechanical features of prostatic CAFs are associated with patient outcome. Previous studies have explored various features of the TME as diagnostic tools, including matrix content and bulk tissue biomechanics^31^. For example, collagen structure anisotropy, quantified using label-free second harmonic imaging, scaled with Gleason score and was, therefore, proposed as a potential biomarker^61^. In our study, the individual morphomechanical parameters were not associated with grade group; however, the apparent Young’s modulus, volume, and combined morphomechanical score were significantly associated with clinical relapse-free survival. This suggests that morphomechanical features and histology provide different measures of tumour risk, so could be complementary parameters. Further cohorts could be used to test this idea. RT-DC is well suited to this, since it is a rapid, label-free and time-efficient approach. Ideally, the morphomechanical features could be evaluated using single cell suspensions from fresh biopsy samples, circumventing the need for *in vitro* culture. A challenge, however, would be to discriminate between cell types given the lack of unique CAF markers.

In summary, across different patients, prostatic CAFs have distinct mechanical and morphological characteristics compared to matched NPFs. Future pan-cancer studies could be used to investigate the morphomechanical features of CAFs from other tumour types and determine whether they are also associated with patient outcomes. Moreover, the morphomechanical features of CAFs can be modulated by small molecules, opening a new direction for screening therapies that target the tumour stroma to disrupt cancer progression.

## Materials & Methods

### Establishing and culturing patient-derived fibroblasts

Written informed consent was obtained from patients to collect fresh tissue from radical prostatectomy specimens according to human ethics approvals from Monash Health and Monash University (1636 and 36762), Cabrini (03-14-04-08) and Epworth HealthCare (53611)With assistance from a board-certified pathologist, tumour tissue and matched benign tissue (from a distant region of the same prostate) were collected from each patient sample, as previously described^12,62^. The pathology of these regions of tissue was confirmed at the time of sample collection with rapid haematoxylin and eosin staining. To establish primary cultures of fibroblasts, tissues were digested with 225 U/mL collagenase and 125 U/mL hyaluronidase as described^12,62^. Small pieces of undigested tissue were retained and formalin-fixed and paraffin embedded. These tissues were re-examined by a uro-pathologist and patient samples were only included in this study if the pathology of the cancer and benign tissues was re-confirmed. From the digested tissues, cell suspensions were plated in RPMI 1640 media (Gibco, ThermoFisher) containing 5% foetal bovine serum (FBS; Gibco, ThermoFisher), 1nM testosterone (Sigma-Aldrich), 10ng/mL human fibroblast growth factor 2 (Miltenyi Biotec), 100 U/mL penicillin, and 100 μg/mL streptomycin (ThermoFisher). Cells were routinely grown at 37°C in 5 % CO_2_. The cultures were confirmed to be free of mycoplasma. Primary fibroblasts were used in experiments between passage 2 and 8.

### Fluorescence staining of nuclei and F-actin fibres

For subsequent staining, 30 000 cells were seeded on Thermanox coverslips (Ø 13 mm, Nunc, ThermoScientific) in 24-well plates and cultured for 14 days. Coverslips were washed with PBS (w/ Mg, Ca) and fixed for 5 min with 4 % (w/v) paraformaldehyde/PBS. After 5 min permeabilization with 0.2 % (v/v) Triton X-100/PBS, cells were stained for 1 h with 5 µg/mL 4′,6-diamidino-2-phenylindole (DAPI, ThermoFisher) and 0.4 µg/mL Phalloidin-TRITC (ThermoFisher) in 2 % (w/v) BSA/PBS at room temperature. Cells were rinsed with PBS and briefly dipped in double-distilled water before mounting with Aqua-Poly/Mount (Polysciences) and imaged with a confocal microscope (Zeiss LSM 780, 40x objective).

### Immunofluorescence staining of **α**SMA

Cells were fixed for five minutes with 4 % (w/v) paraformaldehyde/PBS at room temperature. After permeabilization for five minutes with 0.2 % (v/v) Triton X-100/PBS, samples were blocked for 45 min in 2 % (w/v) BSA/PBS. Primary antibody (αSMA, Sigma A5228, 10 µg/ml, mouse) was diluted in Antibody Dilution Buffer [1 % (w/v) BSA/0.2 % (v/v) Triton X-100/PBS] together with DAPI and Phalloidin-TRITC as described above. Cells were incubated in this solution in a humid chamber at 4°C overnight. On the next day, cells were brought to room temperature and washed 3x5 min with PBS. Secondary antibody (Cy5 anti-mouse IgG, Jackson Immuno Research, 5 µg/ml) was diluted in Antibody Dilution Buffer and cells incubated in a humid chamber at room temperature for 2 h. Subsequently, cells were washed 3x5 min with PBS, rinsed with double-distilled water and mounted with Aqua-Poly/Mount (Polysciences) before imaging with a confocal microscope (Zeiss LSM 780, 40x objective).

### Image analysis

Image analysis was performed with Fiji^63^ (<u>version 1.53o</u>) and the OrientationJ plugin (^64,65^). Briefly, nuclear area and circularity were determined utilising the Analyze Particles function of the ImageJ software with circularity defined as circularity=4π(area/perimeter^2). Representative images of F-actin angles as hue-saturation-brightness colour-coded fibre angles were created with the OrientationJ Analysis function. Angle distributions were calculated using the OrientationJ Distribution function with a local Gaussian window of σ = 2 pixels. The angles were normalised so that the distribution maximum lies at 0°. The width of angle distribution was calculated after fitting to a Gaussian function using an Igor (version 6.3) script.

For comparison of nuclear volume with its projected area, Z-stack images were acquired with 40x objective and 0.6 μm step size on a Nikon Ti-E confocal microscope with spinning disk. For 2D cell area, maximum intensity projections were used and analysed as mentioned above. For 3D segmentation, first the images were processed to remove background by the rolling ball method (radius = 50 px) in Fiji. 3D volumes of the same images were calculated with a custom model on Cellpose version 2.2^66^. The human-in-the-loop pipeline of the Cellpose GUI was used to create the custom model (starting with the pretrained model ‘nuclei’). Z slices containing both nuclei and background were used for training. Incorrect segmentations of overlapping nuclei or nuclei on the edge were cleaned out with napari (version 0.4.17). Nuclear volumes and shape descriptors were calculated from the segmented masks with the 3DSuite ImageJ plugin^67^. Colocalization analysis of αSMA and F-actin was performed with the BIOP JACoP plugin for Fiji (https://github.com/BIOP/ijp-jacop-b). One perinuclear region of interest per cell was analysed with the automatic threshold method Otsu for both channels.

### Atomic force microscopy

30 000 fibroblasts were seeded onto 13 mm diameter Thermanox coverslips (Nunc, ThermoScientific) and cultured as described above. After 14 days, coverslips were mounted in a 35 mm diameter petri dish (TPP) with vacuum grease and CO_2_-independent medium (Gibco) was added for subsequent measurements. Indentation experiments were performed with a Nanowizard 4 (JPK/Bruker, Berlin) AFM mounted on a Zeiss Axio Observer light microscope (Zeiss, Jena) equipped with a petri dish heater (JPK/Bruker). Prior to measurements, Arrow-TL1 cantilevers (nominal spring constant 0.035–0.050 N/m, Nanoworld) that had been modified with polystyrene beads (Ø 5µm, microParticles GmbH) were calibrated using the thermal noise method implemented in the AFM software. For each cell type, three regions of interest were probed using grids of 5x5 points equally spaced over an area of 100 µm x 100 µm. A piezo speed of 5 µm/s and relative set point of 2.5 nN were chosen. Experiments were conducted at 37°C. Force-distance curves were analysed using the data processing software (version 6.1.203, JPK/Bruker) using the Hertz/Sneddon model^68^ for a spherical indenter and assuming a Poisson ratio of 0.5.

### Real-time deformability cytometry (RT-DC)

For real-time deformability cytometry, 10^6^ cells were seeded in a T75 flask 48 h before measurement. Fibroblasts were washed with PBS and detached with TrypLE Express (Gibco). Thereafter, cells were pelleted by centrifugation, resuspended in 0.8mL CO_2_-independent medium (Gibco) with 5 % (v/v) FBS and kept in a suspension culture plate (Greiner) for 1 h at room temperature. Measurements were performed as previously described^34^. 0.5 % (w/v) methylcellulose in PBS with 10 % (v/v) Leibovitz’s L-15 Medium and 100 Kunitz Unit/mL DNase I (Sigma-Aldrich) was utilized as sample and sheath fluid. Cells were flowed through a microfluidic chip with a 30 µm channel constriction (Zellmechanik Dresden) with a flow rate of 0.32 µL/s (sample flow 0.08 µL/s, sheath flow 0.24 µL/s). Data were analysed and apparent Young’s moduli were computed with ShapeOut 2.10.0 (https://github.com/ZELLMECHANIK-DRESDEN/ShapeOut2). Objects within an area range of 50-1000 µm^2^ and a convex to measured area ratio within 1.00 and 1.05 were analysed after applying respective filters (area, porosity filter), thereby removing mostly larger cell clusters and debris. For each dataset, medians of the parameters area and apparent Young’s modulus were calculated for further analysis. Typically, 10 000 cells were recorded in each experimental run.

### RNA sequencing

Total RNA was extracted from CAFs and NPFs using the RNeasy kit (Qiagen) with on-column DNase I digestion and assessed with a Bioanalyzer for quality and Qubit for quantity. Multiplexed RNA sequencing was performed at the Monash Health Translation Precinct Medical Genomics Facility. Sample-specific indexes were added during the polyA priming step, before the pooled samples were amplified using template switching oligo. The samples were profiled with paired-end (19 bp forward, 72 bp reverse) sequencing using an Illumina NextSeq550 in high output mode with v2.5 chemistry. The raw paired-end barcoded multiplexed reads were initially demultiplexed using the Sabre v1.0 tool. Next, the reads were assessed for quality using FastQC v0.11.6, and low-quality bases were trimmed with Cutadapt v2.1. The trimmed reads were then aligned to the reference genome (hg38) using the STAR aligner v2.7.5b, and a count matrix was generated with the HTSeq v0.11.2 package. Finally, the processed count matrix for CAF and NPF samples was converted to logCPM values using the edgeR package in R v4.2.0.

### Analysis of morphomechanical features and matched transcriptomic data

For each morphological (nucleus area, nucleus circularity, width of F-actin angle distribution, cell volume) and mechanical parameter (apparent Young’s modulus measured by RT-DC), we calculated the ratio of the median values for CAFs versus NPF for each patient. Dimensionality reduction by principal component analysis was conducted in R (v4.2.0) with the prcomp function and the PC1 values were used as the morphomechanical score for subsequent analyses with RNA sequencing data. From a principal component analysis, the top 5000 highly variable genes across 35 pairs of CAF and NPF samples were identified. For these 5000 genes, we calculated the ratio of logCPM (CAF/NPF) values for each patient. Spearman’s rank correlation test used to examine the association of gene ratios with the morphomechanical score with *P* values adjusted for the false discovery rate (<0.05) using the p.adjust method in R v4.2.0. A heatmap of the gene expression profile of the 49 genes passing this cut-off was generated using the pheatmap function in R. The overlap of genes correlated with individual morphomechanical features and the morphomechanical score as depicted as an UpSet plot using UpSetR package in R^69^. For gene-set enrichment analyses, we ranked all genes in each sample based on their correlation with the morphomechanical score. We used the fgsea package in R for gene-set enrichment analysis of KEGG and gene ontology (GO) cellular component (CC) and biological process signatures. A subset of GO-CC enriched terms (size >10) were depicted as dot plot using the ggplot2 function in R. To compare the morphomechanical features to fibroblast phenotypes, we first determined the relative enrichment score for previously reported iCAF and myCAF signatures^70^ for each patient based on gene ratios in CAFs versus NPFs. Next, we used the Spearman’s rank correlation test to compare the enrichment scores to each morphological and mechanical features FDR (<0.05).

### Cell treatment with inhibitors (and activators)

For RT-DC, 100 000 cells were seeded into 6-well plates at least 24 h before treatment; 96 h before measurement. Medium was exchanged every 48 h. PDGF-BB and TGF-β1 were from Miltenyi Biotec. All inhibitors were sourced from Selleckchem. Inhibitors were added to medium and assay buffer during RT-DC measurement (table 1). For staining and morphological assessments, 70 000 cells were seeded onto Thermanox coverslips (Ø 13 mm, ThermoFisher) in 24-well plates at least 24 h before treatment, 96 h before fixation. For AFM measurements after treatment with TGF-β1 or TGF-β inhibitor (A83-01), 100 000 cells were seeded in 35 mm culture dishes (TPP) 24 h before treatment.

**Table 1.** Concentrations and treatment periods with clinical drugs, growth factors and small molecule inhibitors. Note, where indicated (*,**) stock solutions were in PBS/ H_2_O, respectively; medium and RT-DC assay buffer were supplemented with 1:1000 DMSO for consistency.

| Condition | Concentration | Duration |
| --- | --- | --- |
| Control | 1:1000 DMSO | 72 h |
| Axitinib | 1 μM | |
| PDGF-BB | 25 ng/mL * |  |
| Enzalutamide | 10 μM | |
| CCT251545 | 1 μM | |
| A83-01 | 5 μM | 48 h |
| TGF-β1 | 10 ng/mL * | |
| Simvastatin | 1 μM | 24 h |
| Docetaxel | 5 nM |  |
| MBQ-167 | 0.5 μM | 2 h |
| Nocodazole | 5 μM | 45 min |
| Docetaxel | 5 μM | |
| Y27632 | 10 μM ** | 30 min |

### Statistical analysis

Statistical analysis was performed with R (v4.2.0) and GraphPad Prism v10 software (GraphPad Software Inc.). Medians of measurements per patient and cell type were used. Normality of data was assessed with the Shapiro-Wilk test. Non-normally distributed data was tested for statistical significance using the Wilcoxon signed rank test for paired data and Mann-Whitney test for unpaired data.

With the R package lme4 ^71^, statistical significance between treatment groups and controls in the TGF-β and inhibitor experiments was determined by linear mixed effects analysis of RT-DC data. The treatment condition was chosen as fixed effect while the random effect was the by-patient random slope. CAFs and NPFs were tested separately and each treatment was tested against its control. Models with and without the fixed effect were constructed and a likelihood-ratio test was conducted to assess statistical significance^72,73^.

Morphomechanical parameter were analysed for correlation with the Spearman method after Shapiro-Wilk normality test. For comparisons of nuclear area and nuclear volume, linear correlation regression was performed in R with the lm function. Spearman correlation analysis was performed after testing for normal distribution with the Shapiro-Wilk test. Differences of survival curves in Kaplan-Meier plots were tested with the survdiff function of the survival R package (v3.5-7, https://cran.r-project.org/web/packages/survival/citation.html). P-values in graphs were indicated as [*, **, ***, ****] for p < 0.05, p < 0.01, p < 0.001, p < 0.0001 respectively. A p -value < 0.05 was considered statistically significant. An overview of all statistical test results is shown in Supplementary Table 4.

### Data availability

The RNA-sequencing data generated are available upon request via the database of Genotypes and Phenotypes (dbGaP) at https://www.ncbi.nlm.nih.gov/gap/ (study identification number phs003369.v3.p1).

## Acknowledgements

We acknowledge the members of the Prostate Cancer Research program, the patients, families and consumers who support our research, and the members of the Melbourne Urological Research Alliance (MURAL). We thank Jenna Kraska and Melissa Papargiris for patient recruitment and clinical follow up; Ana Sofia Rocha da Silva, Ana Ines Ferreira, Ana Margarita Gomez, Vinita Kini for assisting with image analysis; and Trevor Wilson for multiplex RNA sequencing (MHTP Medical Genomics Facility). We also thank Lauren Ryan, Ruth Pidsley and Susan Clark (Garvan Institute of Medical Research) for their helpful advice. This work was supported by the Monash University Faculty of Medicine, Nursing and Health Sciences (Platform Access Grant); Department of Health and Human Services acting through the Victorian Cancer Agency (MCRF18017); the Peter and Lyndy White Foundation; the Rotary Club of Manningham; TissuPath Pathology; the Australia–Germany Joint Research Co-operation Scheme (Universities Australia and DAAD; 57654961); support by the Mildred Scheel Nachwuchszentrum (MSNZ), funded by the German Cancer Aid; DFG (TA-751/4-1); Cancer Council New South Wales (RG 24-01; RP, ML), National Health and Medical Research Council (Investigator Grant #2010156 to RP); a University of New South Wales Postgraduate Award (LR) and the European Union (Erasmus+ student mobility). We also acknowledge technical support by Zellmechanik Dresden and the Bruker BioAFM team. The contents of the published material are solely the responsibility of the administering institution and individual authors and do not reflect the views of the NHMRC.

