## Supplementary Figures and Tables for "Prostate cancer associated fibroblasts have distinct morphomechanical features that are associated with patient outcome"

### **Supplementary material**

| **Feature** | **Summary** |
| --- | --- |
| Patients, number | 35 |
| Age, median (range) | 63.5 (42-76) |
| **Grade Group^a^, number (%)** | |
| GG1 | 0 (0%) |
| GG2 | 14 (40%) |
| GG3 | 10 (29%) |
| GG4 | 1 (3%) |
| GG5 | 9 (26%) |
| **Clinical features, media (range)** | |
| PSA (ng/ml) | 6.6 (2.4-40) |
| Tumour volume (cc) | 4.4 (0.7-30.2) |
| Clinical features, number (%) | |
| Pathologic T stage 2 | 11 (31%) |
| Pathologic T stage 3 | 24 (69%) |
| IDC-P reported | 11 (31%) |
| Positive margins | 8 (23%) |
| Extra-prostatic extension | 24 (69%) |
| Seminal vesicle invasion | 11 (31%) |
| Lymph node metastasis at diagnosis | 6 (17%) |
| **D’Amico risk score^b^, number (%)** | |
| Low risk (0) | 0 |
| Intermediate risk (1-3) | 0 |
| High risk (4-12) | 32 (94%) |
| **Patient follow-up, number (%)** | |
| Biochemical recurrence | 8 (23%) |
| Clinical recurrence | 7 (20%) |
| Any recurrence | 10 (29%) |

**Supplementary Table 1**. **Summary of clinical features and follow-up of the patient cohort.** ^a^The grade group was not determined for one patient who had prior androgen deprivation therapy. ^b^D’Amico scores could not be calculated for three patients.

| Patient Number | Age range at surgery | PSA (ng/mL) | Site of malignant sample | Site of benign sample | Tumour Volume (cc) | Primary Gleason Index | Secondary Gleason Index | Tertiary Gleason Index | Gleason Score | Grade group | D'Amico score | Tumour grade | Focality | IDCP present | Lymph involvement | Extraprostatic extension | Seminal vesicle invasion | Positive margin | Biochemical relapse | Clinical relapse | Any Relapse | Passage at measurement (CAF) | Passage at measurement (NPF) |
| --- | --- | --- | --- | --- | --- | --- | --- | --- | --- | --- | --- | --- | --- | --- | --- | --- | --- | --- | --- | --- | --- | --- | --- |
| 64R | 60s | 4.3 | Left Mid PZ | Right Mid TZ | 0.7 | 4 | 4 | n/a | 8 | 4 | 5 | pT2a | Unifocal | n/a | No | No | No | No | No | No | No | P8 | P8 |
| 76R | 60s | 9 | Left Mid to Apex PZ | Right TZ | 4.4 | 3 | 4 | n/a | 7 | 2 | 6 | T2c | Multifocal | n/a | No | No | No | No | Yes | No | Yes | P7 | P5 |
| 105R | 50s | 5.1 | Right Mid PZ | Left TZ | 2 | 4 | 3 | n/a | 7 | 3 | 6 | pT3a | Unifocal | n/a | n/a | Yes | No | No | No | No | No | P8 | P8 |
| 128R | 50s | 6 | Left peripheral zone | Right transition zone | 1.8 | 3 | 4 | n/a | 7 | 2 | 6 | pT3b | Unifocal | n/a | No | Yes | Yes | Yes | No | No | No | P8 | P9 |
| 140R | 60s | 6.7 | Right mid peripheral zone | Left transition zone | 7.1 | 4 | 3 | n/a | 7 | 3 | 6 | pT3b | Multifocal | No | No | Yes | Yes | No | No | No | No | P8 | P8 |
| 153R | 60s | 40 | Left PZ | Right TZ | 1.8 | 4 | 5 | n/a | 9 | 5 | 9 | pT3bN1 | Multifocal | No | Yes | Yes | Yes | Yes | Yes | Yes | Yes | P5 | P5 |
| 156R | 70s | 16.8 | Right superior and right inferior | TZ | 28.6 | 4 | 5 | n/a | 9 | 5 | 8 | pT3b | Multifocal | No | No | Yes | Yes | Yes | Yes | No | Yes | P3 | P4 |
| 164R | 60s | 6.2 | Left base PZ and right base PZ | Left mid TZ | 11.2 | 4 | 5 | n/a | 9 | 5 | 7 | pT3bN1 | Multifocal | n/a | Yes | Yes | Yes | Yes | Yes | Yes | Yes | P3 | P3 |
| 166R | 60s | 32.5 | Right base | Right TZ | 4.3 | n/a | n/a | n/a | n/a | n/a | n/a | pT3aN1 | Unifocal | No | Yes | Yes | No | No | Yes | Yes | Yes | P5 | P5 |
| 186R | 60s | 8.1 | Right PZ | Left TZ | 4.8 | 3 | 4 | n/a | 7 | 2 | 4 | pT2a | Unifocal | No | n/a | No | No | No | No | No | No | P3 | P3 |
| 189R | 60s | 6.5 | Left mid PZ | Right mid TZ | 6.3 | 4 | 3 | n/a | 7 | 3 | 6 | pT3a | Multifocal | No | n/a | Yes | No | No | No | No | No | P3 | P3 |
| 190R | 60s | n/a | Left mid TZ lateral | Right TZ | 5.2 | 4 | 3 | n/a | 7 | 3 | n/a | pT3a | Multifocal | Yes | n/a | Yes | No | No | No | No | No | P4 | P2 |
| 196R | 70s | 8.9 | Post mid line | Right TZ | 1.4 | 4 | 3 | n/a | 7 | 3 | 6 | pT3bN0 | Multifocal | No | No | Yes | Yes | No | Yes | No | Yes | P5 | P3 |
| 198R | 70s | 10.82 | Left mid posterior | Right TZ | 3.7 | 3 | 4 | n/a | 7 | 2 | 7 | pT3bN0 | Multifocal | No | No | Yes | Yes | No | No | No | No | P4 | P5 |
| 204R | 60s | 2.4 | Left TZ | Right anterior | 3.2 | 3 | 4 | n/a | 7 | 2 | 6 | pT3aN0 | Unifocal | No | No | Yes | No | Yes | No | No | No | P4 | P2 |
| 210R | 40s | 2.7 | Left base | Right mid | 2.1 | 3 | 4 | n/a | 7 | 2 | 6 | pT2c | Multifocal | No | n/a | No | No | No | No | No | No | P3 | P3 |
| 213R | 60s | 5.8 | Right PZ mid and base | Left TZ mid | 3.6 | 3 | 4 | n/a | 7 | 2 | 6 | pT3a | mulitfocal | No | n/a | Yes | No | No | No | No | No | P3 | P4 |
| 218R | 60s | 10 | Right posterior | Left TZ | 1.6 | 3 | 4 | n/a | 7 | 2 | 6 | pT2b | Multifocal | No | No | No | No | No | No | No | No | P5 | P4 |
| 224R | 60s | 5.1 | Left PZ | Right TZ | 8 | 4 | 5 | n/a | 9 | 5 | 7 | pT3b | Multifocal | Yes | No | Yes | Yes | Yes | No | Yes | Yes | P3 | P3 |
| 225R | 50s | 18 | Right mid PZ | Left TZ | 4.6 | 3 | 4 | n/a | 7 | 2 | 7 | pT3a | Multifocal | No | No | Yes | No | No | No | No | No | P2 | P2 |
| 228R | 60s | 4.7 | Midline posterior and right mid PZ | Right TZ | 7.2 | 3 | 4 | n/a | 7 | 2 | 6 | pT2c | Multifocal | Yes | No | No | No | No | No | No | No | P5 | P3 |
| 229R | 60s | 7 | Left mid | Right TZ | 19.7 | 4 | 5 | n/a | 9 | 5 | 7 | pT3bN1 | Unifcoal | Yes | Yes | Yes | Yes | Yes | Yes | Yes | Yes | P4 | P4 |
| 262R | 70s | 9.9 | Left post mid | Right TZ | 4.4 | 4 | 3 | 5 | 7 | 3 | 6 | pT2cN0 | Multifocal | Yes | Yes | No | No | No | No | No | No | P4 | P4 |
| 265R | 50s | 4.2 | Left mid | Right TZ | 2.4 | 3 | 4 | n/a | 7 | 2 | 6 | pT3a | Multifocal | No | n/a | Yes | No | No | No | No | No | P3 | P3 |
| 266R | 70s | 5.3 | Right mid PZ | Left TZ | 3.2 | 4 | 3 | n/a | 7 | 3 | 6 | pT3aN0 | Multifocal | Yes | No | Yes | No | No | No | No | No | P3 | P3 |
| 270R | 60s | 3.2 | Right PZ mid | Left TZ mid | 2.2 | 4 | 3 | n/a | 7 | 3 | 6 | pT3a | Multifocal | No | n/a | Yes | No | No | No | No | No | P6 | P4 |
| 289R | 60s | 4.8 | Right mid posterior | Left mid posterior | 0.8 | 4 | 3 | n/a | 7 | 3 | 4 | pT2aN0 | Unifocal | No | No | No | No | No | No | No | No | P4 | P3 |
| 294R | 60s | 5.6 | left mid PZ | Right mid PZ | 30.2 | 5 | 4 | n/a | 9 | 5 | 7 | PT3bN0 | Unifocal | Yes | No | Yes | Yes | No | Yes | Yes | Yes | P4 | P2 |
| 307R | 70s | 8 | Left posterior | Right TZ | 6.1 | 4 | 3 | 5 | 7 | 3 | 6 | pT2cN0 | Multifocal | Yes | No | No | No | No | No | No | No | P3 | P2 |
| 332R | 60s | 11 | Left post | Right mid post | 5.4 | 4 | 5 | n/a | 9 | 5 | 8 | pT3a | Multifocal | Yes | No | Yes | No | Yes | No | No | No | P3 | P3 |
| 338R | 40s | 8.1 | Right mid post | Left TZ | 13 | 3 | 4 | n/a | 7 | 2 | 6 | pT3a | Multifocal | No | n/a | Yes | No | No | No | No | No | P3 | P3 |
| 365R | 60s | 6.3 | Left mid PZ | Right TZ | 8.6 | 4 | 5 | n/a | 9 | 5 | 7 | pT3aN1 | Unifocal | Yes | Yes | Yes | No | No | No | Yes | Yes | P4 | P3 |
| 397R | 50s | 6 | Left Apex | Right mid | 0.9 | 3 | 4 | n/a | 7 | 2 | 6 | pT2c | Multifocal | No | n/a | No | No | No | No | No | No | P3 | P3 |
| 405R | 70s | 9.2 | Left Posterior Mid | Right TZ | 10.3 | 4 | 5 | n/a | 9 | 5 | 7 | PT3b NO | Multifocal | Yes | No | Yes | Yes | No | No | No | No | P3 | P3 |
| 441R | 70s | 9.4 | Left PZ | Right TZ | 3.6 | 3 | 4 | n/a | 7 | 2 | n/a | PT2 | Multifocal | No | n/a | No | No | No | No | No | No | P3 | P4 |

**Supplementary Table 2: Clinical features and follow up of the patient cohort with passage number of cells at measurement**

| Patient | Celltype | Number of replicates RT-DC | Passages measured | Patient | Celltype | Number of replicates RT-DC | Passages measured | Patient | Celltype | Number of replicates RT-DC | Passages measured |
| --- | --- | --- | --- | --- | --- | --- | --- | --- | --- | --- | --- |
| 64 | CAF | 2 | P8, P10 | 196 | CAF | 3 | P5, P7, P10 | 266 | CAF | 2 | P3, P4 |
| 64 | NPF | 2 | P8, P10 | 196 | NPF | 3 | P3, P5, P8 | 266 | NPF | 2 | P3, P4 |
| 76 | CAF | 3 | P7, P9, P12 | 198 | CAF | 2 | P4, P7 | 270 | CAF | 1 | P6 |
| 76 | NPF | 3 | P5, P7, P10 | 198 | NPF | 2 | P5, P8 | 270 | NPF | 1 | P4 |
| 105 | CAF | 1 | P8 | 204 | CAF | 2 | P4, P6 | 289 | CAF | 1 | P4 |
| 105 | NPF | 1 | P8 | 204 | NPF | 2 | P2, P4 | 289 | NPF | 1 | P3 |
| 128 | CAF | 1 | P8 | 210 | CAF | 2 | P3, P5 | 294 | CAF | 2 | P4, P6 |
| 128 | NPF | 1 | P9 | 210 | NPF | 2 | P3, P5 | 294 | NPF | 2 | P2, P4 |
| 140 | CAF | 2 | P8, P10 | 213 | CAF | 2 | P3, P5 | 307 | CAF | 2 | P3, P4 |
| 140 | NPF | 2 | P8, P10 | 213 | NPF | 2 | P4, P6 | 307 | NPF | 2 | P2, P3 |
| 153 | CAF | 1 | P5 | 218 | CAF | 1 | P5 | 332 | CAF | 2 | P3, P7 |
| 153 | NPF | 1 | P5 | 218 | NPF | 1 | P4 | 332 | NPF | 2 | P3, P7 |
| 156 | CAF | 1 | P3 | 224 | CAF | 2 | P3, P5 | 338 | CAF | 2 | P3, P4 |
| 156 | NPF | 1 | P4 | 224 | NPF | 2 | P3, P5 | 338 | NPF | 2 | P3, P4 |
| 164 | CAF | 5 | P3, P4, P5, P7, P10 | 225 | CAF | 2 | P2, P4 | 365 | CAF | 1 | P4 |
| 164 | NPF | 5 | P3, P4, P5, P7, P10 | 225 | NPF | 2 | P2, P4 | 365 | NPF | 1 | P3 |
| 166 | CAF | 1 | P5 | 228 | CAF | 1 | P5 | 397 | CAF | 2 | P3, P5 |
| 166 | NPF | 1 | P5 | 228 | NPF | 1 | P3 | 397 | NPF | 2 | P3, P5 |
| 186 | CAF | 2 | P3, P4 | 229 | CAF | 4 | P4, P5, P7, P10 | 405 | CAF | 2 | P3, P5 |
| 186 | NPF | 2 | P3, P4 | 229 | NPF | 4 | P4, P5, P7, P10 | 405 | NPF | 2 | P3, P5 |
| 189 | CAF | 1 | P3 | 262 | CAF | 1 | P4 | 441 | CAF | 2 | P3, P6 |
| 189 | NPF | 1 | P3 | 262 | NPF | 1 | P4 | 441 | NPF | 2 | P4, P6 |
| 190 | CAF | 1 | P4 | 265 | CAF | 1 | P3 |  |  |  |  |
| 190 | NPF | 1 | P2 | 265 | NPF | 1 | P3 |  |  |  |  |

**Supplementary Table 3.** Overview table of the repeat measurements over various passages.

| Figure | Group | *P*-value | Test | n |
| --- | --- | --- | --- | --- |
| 2 f - F-actin angles Gaussian width | CAF vs NPF | 0.0034 | Wilcoxon Signed-Rank Test (paired) | 35 (pairs) |
| 2 i - Nuclear area | CAF vs NPF | 0.00016 | Wilcoxon Signed-Rank Test (paired) | 35 (pairs) |
| 2 l - Nuclear circularity | CAF vs NPF | 5.60E-05 | Wilcoxon Signed-Rank Test (paired) | 35 (pairs) |
| 3 c - Apparent Young's modulus RT-DC | CAF vs NPF | 0.00017 | Wilcoxon Signed-Rank Test (paired) | 35 (pairs) |
| 3 d - Apparent Young's modulus AFM | CAF vs NPF | 0.004 | Wilcoxon Signed-Rank Test (paired) | 14 (pairs) |
| 3 e - Cell volume RT-DC | CAF vs NPF | 0.00017 | Wilcoxon Signed-Rank Test (paired) | 35 (pairs) |
| 4 b - PC1 vs GG | CAF vs NPF (2&3) | 0.00085 | Wilcoxon Signed-Rank Test (paired) | 24 (pairs) |
|  | CAF vs NPF (4&5) | 0.01367 |  | 10 (pairs) |
| 6 c - Nuclear Area | CAF - A83-01 vs ctrl | 0.377 | Likelihood ratio test of linear mixed effects model | 282 vs 229 |
|  | NPF -A83-01 vs ctrl | 0.222 |  | 311 vs 261 |
|  | CAF - TGF-β1 vs ctrl | 0.002 |  | 171 vs 229 |
|  | NPF - TGF-β1 vs ctrl | 0.051 |  | 211 vs 261 |
| 6 d - Nuclear circularity | CAF - A83-01 vs ctrl | 0.473 | Likelihood ratio test of linear mixed effects model | 282 vs 229 |
|  | NPF -A83-01 vs ctrl | 0.478 |  | 311 vs 261 |
|  | CAF - TGF-β1 vs ctrl | 0.77 |  | 171 vs 229 |
|  | NPF - TGF-β1 vs ctrl | 0.678 |  | 211 vs 261 |
| 6 e - F-actin angle Gaussian width | CAF - A83-01 vs ctrl | 0.408196 | Likelihood ratio test of linear mixed effects model | 13 vs 14 |
|  | NPF -A83-01 vs ctrl | 0.886074 |  | 14 vs 12 |
|  | CAF - TGF-β1 vs ctrl | 0.08484 |  | 13 vs 14 |
|  | NPF - TGF-β1 vs ctrl | 0.563148 |  | 13 vs 12 |
| 6 f - Apparent Young's modulus AFM | CAF - A83-01 vs ctrl | 0.039767 | Likelihood ratio test of linear mixed effects model | 202 vs 200 |
|  | NPF -A83-01 vs ctrl | 0.035664 |  | 202 vs 200 |
|  | CAF - TGF-β1 vs ctrl | 0.116367 |  | 203 vs 200 |
|  | NPF - TGF-β1 vs ctrl | 0.00247 |  | 195 vs 200 |
| 6 h - Apparent Young's modulus RT-DC | CAF - A83-01 vs ctrl | 0.000716 | Likelihood ratio test of linear mixed effects model | 64325 vs 51366 |
|  | NPF -A83-01 vs ctrl | 0.06439 |  | 56546 vs 59611 |
|  | CAF - TGF-β1 vs ctrl | 0.01041 |  | 48917 vs 51366 |
|  | NPF - TGF-β1 vs ctrl | 0.02754 |  | 53944 vs 59611 |
| 6 i - Cell volume RT-DC | CAF - A83-01 vs ctrl | 0.008689 | Likelihood ratio test of linear mixed effects model | 64325 vs 51366 |
|  | NPF -A83-01 vs ctrl | 0.05481 |  | 56546 vs 59611 |
|  | CAF - TGF-β1 vs ctrl | 0.000979 |  | 48917 vs 51366 |
|  | NPF - TGF-β1 vs ctrl | 0.01546 |  | 53944 vs 59611 |
| 7 b - Nuclear area | CAF - Axi vs ctrl | 0.003 | Likelihood ratio test of linear mixed effects model | 316 vs 420 |
|  | NPF - Axi vs ctrl | 0.009 |  | 432 vs 487 |
|  | CAF - Doce vs ctrl | 0.431 |  | 155 vs 420 |
|  | NPF - Doce vs ctrl | 0.201 |  | 284 vs 487 |
|  | CAF - Enz vs ctrl | 0.364 |  | 442 vs 420 |
|  | NPF - Enz vs ctrl | 0.591 |  | 572 vs 487 |
|  | CAF - Sim vs ctrl | 0.086 |  | 254 vs 420 |
|  | NPF - Sim vs ctrl | 0.001 |  | 379 vs 487 |
|  | CAF - CCT vs ctrl | 0.377 |  | 366 vs 420 |
|  | NPF - CCT vs ctrl | 0.053 |  | 400 vs 487 |
|  | CAF - PDGF vs ctrl | 0.031 |  | 431 vs 420 |
|  | NPF - PDGF vs ctrl | 0.828 |  | 469 vs 487 |
| 7 c - Nuclear circularity | CAF - Axi vs ctrl | 0.003 | Likelihood ratio test of linear mixed effects model | 316 vs 420 |
|  | NPF - Axi vs ctrl | 0.723 |  | 432 vs 487 |
|  | CAF - Doce vs ctrl | 0.002 |  | 155 vs 420 |
|  | NPF - Doce vs ctrl | 0.154 |  | 284 vs 487 |
|  | CAF - Enz vs ctrl | 0.088 |  | 442 vs 420 |
|  | NPF - Enz vs ctrl | 0.967 |  | 572 vs 487 |
|  | CAF - Sim vs ctrl | 0.423 |  | 254 vs 420 |
|  | NPF - Sim vs ctrl | 0.401 |  | 379 vs 487 |
|  | CAF - CCT vs ctrl | 0.011 |  | 366 vs 420 |
|  | NPF - CCT vs ctrl | 0.099 |  | 400 vs 487 |
|  | CAF - PDGF vs ctrl | 0.087 |  | 431 vs 420 |
|  | NPF - PDGF vs ctrl | 0.393 |  | 469 vs 487 |
| 7 e - Apparent Young's modulus | CAF - Axi vs ctrl | 0.003328 | Likelihood ratio test of linear mixed effects model | 23518 vs 45909 |
|  | NPF - Axi vs ctrl | 0.005438 |  | 31771 vs 49314 |
|  | CAF - Doce vs ctrl | 0.002123 |  | 29363 vs 29624 |
|  | NPF - Doce vs ctrl | 0.006711 |  | 23749 vs 25707 |
|  | CAF - Enz vs ctrl | 0.1439 |  | 50522 vs 45692 |
|  | NPF - Enz vs ctrl | 0.3086 |  | 44332 vs 49665 |
|  | CAF - Sim vs ctrl | 0.1414 |  | 35688 vs 35942 |
|  | NPF - Sim vs ctrl | 0.1632 |  | 37884 vs 39352 |
|  | CAF - CCT vs ctrl | 0.1741 |  | 28553 vs 28797 |
|  | NPF - CCT vs ctrl | 0.1479 |  | 33222 vs 30892 |
|  | CAF - PDGF vs ctrl | 0.05177 |  | 35084 vs 35554 |
|  | NPF - PDGF vs ctrl | 0.3335 |  | 33627 vs 32347 |
|  | CAF - MBQ vs ctrl | 0.1923 |  | 45711 vs 42765 |
|  | NPF - MBQ vs ctrl | 0.0363 |  | 36794 vs 46223 |
|  | CAF - Noco vs ctrl | 0.004724 |  | 39081 vs 39374 |
|  | NPF - Noco vs ctrl | 0.001052 |  | 38421 vs 34919 |
|  | CAF - Y27 vs ctrl | 0.9802 |  | 93408 vs 97712 |
|  | NPF - Y27 vs ctrl | 0.4079 |  | 103127 vs 90777 |
| 7 f - Cell volume | CAF - Axi vs ctrl | 0.000471 | Likelihood ratio test of linear mixed effects model | 24215 vs 46255 |
|  | NPF - Axi vs ctrl | 0.002057 |  | 32158 vs 49638 |
|  | CAF - Doce vs ctrl | 0.4749 |  | 29969 vs 29981 |
|  | NPF - Doce vs ctrl | 0.2515 |  | 24835 vs 26048 |
|  | CAF - Enz vs ctrl | 0.9929 |  | 51140 vs 46253 |
|  | NPF - Enz vs ctrl | 0.4034 |  | 43867 vs 49178 |
|  | CAF - Sim vs ctrl | 0.4558 |  | 36370 vs 36249 |
|  | NPF - Sim vs ctrl | 0.5 |  | 38809 vs 39636 |
|  | CAF - CCT vs ctrl | 0.01877 |  | 28835 vs 29078 |
|  | NPF - CCT vs ctrl | 0.02106 |  | 33611 vs 31162 |
|  | CAF - PDGF vs ctrl | 0.4026 |  | 35925 vs 35808 |
|  | NPF - PDGF vs ctrl | 0.00628 |  | 34397 vs 32636 |
|  | CAF - MBQ vs ctrl | 0.3544 |  | 46292 vs 43078 |
|  | NPF - MBQ vs ctrl | 0.6052 |  | 37443 vs 46548 |
|  | CAF - Noco vs ctrl | 0.101 |  | 39512 vs 39588 |
|  | NPF - Noco vs ctrl | 0.1723 |  | 39375 vs 35226 |
|  | CAF - Y27 vs ctrl | 0.1838 |  | 94057 vs 98360 |
|  | NPF - Y27 vs ctrl | 0.3876 |  | 103871 vs 91214 |

**Supplementary Table 4.** Results of statistical tests in main figures.

| #All genes with adjusted P value < 0.05 | | | |
| --- | --- | --- | --- |
| **Gene** | **Correlation** | **P value** | **Adjusted P value** |
| NAV3 | 0.63697479 | 6.28E-06 | 0.000505 |
| CTHRC1 | 0.59887955 | 0.000519 | 0.024771 |
| SLC14A1 | 0.59663866 | 0.000997 | 0.042958 |
| CDKN3 | 0.59439776 | 1.09E-05 | 0.000828 |
| TNFRSF12A | 0.59047619 | 0.000569 | 0.026781 |
| TK1 | 0.57871148 | 0.0001 | 0.006013 |
| PTTG1 | 0.56778711 | 1.70E-05 | 0.001241 |
| SGO2 | 0.55994398 | 1.61E-05 | 0.001182 |
| KNSTRN | 0.55014006 | 5.48E-05 | 0.003527 |
| IL7R | 0.54957983 | 0.000933 | 0.040636 |
| MT2A | 0.53501401 | 4.66E-05 | 0.003054 |
| SPDL1 | 0.52184874 | 0.000446 | 0.021766 |
| TOP2A | 0.48095238 | 1.44E-05 | 0.001069 |
| INSIG1 | 0.45378151 | 0.000192 | 0.010572 |
| DHCR24 | 0.43865546 | 0.000246 | 0.013091 |
| PRC1 | 0.41932773 | 0.000941 | 0.040927 |
| PTX3 | 0.41372549 | 0.000841 | 0.03726 |
| SERINC2 | 0.40644258 | 0.001043 | 0.044597 |
| HIST1H1A | 0.34985994 | 0.000871 | 0.038383 |
| CD160 | -0.2420338 | 7.87E-05 | 0.004853 |
| OR7E47P | -0.2887955 | 0.000582 | 0.027314 |
| RGN | -0.367507 | 0.000128 | 0.007424 |
| PTGDS | -0.372549 | 0.000723 | 0.032836 |
| PF4 | -0.4031095 | 0.000641 | 0.029644 |
| SPTSSA | -0.4383754 | 0.000306 | 0.015776 |
| BEAN1 | -0.4557052 | 0.00111 | 0.046946 |
| LINC01082 | -0.4596639 | 0.000593 | 0.027751 |
| EFNA1 | -0.4742297 | 0.000989 | 0.042646 |
| GPX3 | -0.4823529 | 0.000177 | 0.009867 |
| ZNF385D | -0.4927516 | 0.001135 | 0.047794 |
| GHR | -0.4960784 | 0.000522 | 0.024891 |
| RARRES2 | -0.4980392 | 5.21E-05 | 0.003371 |
| ARHGAP28 | -0.4980392 | 0.000704 | 0.03208 |
| MYOCD | -0.5120448 | 0.00026 | 0.013727 |
| RGS9 | -0.5148459 | 0.001085 | 0.046041 |
| S1PR1 | -0.519888 | 0.000847 | 0.037487 |
| ANGPTL4 | -0.5229692 | 0.000197 | 0.010812 |
| ERFE | -0.52493 | 0.000194 | 0.010644 |
| FOSB | -0.5492997 | 0.000114 | 0.006744 |
| RPL10P9 | -0.57507 | 0.000607 | 0.028308 |
| CFD | -0.5767507 | 2.96E-05 | 0.002036 |
| IGFBP5 | -0.5840336 | 0.00012 | 0.007023 |
| TXLNGY | -0.6005602 | 1.46E-05 | 0.001078 |
| MAPK10 | -0.6056022 | 0.000176 | 0.009793 |
| HES1 | -0.6313725 | 6.08E-05 | 0.003869 |
| C11orf96 | -0.637535 | 0.000455 | 0.022149 |
| AKR1C1 | -0.6498599 | 0.00052 | 0.024782 |
| CRISPLD2 | -0.6546218 | 3.50E-07 | 3.52E-05 |
| FOS | -0.6960784 | 1.20E-06 | 0.000111 |

**Supplementary Table 5.** Candidate genes that are significantly correlated with the morphomechanical score.

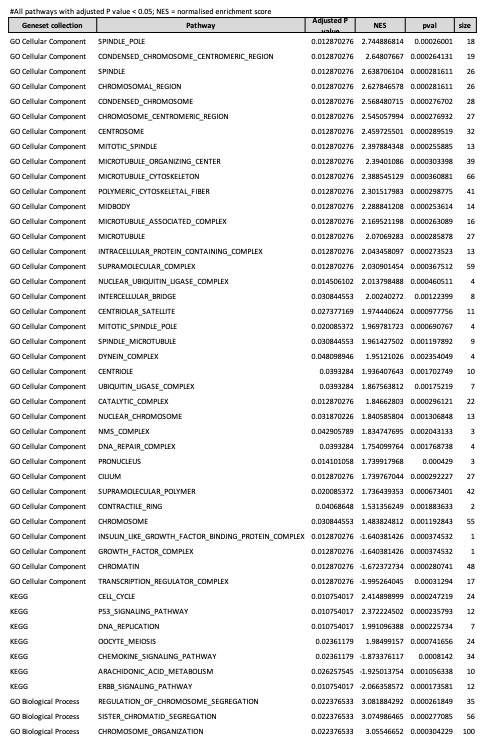

**Supplementary Table 6.** Geneset enrichment analysis based on the correlation of gene ratios with the morphomechanical score.

| **Feature 1** | **Feature 2** | **Correlation (r)** | **P value** |
| --- | --- | --- | --- |
| iCAF | myCAF | 0.07 | 0.633127 |
| iCAF | morphomechanics score | -0.13 | 0.308822 |
| iCAF | ratio_angle | 0.32 | 0.071517 |
| iCAF | ratio_nuc_area | -0.3 | 0.226878 |
| iCAF | ratio_nuc_circ | -0.17 | 0.901766 |
| iCAF | ratio_volume | 0.01 | 0.838264 |
| iCAF | ratio_young | -0.1 | 0.527328 |
| myCAF | morphomechanics score | 0.36 | 0.03837 |
| myCAF | ratio_angle | 0.14 | 0.582547 |
| myCAF | ratio_nuc_area | 0.3 | 0.157802 |
| myCAF | ratio_nuc_circ | -0.17 | 0.238442 |
| myCAF | ratio_volume | 0.25 | 0.144083 |
| myCAF | ratio_young | 0.43 | 0.004371 |

**Supplementary Table 7.** Correlation between CAF subtype signatures and morphological and biomechanical features.

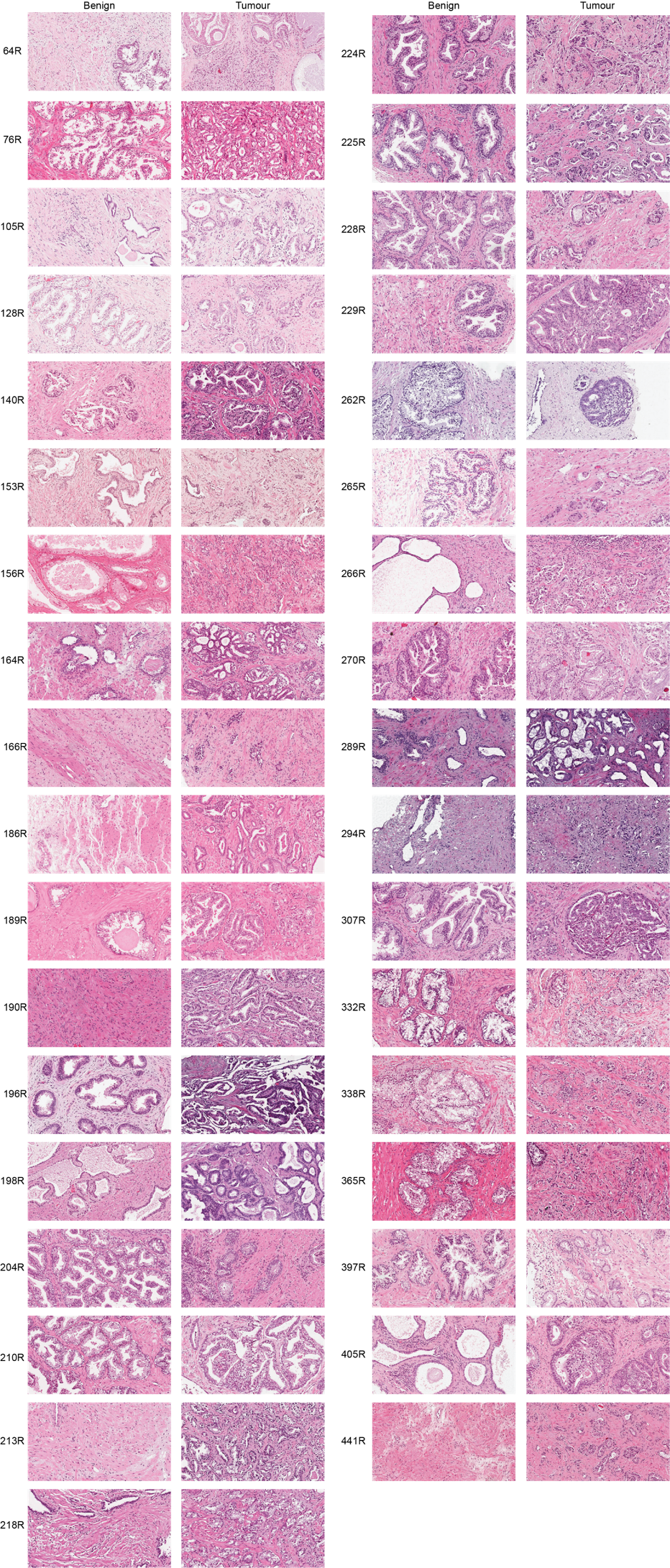
**Supplementary Figure 1. Summary of the pathology of each patient sample.** Representative images of haematoxylin and eosin stained tissue from the benign and tumour region for each patient. Small pieces of tissue were retained from each sample, while the remaining tissue was digested to establish primary cultures of CAFs and NPFs.

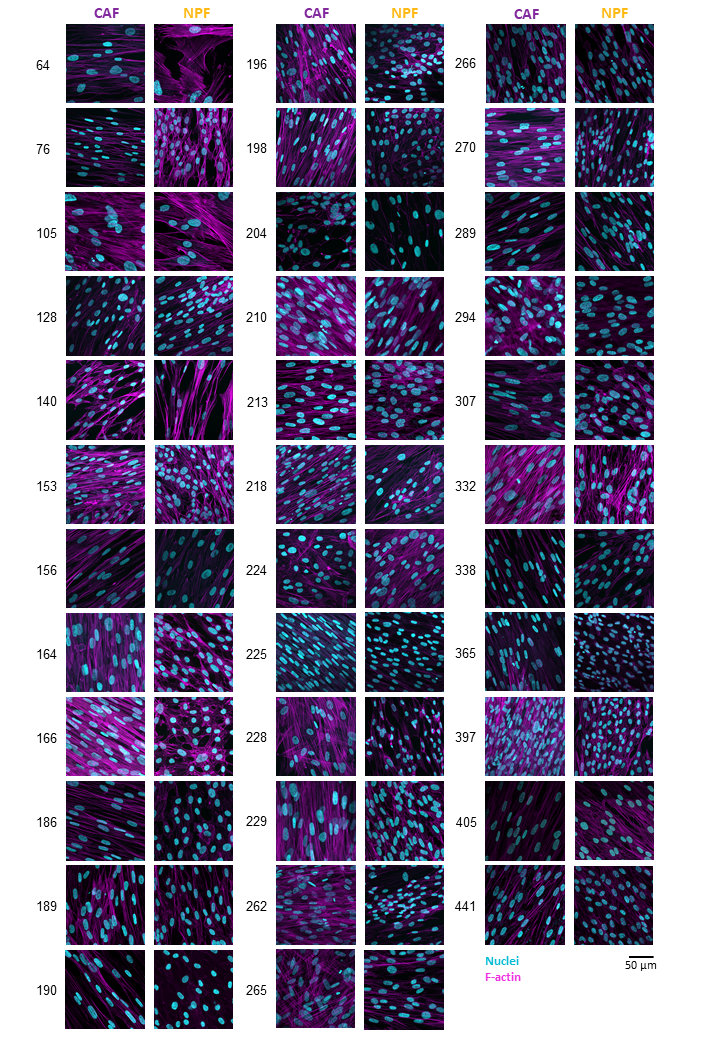

**Supplementary Figure 2. Overview of CAF and NPF morphology.** Representative images of CAFs and NPFs after 14 days in culture and stained with DAPI (DNA - cyan) and phalloidin-TRITC (F-actin - magenta). Images were recorded using a confocal microscope (Zeiss LSM780).

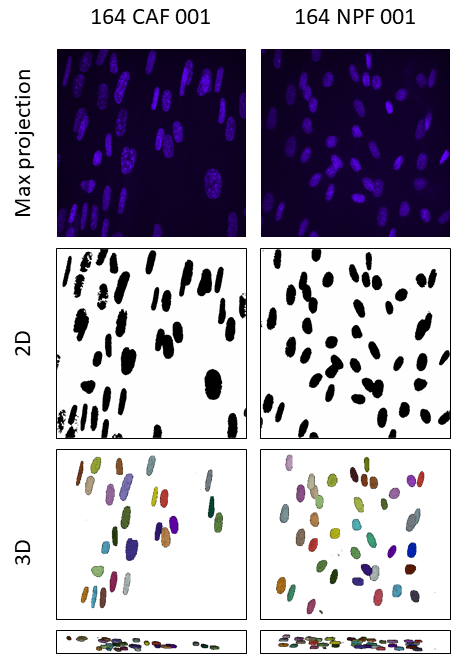

**Supplementary Figure 3. Segmentation of nuclei for morphometric analysis.** Example images for the analysis for comparison of nuclear projected area and volume. CAF/NPF cultures were stained for their nuclei (blue) using DAPI and imaged using a confocal microscope (top). 3D projected nuclei were segmented with Fiji (version 1.53o) (middle). 3D segmentation was conducted with Cellpose (version 2.2) and nuclei were viewed with Napari (version 0.4.17) (bottom).

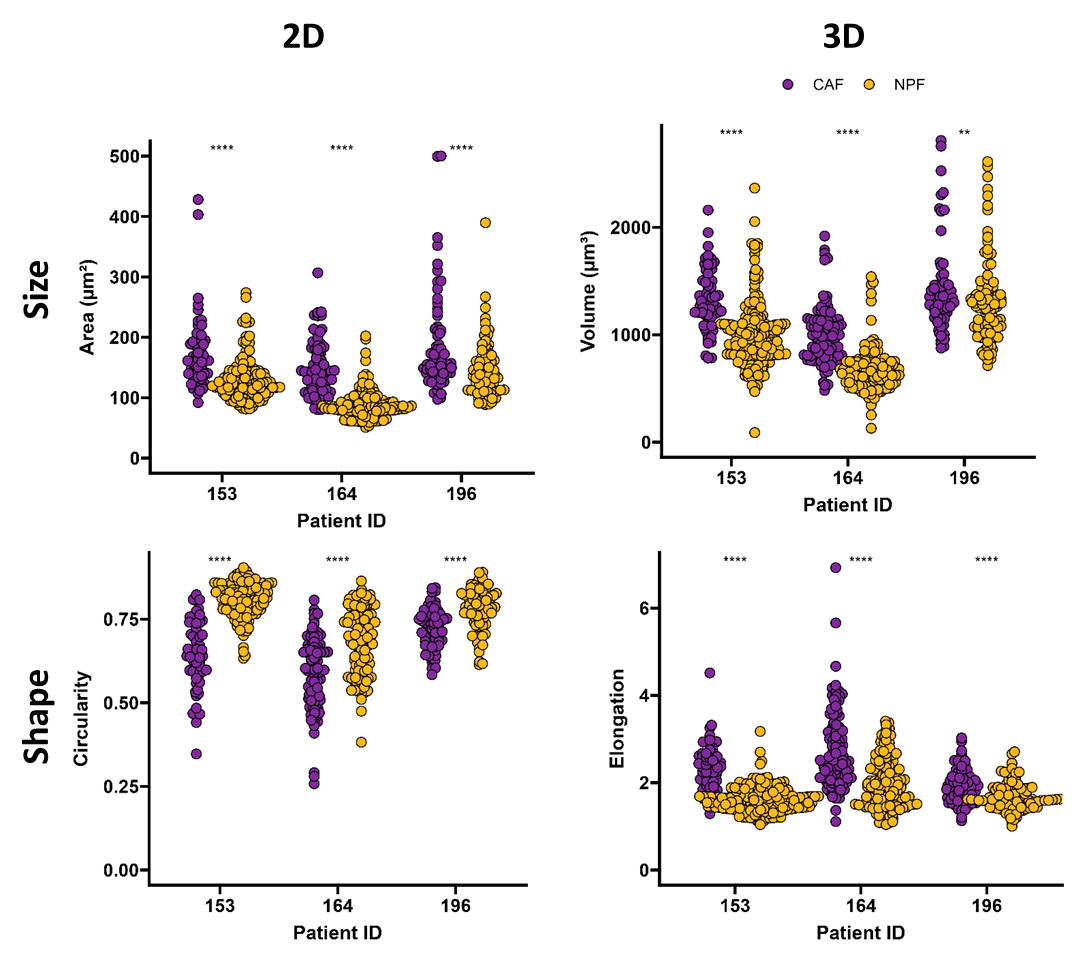

**Supplementary Figure 4. Comparison of CAFs (purple) and NPFs (orange) size and shape descriptors of the same images.** Three representative patients are shown. Each dot represents one nucleus. n_2D_ = 65 - 165, n_3D_ = 74 – 218. Results of a Mann-Whitney test shown (****: p<0.001, **: p<0.01)). 2D Area: 153: *P*=1.4∙10^-12^, 164: *P*< 2∙10^-16^, 196: *P*=1.6∙10^-5^; 2D Circularity: 153: *P*< 2∙10^-16^, 164: *P*=8.3∙10^-12^, 196: *P*=9.1∙10^-11^; 3D Volume: 153: *P*<2 ∙ 10^-16^, 164: *P*<2∙10^-16^, 196: *P*=0.0074; 3D Elongation: 153: *P*<2 ∙ 10^-16^, 164: *P*<2∙10^-16^, 196: *P*=3∙10^-6^.

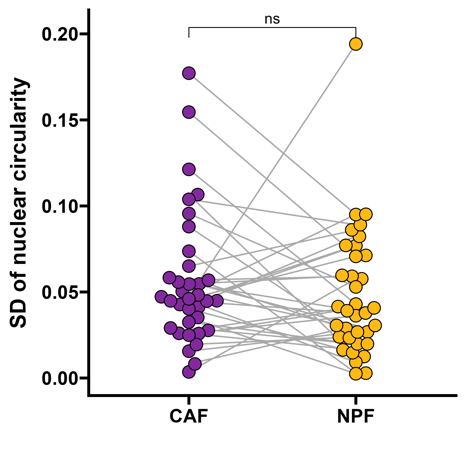

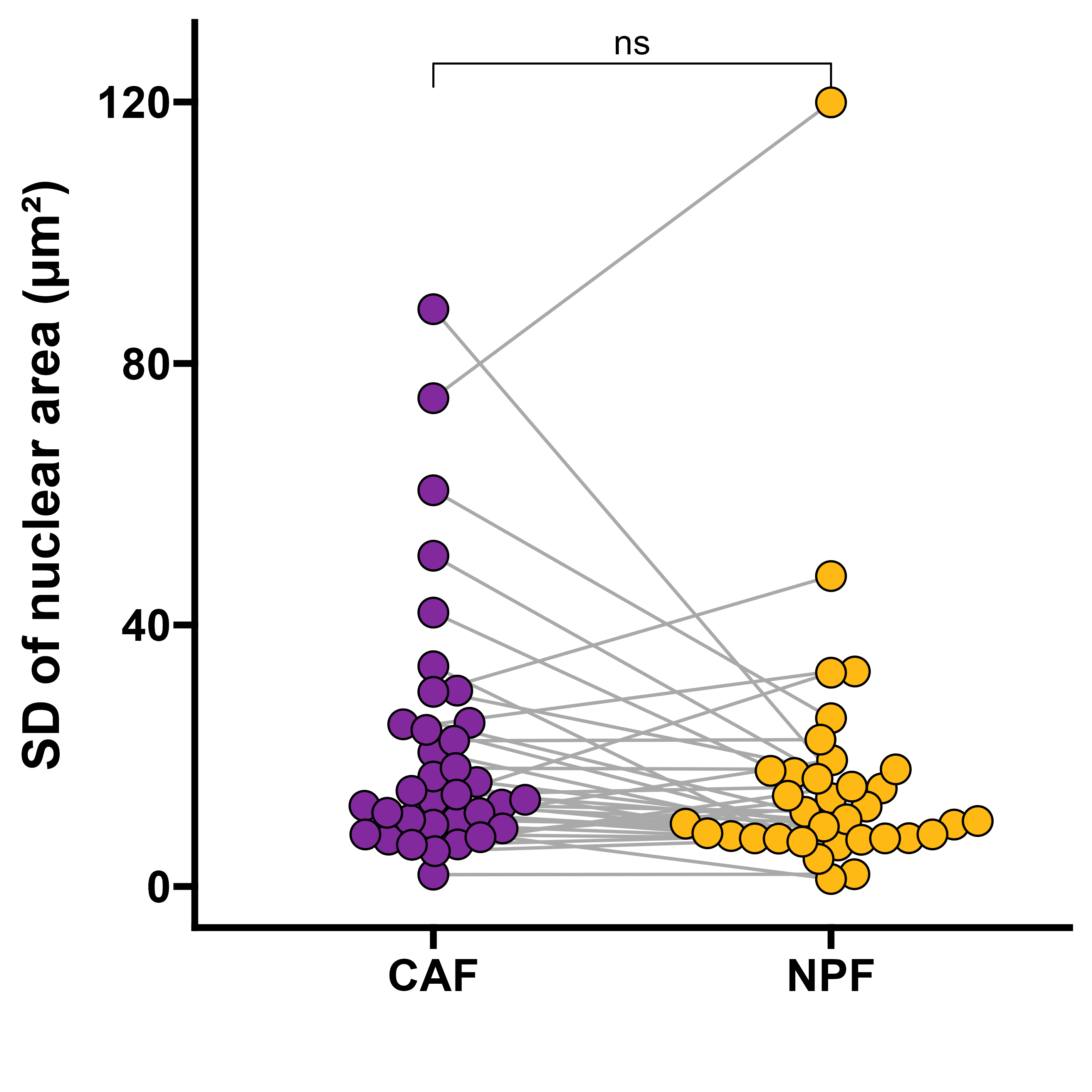

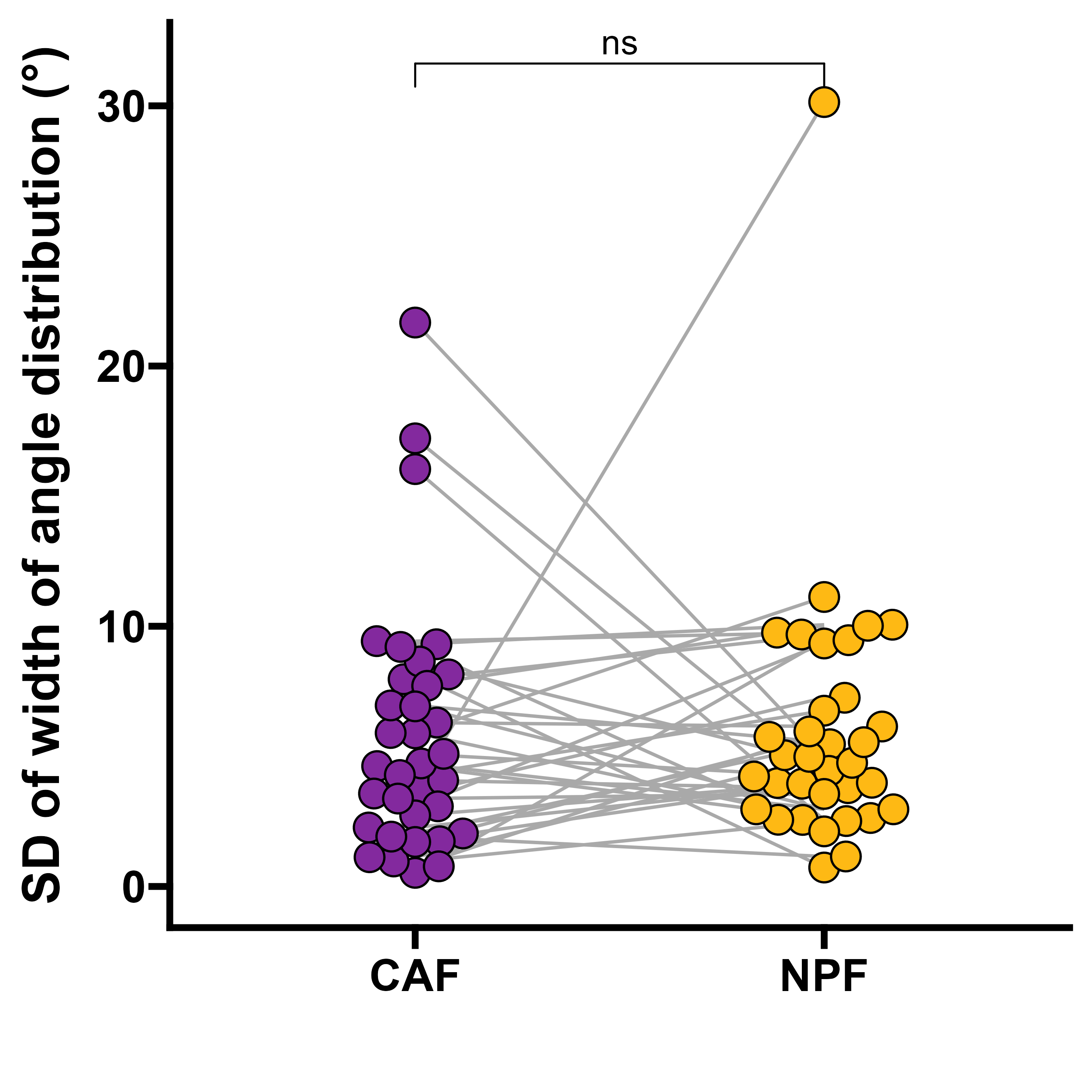

**Supplementary Figure 5. Standard deviations of morphological parameters.** Standard deviations of nuclear area, circularity and angle distribution were calculated per patient and cell type. Results of a Wilcoxon Signed-Rank test are shown (ns – non-significant, *P*(area) = 0.097; *P*(circ.) = 0.29; *P*(angle distr.) = 0.64). n=35 (donor pairs, medians).

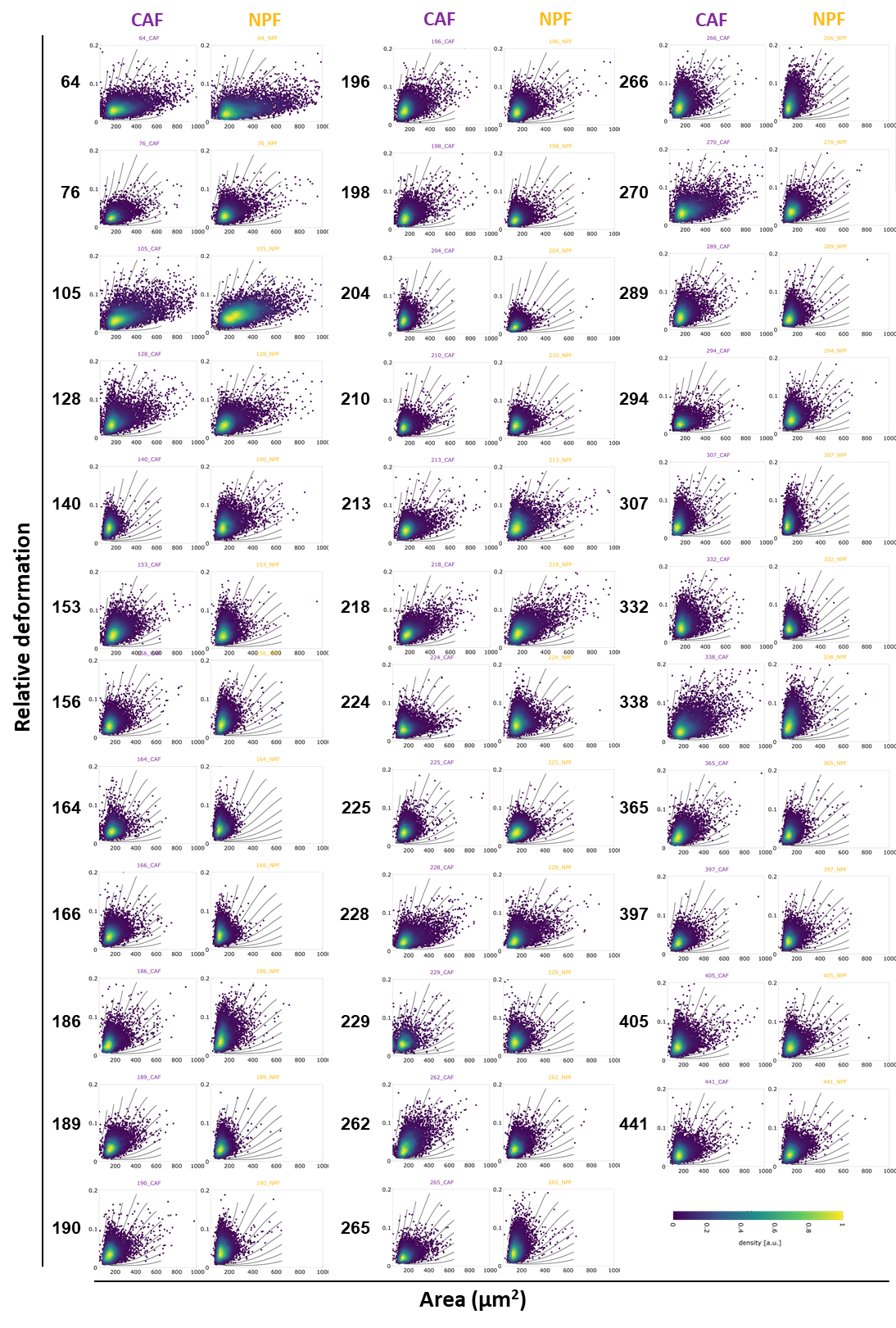

**Supplementary Figure 6. Overview of area versus deformation scatter plots of all patients (numbered on the left).** Each dot represents the measurement of one cell. Colours indicate the density of overlaid points.

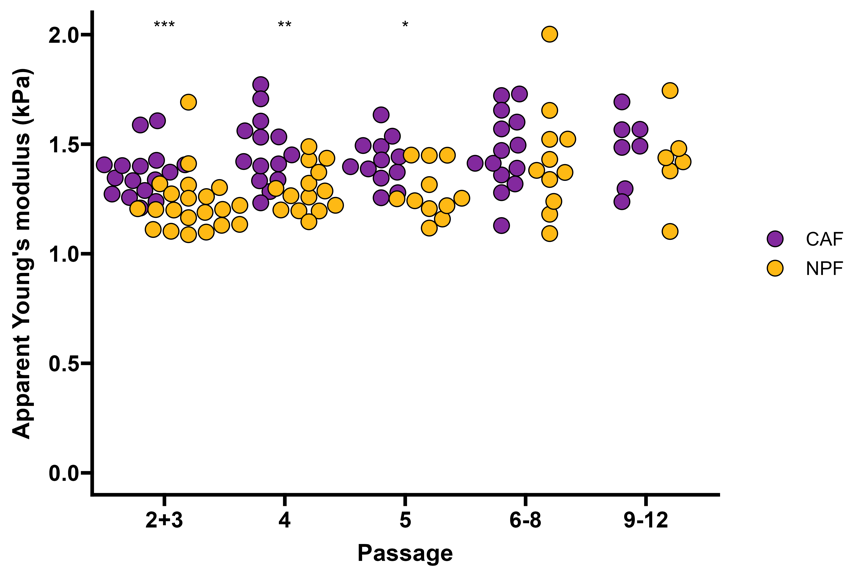

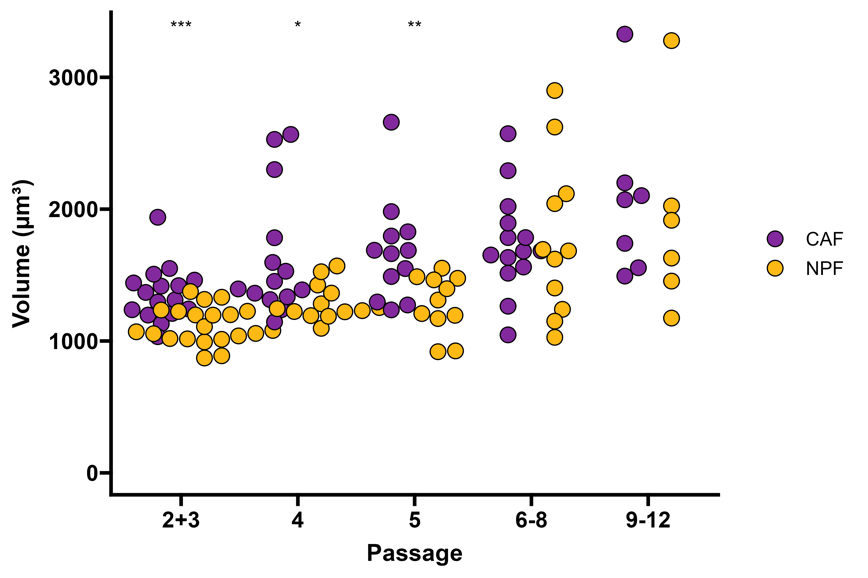

**Supplementary Figure 7. Cell volumes and apparent Young’s moduli across passage number.** Dots represent single measurements of different patients at different passages with 1 to 4 measurements per patient at different passages. Results of a Mann-Whitney test between CAFs and NPFs are shown (Volumes, passages: 2&3: *P*=0.00017; 4: *P*=0.01247; 5: *P*=0.00356; 6-8: *P*=0.93576; 9-12: *P*=0.3597; apparent Young’s modulus, passage 2+3: *P*=0.00027; 4: *P*=0.00249; 5: *P*=0.01289; 6-8: *P*=0.46682; 9-12: *P*=0.44522), * *P* <0.05. ** *P*<0.01, *** *P* <0.001.

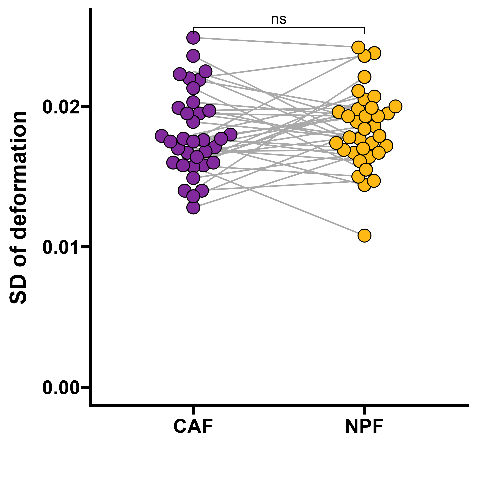

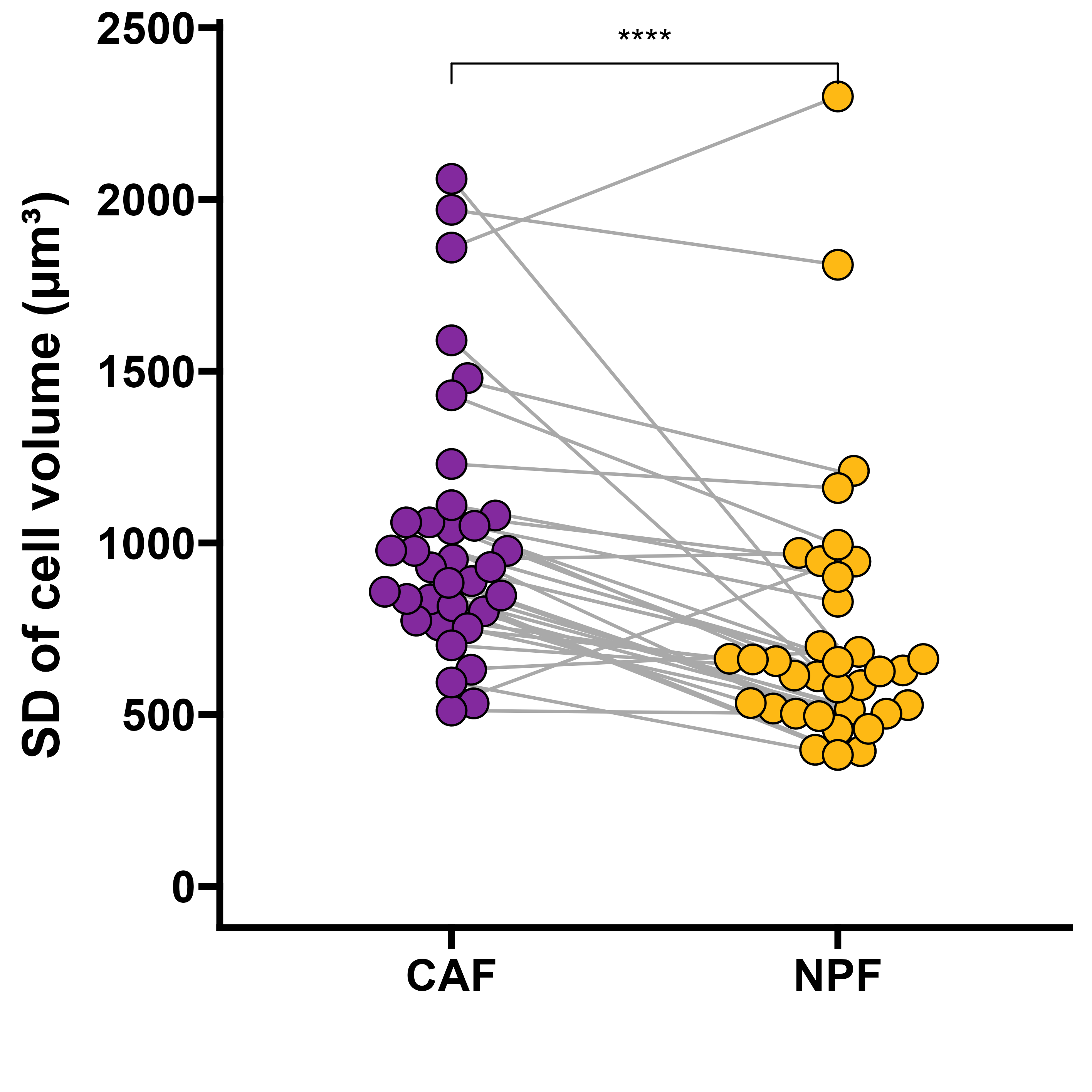

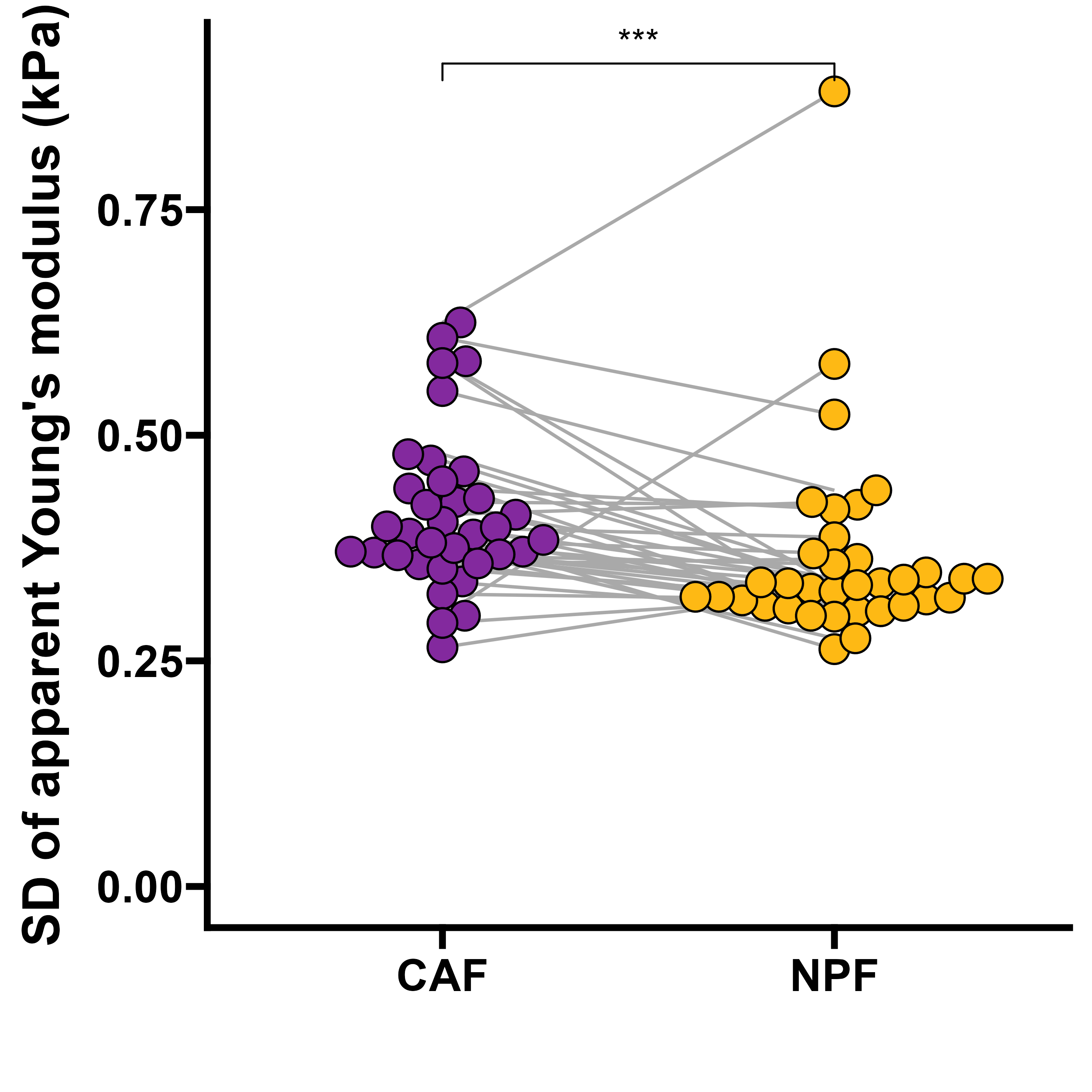

**Supplementary Figure 8.** **Standard deviations of parameters measured by RT-DC.** Standard deviations of cell volume, deformation and apparent Young’s modulus as measured by RT-DC with each dot representing one patient (n=35 pairs, medians). Results of a Wilcoxon Signed-Rank test are shown (volume: *P*=0.000031; deformation: *P*=0.62; apparent Young’s modulus: *P* = 0.00055, ns – non-significant, *** *P* <0.001, **** *P* <0.0001.

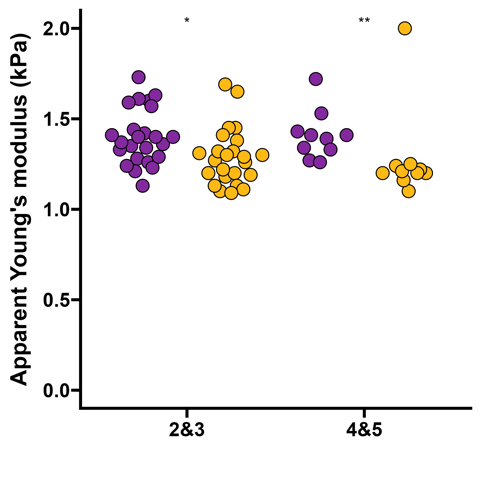

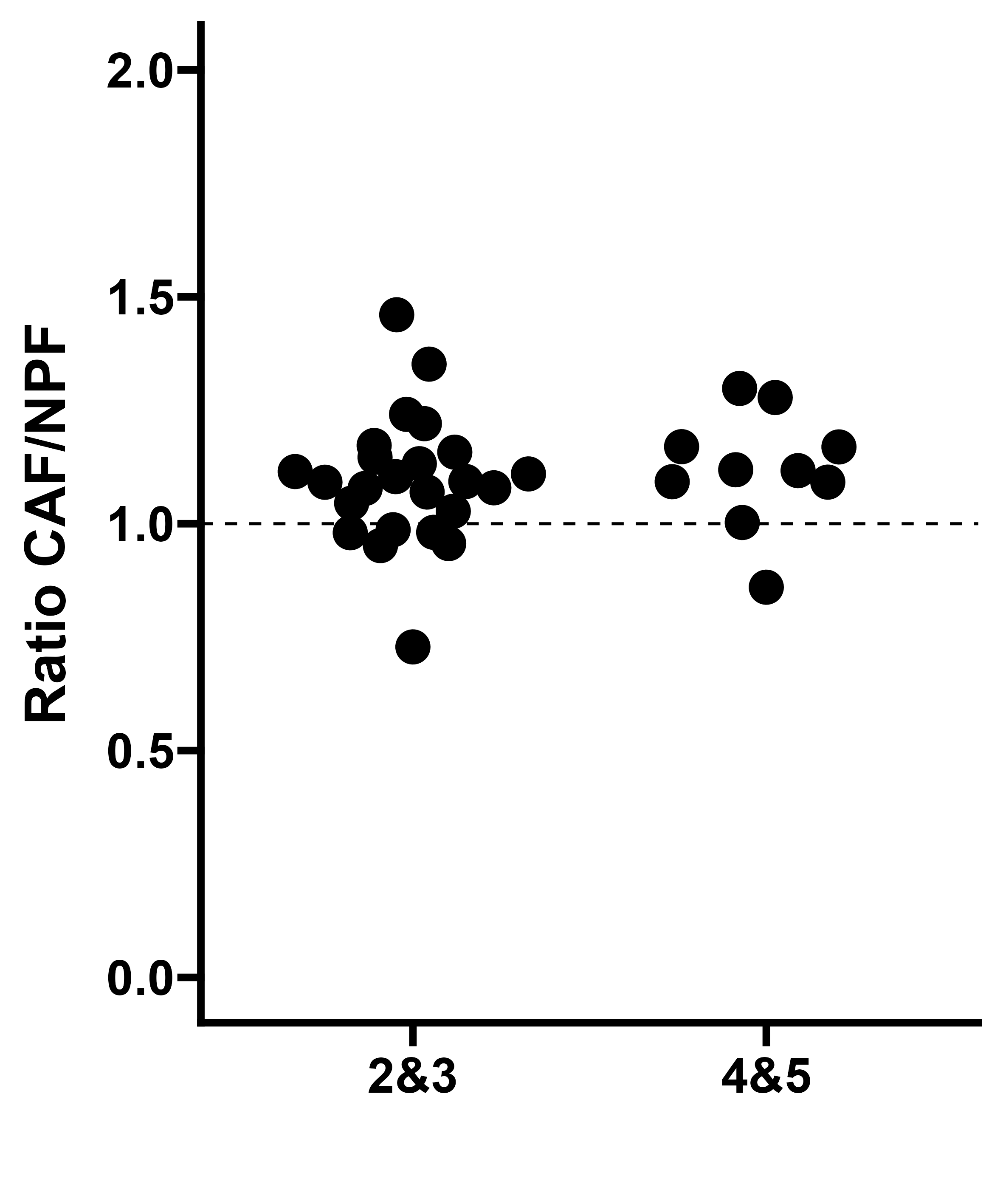

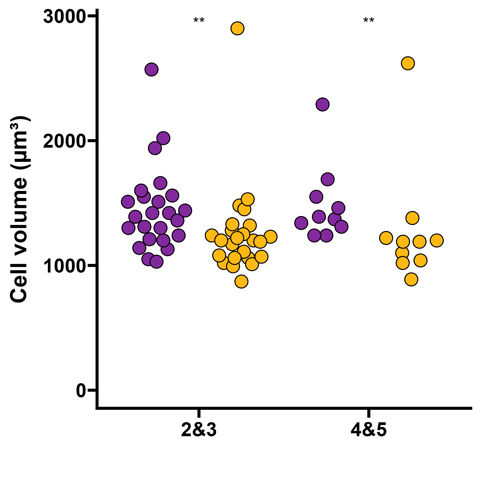

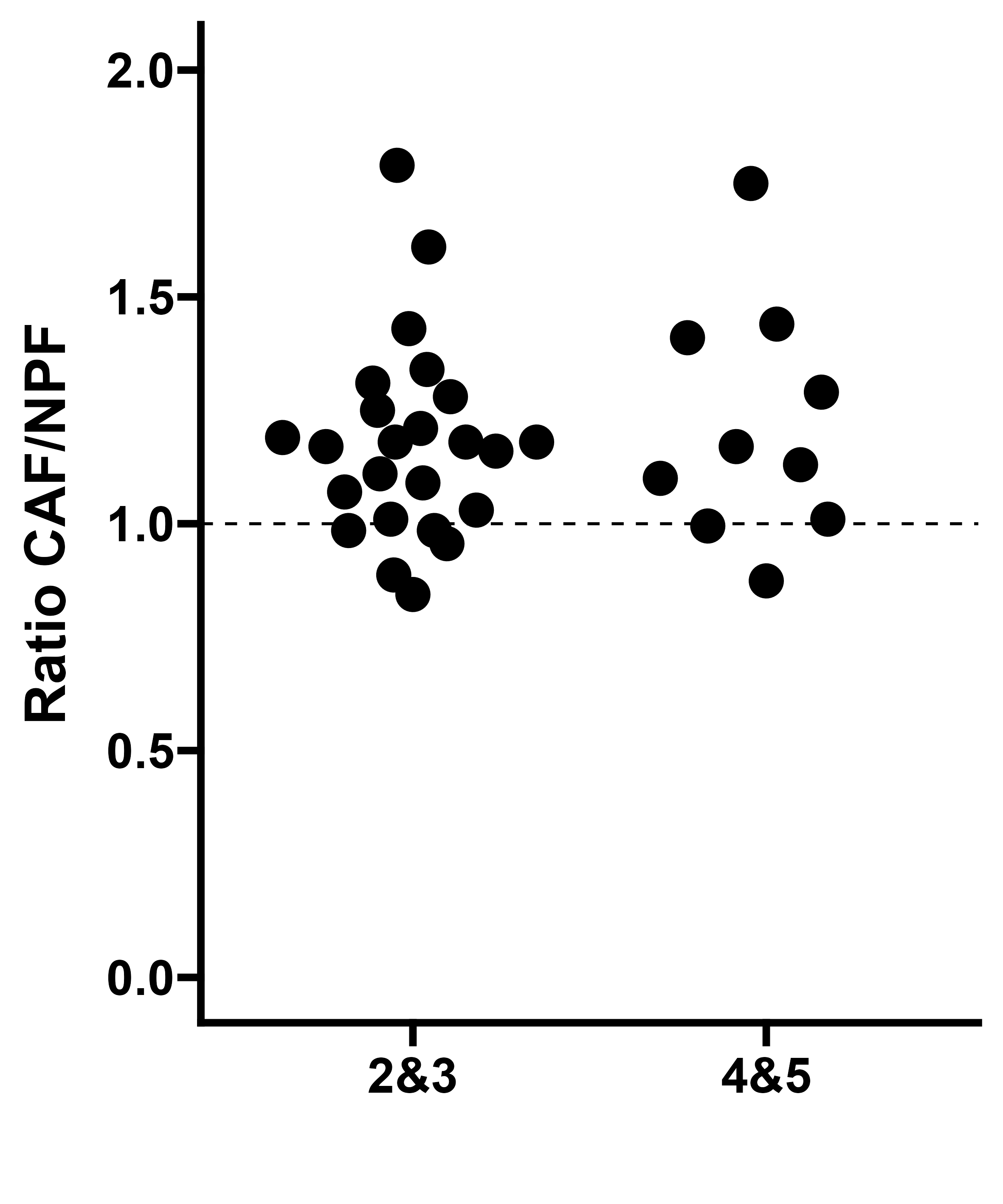

Tumor grade group

**Supplementary Figure 9. RT-DC measurements by tumour grade group.** Ratios of apparent Young’s moduli and volumes are shown on the right. Dots represent median value per patient or ratio of median values per patient (n = 34 pairs/ratios, medians). Results of a Mann Whitney test comparing CAFs and NPFs are shown for the data per patient and cell type (apparent Young’s modulus: *P*(2&3) = 0.0133; *P*(4&5) = 0.0028; volume: *P*(2&3) = 0.0047; *P*(4&5) = 0.0091), * *P*<0.05, ** *P*<0.01. Ratios were tested with a Mann Whitney test (non-significant, ratio apparent Young’s modulus*: P*=0.36; ratio volume *P*=0.81).

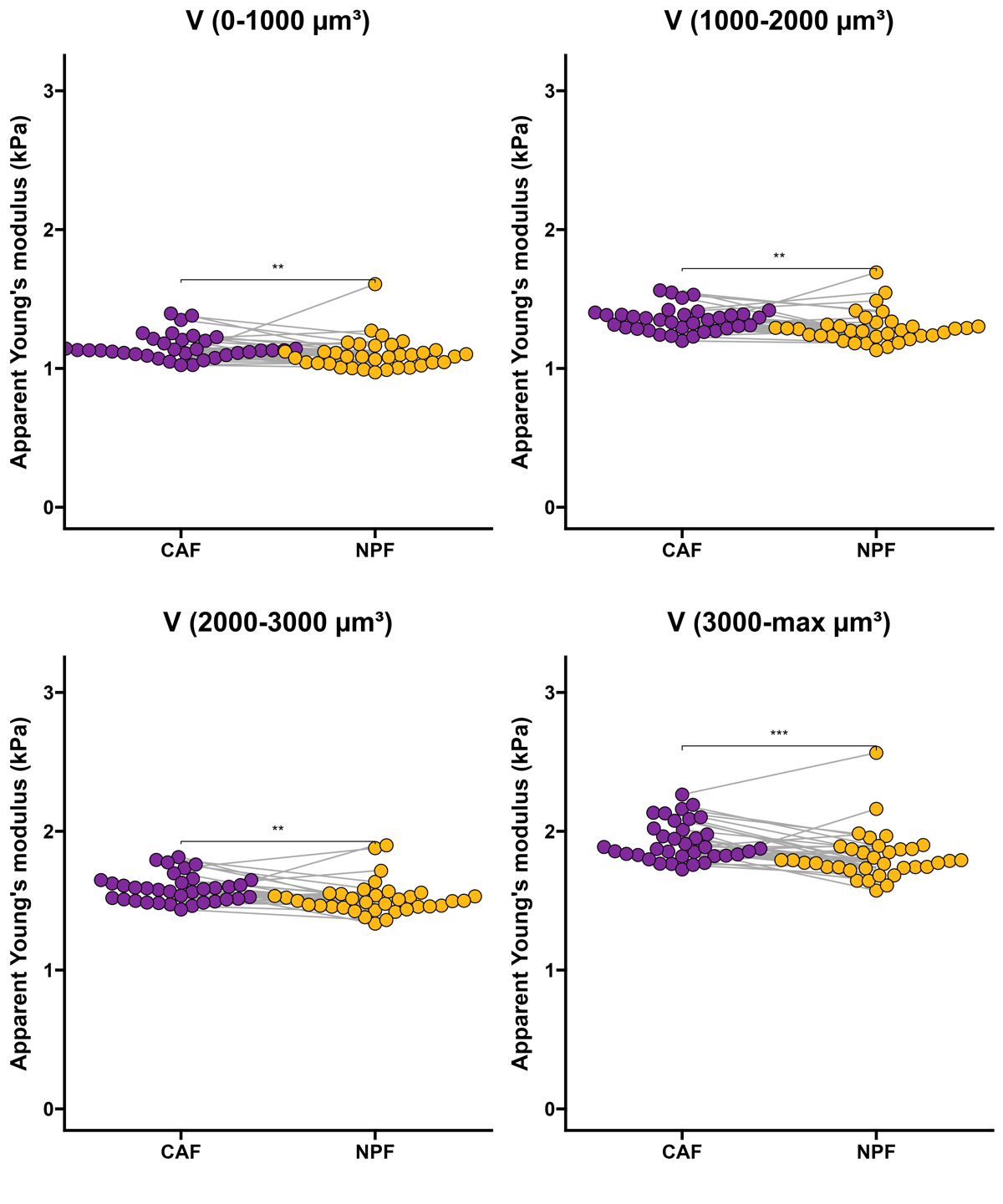

**Supplementary Figure 10. Apparent Young’s moduli for comparable volume ranges.** Individual dots represent median apparent Young’s modulus per patient binned by volume. Results of a Wilcoxon Signed-Rank test are shown. Volumes 0-1000 μm^3^: *P* = 0.0013; 1000-2000 μm^3^: *P*=0.0014; 2000-3000μm^3^: *P*=0.0028; 3000-max μm^3^: *P*= 0.00048, ** *P*<0.01, *** *P*<0.001. n=35 (donor pairs, medians).

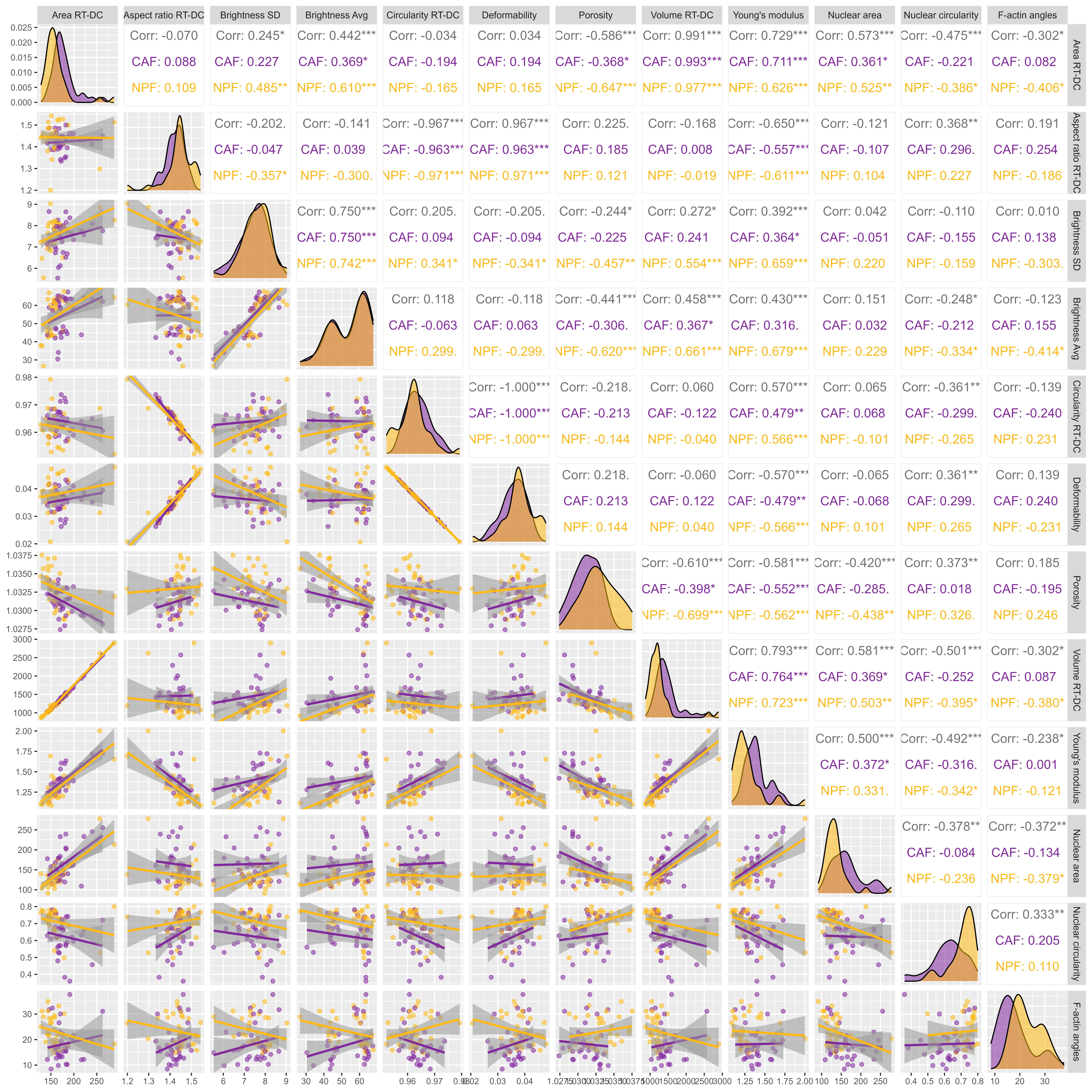

**Supplementary Figure 11. Correlation of additional RT-DC parameters and morphological parameters.** Correlation matrix with scatter plots and histograms of features extracted from RT-DC measurement and morphological parameters. Spearman correlation coefficients with associated p-values are shown (Corr. – all data, both cell types), * *P*<0.05. ** *P*<0.01, *** *P*<0.001. n = 35 donor pairs (medians).

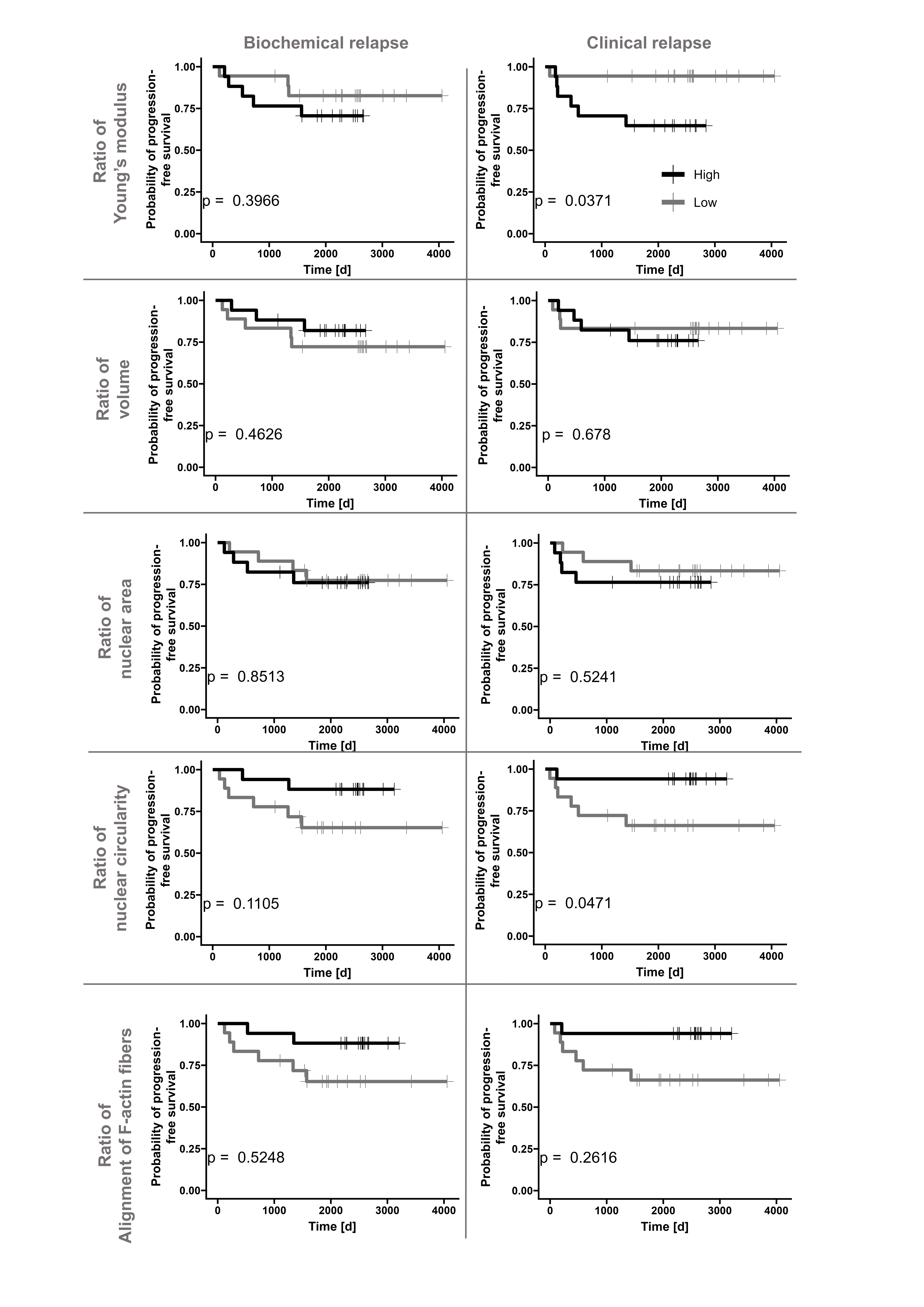

**Supplementary Figure 12. Kaplan-Meier plots of morphological and mechanical parameters.** Kaplan-Meier plots for biochemical (PSA) and clinical relapse for patients with ratios of parameter in the lower (grey, n = 18) and upper 50 percentile (black, n=17). The between-group significance was tested using a log-rank test and is shown in each plot.

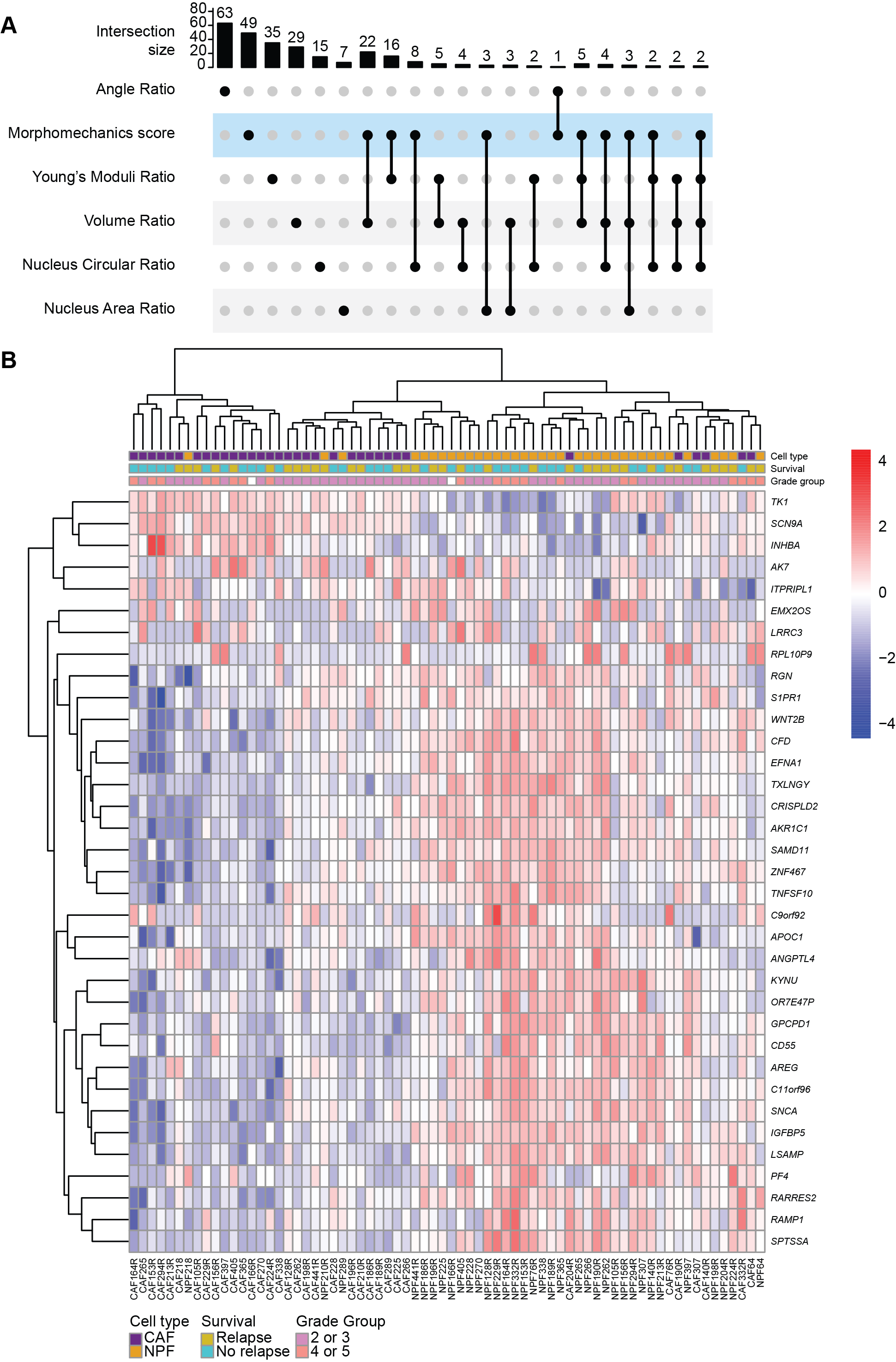

**Supplementary Figure 13. Genes associated with morphological and biomechanical features of CAFs and NPFs.** Heatmap of genes that are significantly correlated with the apparent Young’s modulus ratios and/or volume.

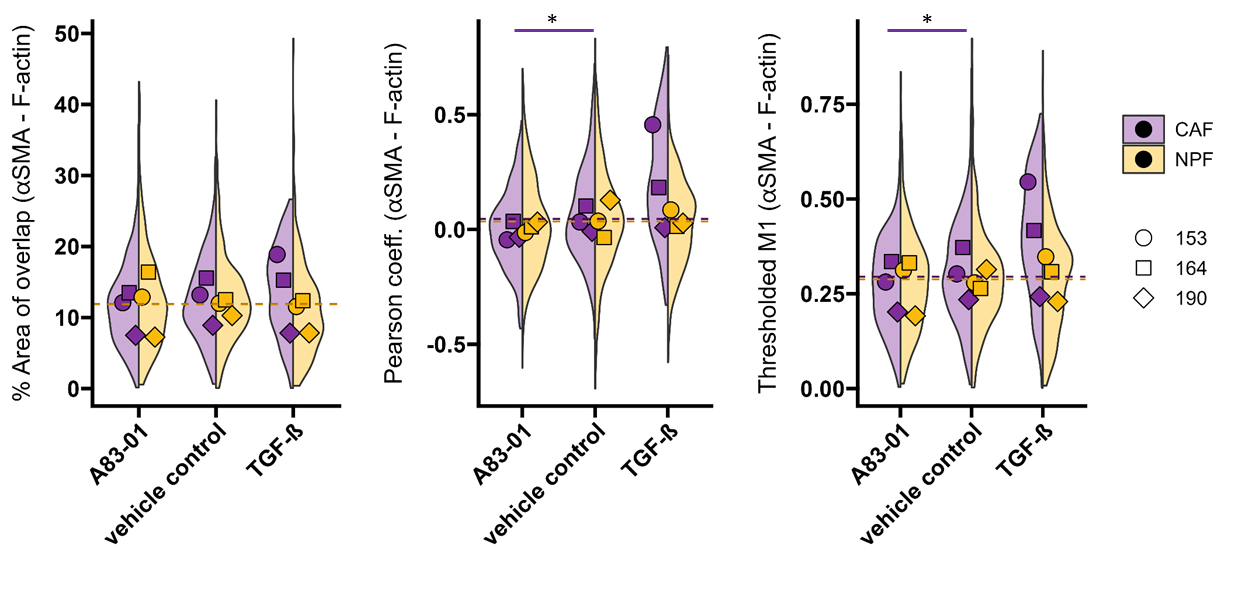

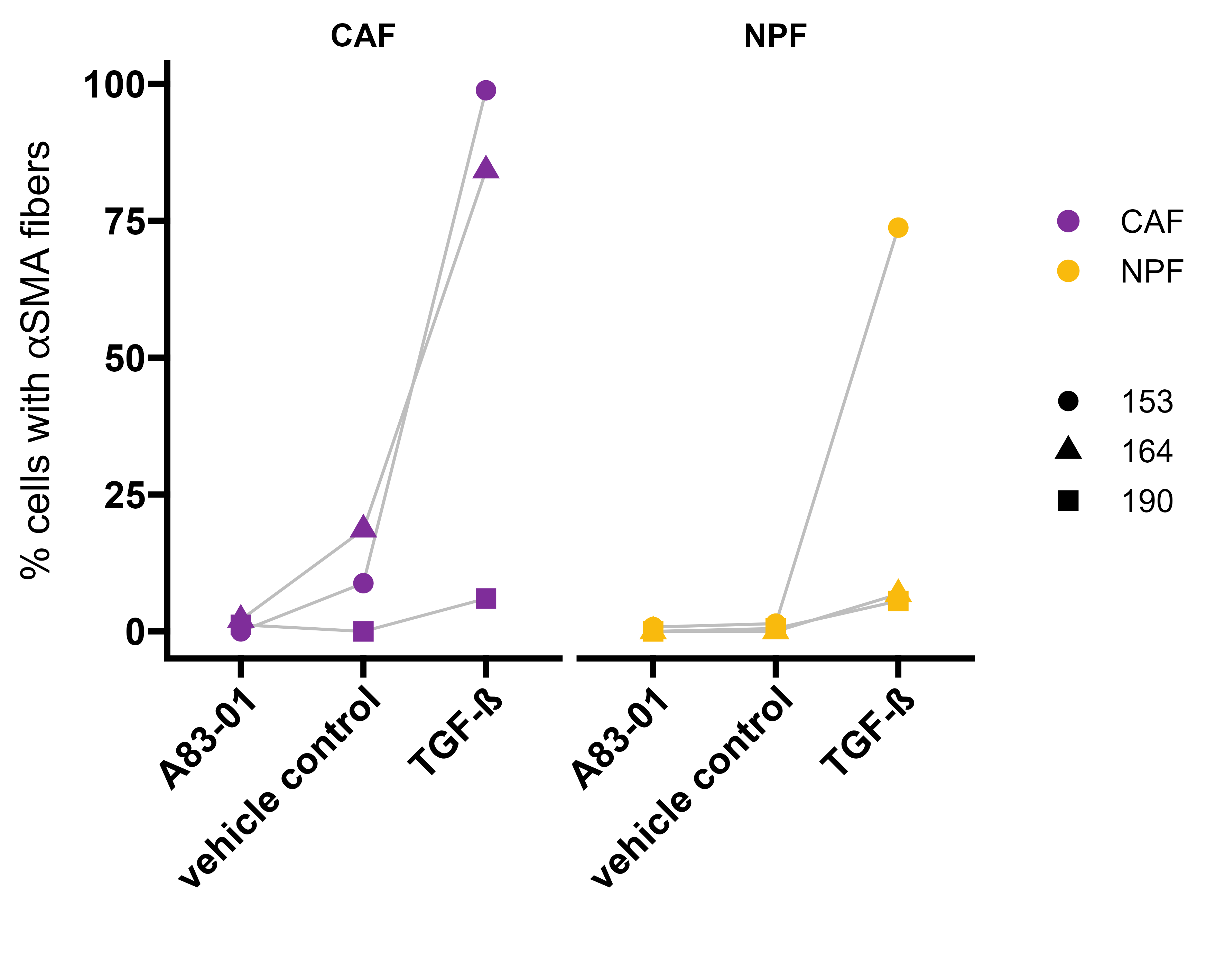

**Supplementary Figure 14. Analysis of αSMA staining after treatment with TGFβ.** Colocalization analysis of αSMA and F-actin was performed with the BIOP JACoP plugin for Fiji (https://github.com/BIOP/ijp-jacop-b). One perinuclear region of interest per cell was analysed with the automatic threshold method Otsu for both channels. Person coefficient, thresholded Manders’ coefficient 1 and percentage of overlapping area between αSMA and F-actin are shown for three patients. Percentage of cells with αSMA fibres was determined by counting cells with visible perinuclear fibres and the number of nuclei per image with 3 to 5 images analysed per condition. Medians per patient and cell type are shown. Top: Results of a likelihood-ratio test of linear mixed effects model is shown (% area: CAF, A83-01 vs ctrl: *P*=0.071; CAF TGF-β1 vs ctrl: *P*=0.55; NPF A83-01 vs ctrl: *P*=0.89; NPF TGF-β1 vs ctrl: *P*=0.13; Pearson: CAF: A83-01 vs ctrl: *P*=0.019; CAF TGF-β1 vs ctrl: *P*=0.18; NPF A83-01 vs ctrl: *P*=0.47; NPF TGF-β1 vs ctrl: *P*=0.59; tM1: CAF A83-01 vs ctrl: *P*=0.013; CAF TGF-β1 vs ctrl: *P*=0.14; NPF A83-01 vs ctrl: *P*=0.70; NPF TGF-β1 vs ctrl: *P*=0.70), * p<0.05. Bottom: Results of a Mann-Whitney test are shown (CAF A83-01 vs ctrl: *P*=0.507; CAF TGF-β1 vs ctrl *P*=0.4; CAF TGF-β1 vs A83-01: *P*=0.1; NPF A83-01 vs ctrl: *P*=0.643; NPF TGF-β1 vs ctrl: *P*=0.1; NPF TGF-β1 vs A83-01: *P*=0.076.

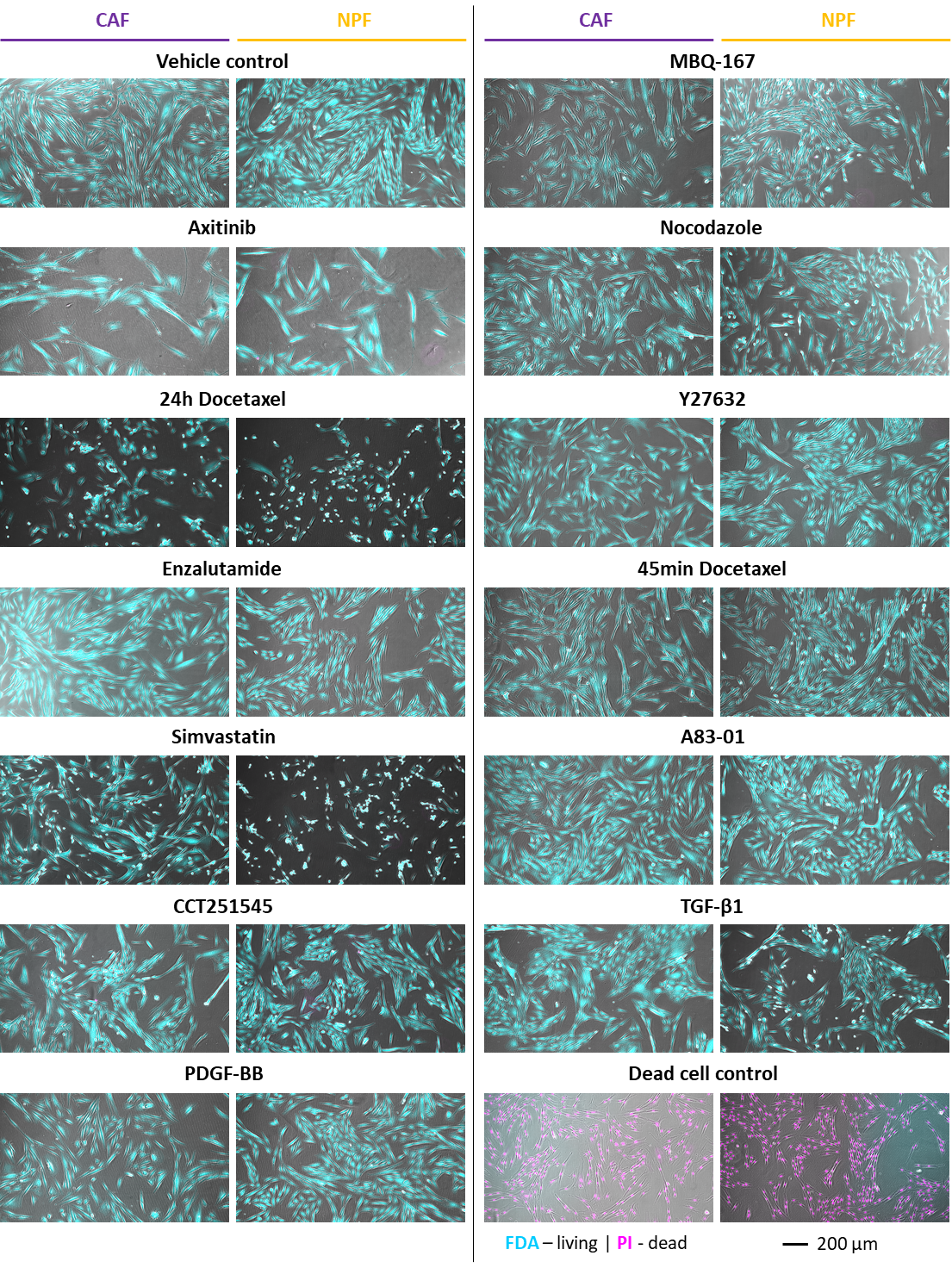

**Supplementary Figure 15. Live/dead staining of cells treated with different drugs.** Epifluorescence imaged of cells treated with different drugs. Cells were treated with the conditions mentioned in figure 6 and 7 (72h 1μM Axitinib; 24h 5nM Docetaxel; 72h 10 μM Enzalutamide; 24h 1μM Simvastatin; 72h 1μM CCT251545; 72h 25ng/mL PDFG-BB; 2h 0.5μM MBQ-167; 45min 5μM Nocodazole; 30min 10μM Y27632; 48h 5μM A83-01; 48h 10 ng/mL TGF-β1; 45min 5μM Docetaxel). After treatment, cells were stained with 8μg/mL fluorescein diacetate (FDA) and 20μg/mL propidium iodide in CO_2_-independent medium (Gibco) for 5min at room temperature.

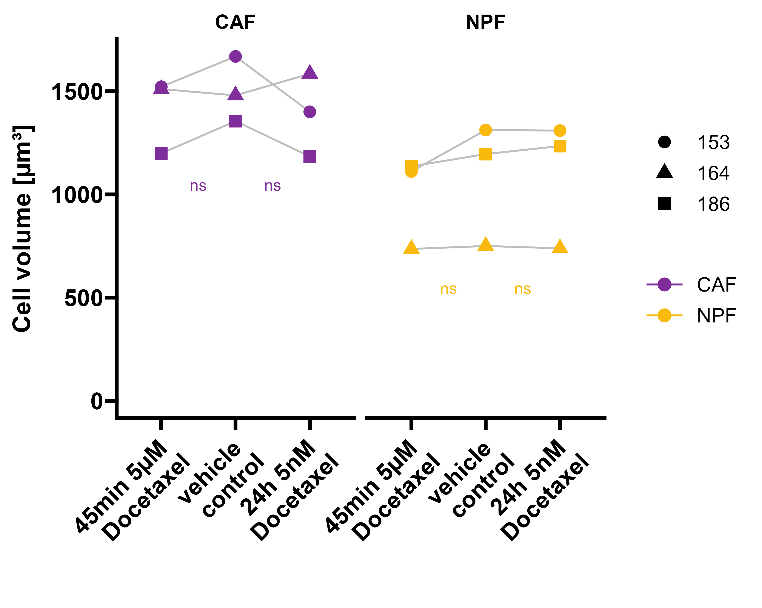

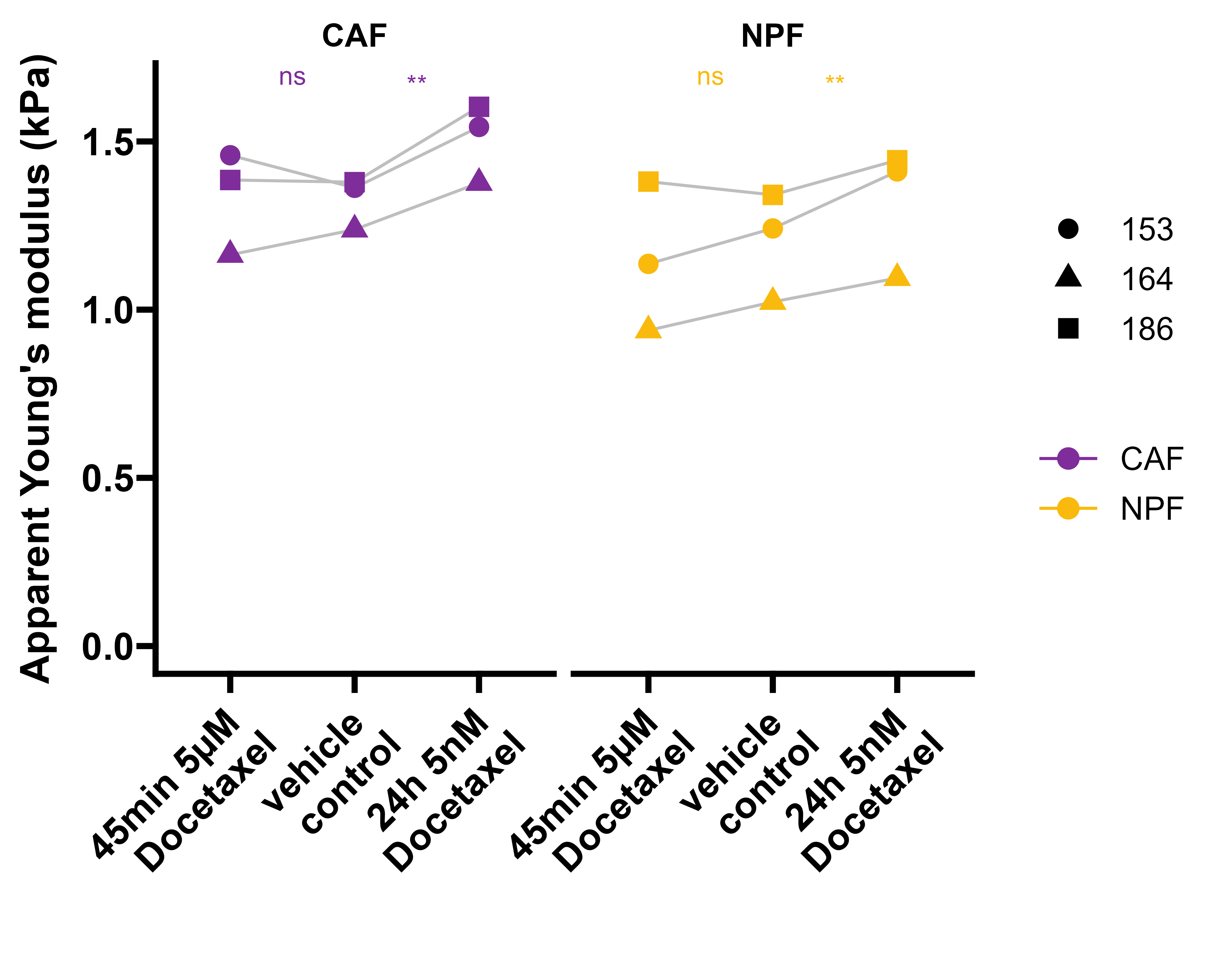

**Supplementary Figure 16. RT-DC measurement of cells treated with Docetaxel for longer time periods (24hrs) or acutely in suspension (45min).** Adherent cells were treated with 5nM Docetaxel or vehicle controls (DMSO 1:1000) for 24h; For acute treatment, cells were detached prior RT-DC measurements (as described in the methods) and treated in suspension with 5μM Docetaxel for 45min. Respective docetaxel concentrations were also present in the RT-DC assay buffer. Results of a likelihood-ratio test of linear mixed effects model is shown. (Volume: CAF 24h Docetaxel *P*=0.4749; CAF 45 min Docetaxel: *P*=0.1542; NPF 24h Docetaxel: *P*=0.2515; NPF 45min Docetaxel: *P*=0.0954; apparent Young’s modulus: CAF 24h Docetaxel: *P*=0.0021; CAF 45min Docetaxel: *P*=0.9198; NPF 24h Docetaxel: *P*=0.0067; NPF 45min Docetaxel: *P*=0.2540, ns – non-significant, ** *P*<0.01. n = 3 donors, >9500 cells for each measurement were measured. Medians of one experiment are presented.

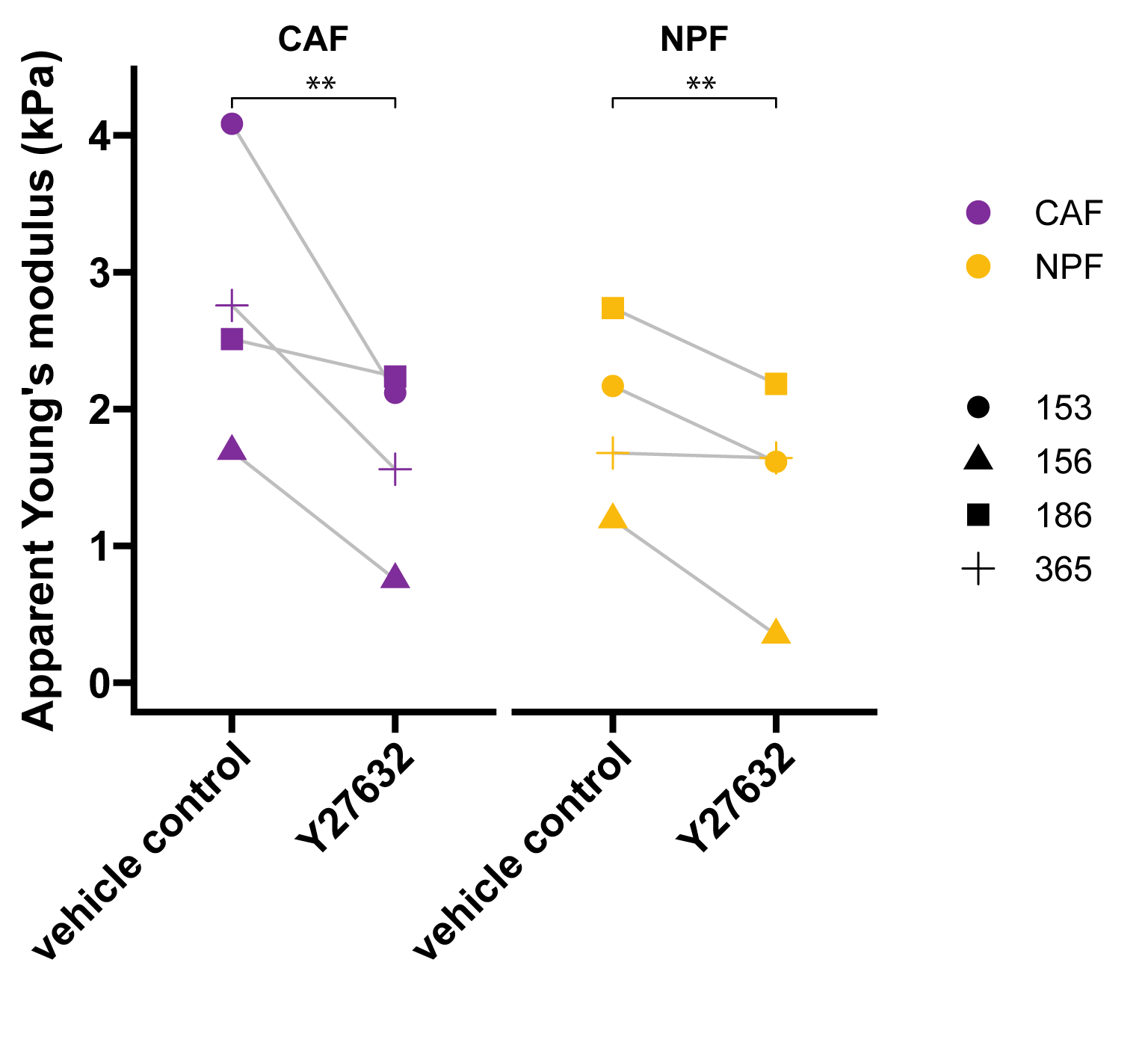

**Supplementary Figure 17. AFM probing of adherent cells after treatment with ROCK inhibitor Y27632.** Cells were seeded into cell culture dishes 24h before measurement. 10μM Y27632 or vehicle control (H_2_O) were added to the measurement medium (CO_2_-independent medium) 30 min prior measurements. Cells were probed using a spherical indenter of 5µm diameter (see methods). Results of a likelihood-ratio test of linear mixed effects model is shown (CAF: *P*=0.007; NPF: *P*=0.0024, n = 4 donors, >50 cells for each. Medians are presented.
