## Supplementary Figures and Tables for "Prostate cancer associated fibroblasts have distinct morphomechanical features that are associated with patient outcome"

**Supplementary Table 5.** Candidate genes that are significantly correlated with the morphomechanical score.

#All pathways with adjusted P value < 0.05; NES = normalised enrichment score

| Geneset collection | Pathway | Adjusted P value | NES | pval | size |
|---|---|---|---|---|---|
| GO Cellular Component | SPINDLE_POLE | 0.012870276 | 2.744886814 | 0.00026001 | 18 |
| GO Cellular Component | CONDENSED_CHROMOSOME_CENTROMERIC_REGION | 0.012870276 | 2.64807667 | 0.000264131 | 19 |
| GO Cellular Component | SPINDLE | 0.012870276 | 2.638706104 | 0.000281611 | 26 |
| GO Cellular Component | CHROMOSOMAL_REGION | 0.012870276 | 2.627846578 | 0.000281611 | 26 |
| GO Cellular Component | CONDENSED_CHROMOSOME | 0.012870276 | 2.568480715 | 0.000276702 | 28 |
| GO Cellular Component | CHROMOSOME_CENTROMERIC_REGION | 0.012870276 | 2.545057994 | 0.000276932 | 27 |
| GO Cellular Component | CENTROSOME | 0.012870276 | 2.459725501 | 0.000289519 | 32 |
| GO Cellular Component | MITOTIC_SPINDLE | 0.012870276 | 2.397884348 | 0.000255885 | 13 |
| GO Cellular Component | MICROTUBULE_ORGANIZING_CENTER | 0.012870276 | 2.39401086 | 0.000303398 | 39 |
| GO Cellular Component | MICROTUBULE_CYTOSKELETON | 0.012870276 | 2.388545129 | 0.000360881 | 66 |
| GO Cellular Component | POLYMERIC_CYTOSKELETAL_FIBER | 0.012870276 | 2.301517983 | 0.000298775 | 41 |
| GO Cellular Component | MIDBODY | 0.012870276 | 2.288841208 | 0.000253614 | 14 |
| GO Cellular Component | MICROTUBULE_ASSOCIATED_COMPLEX | 0.012870276 | 2.169521198 | 0.000263089 | 16 |
| GO Cellular Component | MICROTUBULE | 0.012870276 | 2.07069283 | 0.000285878 | 27 |
| GO Cellular Component | INTRACELLULAR_PROTEIN_CONTAINING_COMPLEX | 0.012870276 | 2.043458097 | 0.000273523 | 13 |
| GO Cellular Component | SUPRAMOLECULAR_COMPLEX | 0.012870276 | 2.030901454 | 0.000367512 | 59 |
| GO Cellular Component | NUCLEAR_UBIQUITIN_LIGASE_COMPLEX | 0.014506102 | 2.013798488 | 0.000460511 | 4 |
| GO Cellular Component | INTERCELLULAR_BRIDGE | 0.030844553 | 2.00240272 | 0.00122399 | 8 |
| GO Cellular Component | CENTRIOLAR_SATELLITE | 0.027377169 | 1.974440624 | 0.000977756 | 11 |
| GO Cellular Component | MITOTIC_SPINDLE_POLE | 0.020085372 | 1.969781723 | 0.000690767 | 4 |
| GO Cellular Component | SPINDLE_MICROTUBULE | 0.030844553 | 1.961427502 | 0.001197892 | 9 |
| GO Cellular Component | DYNEIN_COMPLEX | 0.048098946 | 1.95121026 | 0.002354049 | 4 |
| GO Cellular Component | CENTRIOLE | 0.0393284 | 1.936407643 | 0.001702749 | 10 |
| GO Cellular Component | UBIQUITIN_LIGASE_COMPLEX | 0.0393284 | 1.867563812 | 0.00175219 | 7 |
| GO Cellular Component | CATALYTIC_COMPLEX | 0.012870276 | 1.84662803 | 0.000296121 | 22 |
| GO Cellular Component | NUCLEAR_CHROMOSOME | 0.031870226 | 1.840585804 | 0.001306848 | 13 |
| GO Cellular Component | NMS_COMPLEX | 0.042905789 | 1.834747695 | 0.002043133 | 3 |
| GO Cellular Component | DNA_REPAIR_COMPLEX | 0.0393284 | 1.754099764 | 0.001768738 | 4 |
| GO Cellular Component | PRONUCLEUS | 0.014101058 | 1.739917968 | 0.000429 | 3 |
| GO Cellular Component | CILIUM | 0.012870276 | 1.739767044 | 0.000292227 | 27 |
| GO Cellular Component | SUPRAMOLECULAR_POLYMER | 0.020085372 | 1.736439353 | 0.000673401 | 42 |
| GO Cellular Component | CONTRACTILE_RING | 0.04068648 | 1.531356249 | 0.001883633 | 2 |
| GO Cellular Component | CHROMOSOME | 0.030844553 | 1.483824812 | 0.001192843 | 55 |
| GO Cellular Component | INSULIN_LIKE_GROWTH_FACTOR_BINDING_PROTEIN_COMPLEX | 0.012870276 | -1.640381426 | 0.000374532 | 1 |
| GO Cellular Component | GROWTH_FACTOR_COMPLEX | 0.012870276 | -1.640381426 | 0.000374532 | 1 |
| GO Cellular Component | CHROMATIN | 0.012870276 | -1.672372734 | 0.000280741 | 48 |
| GO Cellular Component | TRANSCRIPTION_REGULATOR_COMPLEX | 0.012870276 | -1.995264045 | 0.00031294 | 17 |
| KEGG | CELL_CYCLE | 0.010754017 | 2.414898999 | 0.000247219 | 24 |
| KEGG | P53_SIGNALING_PATHWAY | 0.010754017 | 2.372224502 | 0.000235793 | 12 |
| KEGG | DNA_REPLICATION | 0.010754017 | 1.991096388 | 0.000225734 | 7 |
| KEGG | OOCYTE_MEIOSIS | 0.02361179 | 1.98499157 | 0.000741656 | 24 |
| KEGG | CHEMOKINE_SIGNALING_PATHWAY | 0.02361179 | -1.873376117 | 0.0008142 | 34 |
| KEGG | ARACHIDONIC_ACID_METABOLISM | 0.026257545 | -1.925013754 | 0.001056338 | 10 |
| KEGG | ERBB_SIGNALING_PATHWAY | 0.010754017 | -2.066358572 | 0.000173581 | 12 |
| GO Biological Process | REGULATION_OF_CHROMOSOME_SEGREGATION | 0.022376533 | 3.081884292 | 0.000261849 | 35 |
| GO Biological Process | SISTER_CHROMATID_SEGREGATION | 0.022376533 | 3.074986465 | 0.000277085 | 56 |
| GO Biological Process | CHROMOSOME_ORGANIZATION | 0.022376533 | 3.05546652 | 0.000304229 | 100 |

**Supplementary Table 6.** Geneset enrichment analysis based on the correlation of gene ratios with the morphomechanical score.

| Feature 1 | Feature 2 | Correlation (r) | P value |
|---|---|---|---|
| iCAF | myCAF | 0.07 | 0.633127 |
| iCAF | morphomechanics score | -0.13 | 0.308822 |
| iCAF | ratio_angle | 0.32 | 0.071517 |
| iCAF | ratio_nuc_area | -0.3 | 0.226878 |
| iCAF | ratio_nuc_circ | -0.17 | 0.901766 |
| iCAF | ratio_volume | 0.01 | 0.838264 |
| iCAF | ratio_young | -0.1 | 0.527328 |
| myCAF | morphomechanics score | 0.36 | 0.03837 |
| myCAF | ratio_angle | 0.14 | 0.582547 |
| myCAF | ratio_nuc_area | 0.3 | 0.157802 |
| myCAF | ratio_nuc_circ | -0.17 | 0.238442 |
| myCAF | ratio_volume | 0.25 | 0.144083 |
| myCAF | ratio_young | 0.43 | 0.004371 |

**Supplementary Table 7.** Correlation between CAF subtype signatures and morphological and biomechanical features.

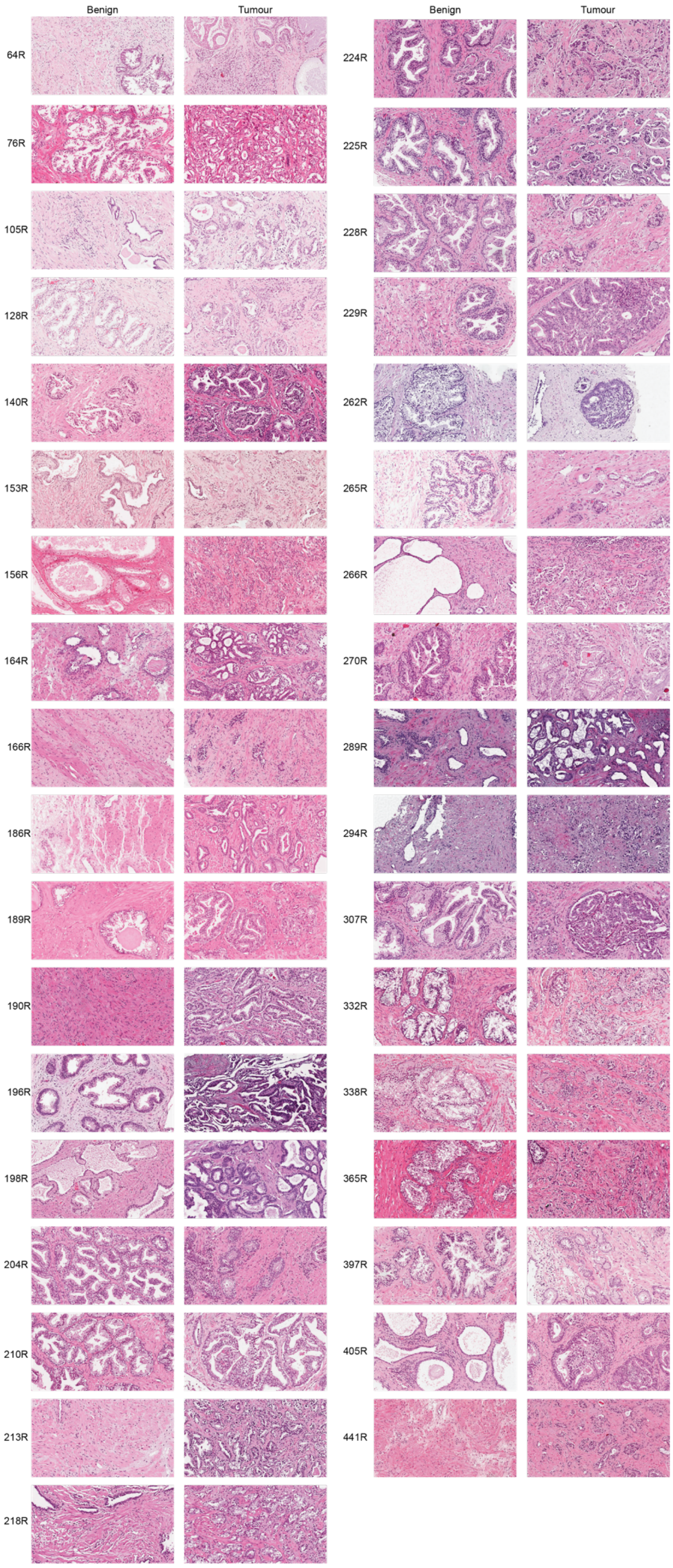


**Supplementary Figure 1. Summary of the pathology of each patient sample.** Representative images of haematoxylin and eosin stained tissue from the benign and tumour region for each patient. Small pieces of tissue were retained from each sample, while the remaining tissue was digested to establish primary cultures of CAFs and NPFs.

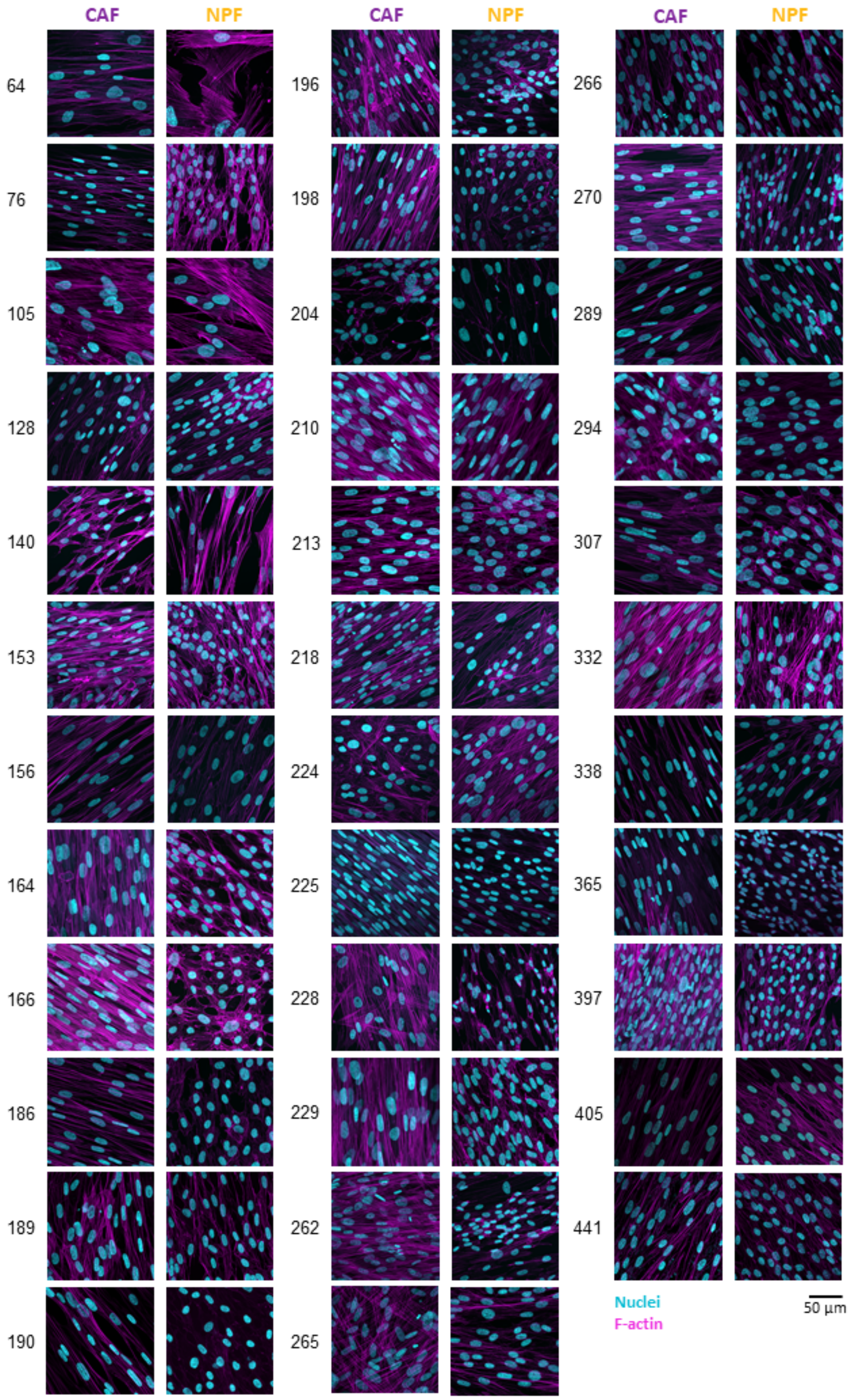


**Supplementary Figure 2. Overview of CAF and NPF morphology.** Representative images of CAFs and NPFs after 14 days in culture and stained with DAPI (DNA - cyan) and phalloidin-TRITC (F-actin - magenta). Images were recorded using a confocal microscope (Zeiss LSM780).

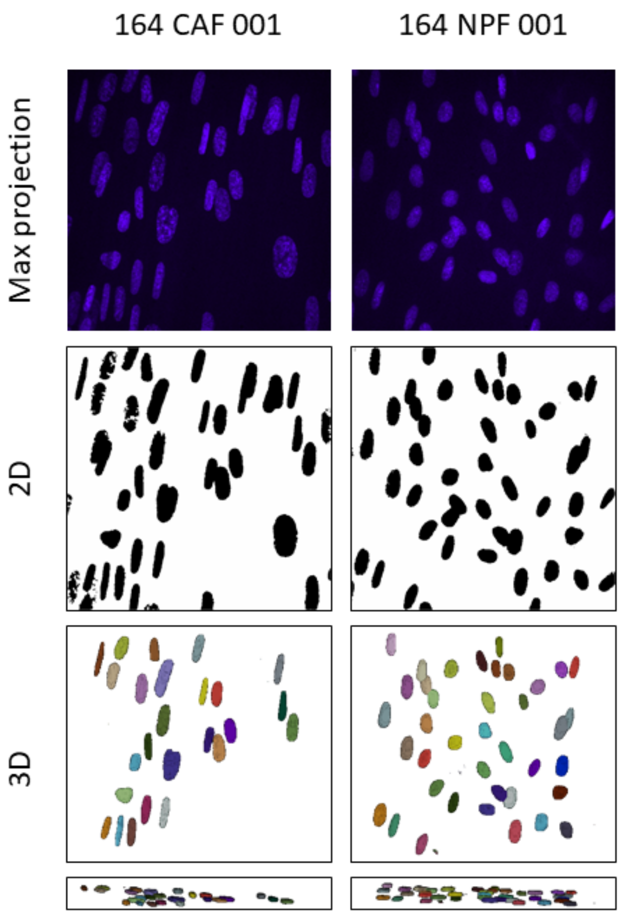


**Supplementary Figure 3. Segmentation of nuclei for morphometric analysis.** Example images for the analysis for comparison of nuclear projected area and volume. CAF/NPF cultures were stained for their nuclei (blue) using DAPI and imaged using a confocal microscope (top). 3D projected nuclei were segmented with Fiji (version 1.53o) (middle). 3D segmentation was conducted with Cellpose (version 2.2) and nuclei were viewed with Napari (version 0.4.17) (bottom).

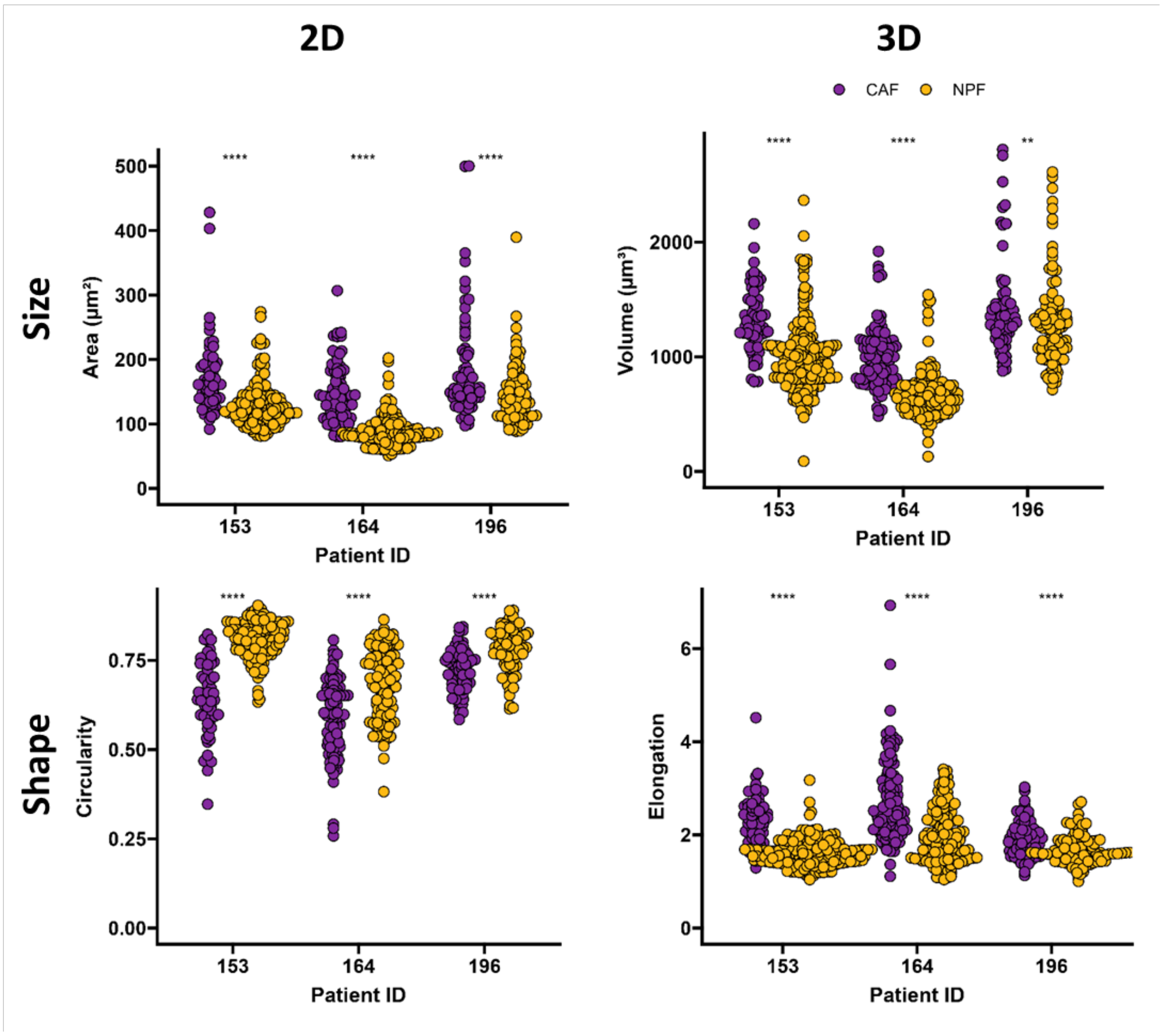


**Supplementary Figure 4. Comparison of CAFs (purple) and NPFs (orange) size and shape descriptors of the same images.** Three representative patients are shown. Each dot represents one nucleus. $n_{2D}$ = 65 - 165, $n_{3D}$ = 74 – 218. Results of a Mann-Whitney test shown (****: p<0.001, **: p<0.01)). 2D Area: 153: $P$=1.4·10$^{-12}$, 164: $P$< 2·10$^{-16}$, 196: $P$=1.6·10$^{-5}$; 2D Circularity: 153: $P$< 2·10$^{-16}$, 164: $P$=8.3·10$^{-12}$, 196: $P$=9.1·10$^{-11}$; 3D Volume: 153: $P$<2 · 10$^{-16}$, 164: $P$<2·10$^{-16}$, 196: $P$=0.0074; 3D Elongation: 153: $P$<2 · 10$^{-16}$, 164: $P$<2·10$^{-16}$, 196: $P$=3·10$^{-6}$.

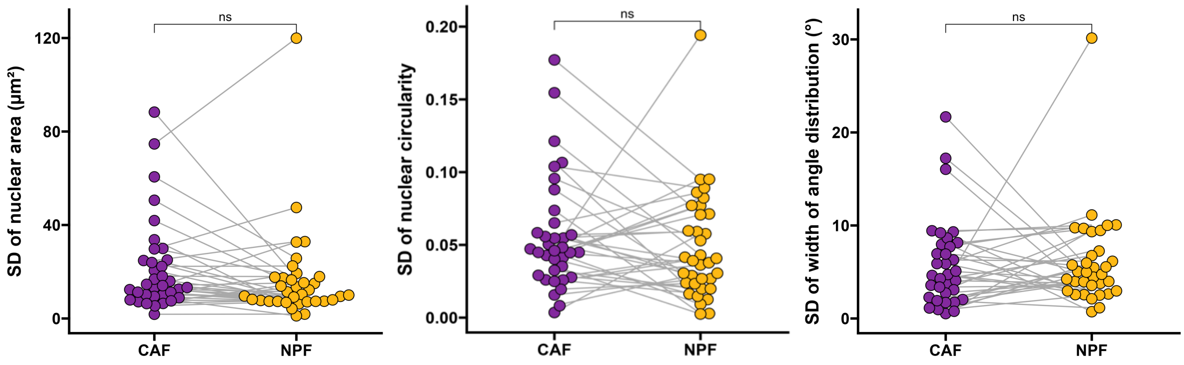


**Supplementary Figure 5. Standard deviations of morphological parameters.** Standard deviations of nuclear area, circularity and angle distribution were calculated per patient and cell type. Results of a Wilcoxon Signed-Rank test are shown (ns – non-significant, $P$(area) = 0.097; $P$(circ.) = 0.29; $P$(angle distr.) = 0.64). n=35 (donor pairs, medians).

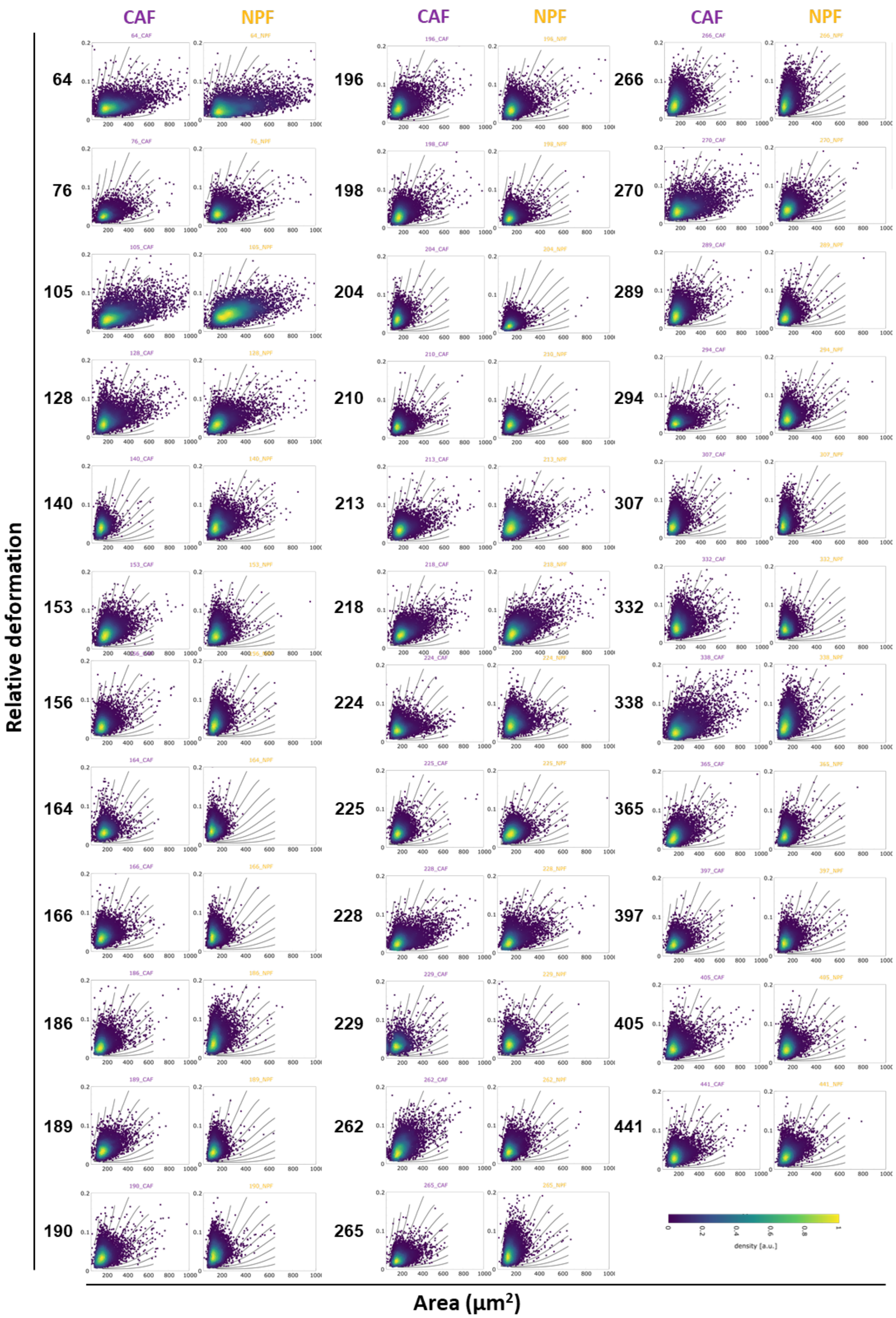


**Supplementary Figure 6. Overview of area versus deformation scatter plots of all patients (numbered on the left).** Each dot represents the measurement of one cell. Colours indicate the density of overlaid points.

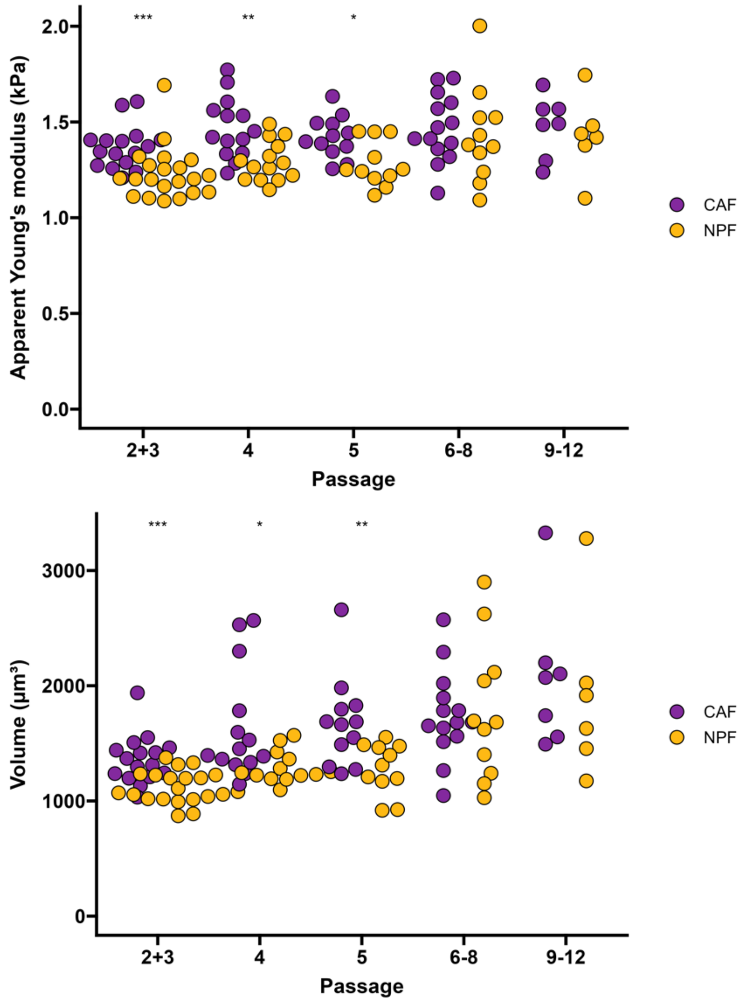


**Supplementary Figure 7. Cell volumes and apparent Young's moduli across passage number.** Dots represent single measurements of different patients at different passages with 1 to 4 measurements per patient at different passages. Results of a Mann-Whitney test between CAFs and NPFs are shown (Volumes, passages: 2&3: *P*=0.00017; 4: *P*=0.01247; 5: *P*=0.00356; 6-8: *P*=0.93576; 9-12: *P*=0.3597; apparent Young's modulus, passage 2+3: *P*=0.00027; 4: *P*=0.00249; 5: *P*=0.01289; 6-8: *P*=0.46682; 9-12: *P*=0.44522), * *P* <0.05. ** *P*<0.01, *** *P* <0.001.

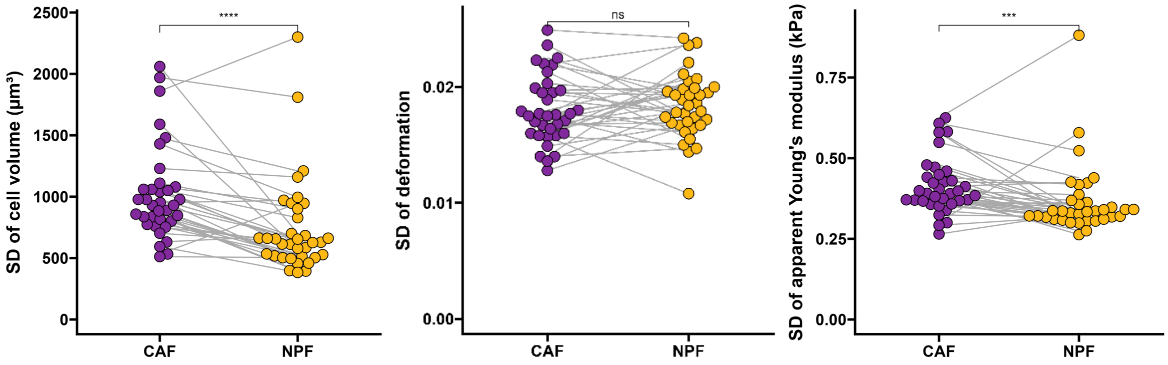


**Supplementary Figure 8. Standard deviations of parameters measured by RT-DC.** Standard deviations of cell volume, deformation and apparent Young's modulus as measured by RT-DC with each dot representing one patient (n=35 pairs, medians). Results of a Wilcoxon Signed-Rank test are shown (volume: $P$=0.000031; deformation: $P$=0.62; apparent Young's modulus: $P$ = 0.00055, ns – non-significant, *** $P$ <0.001, **** $P$ <0.0001.

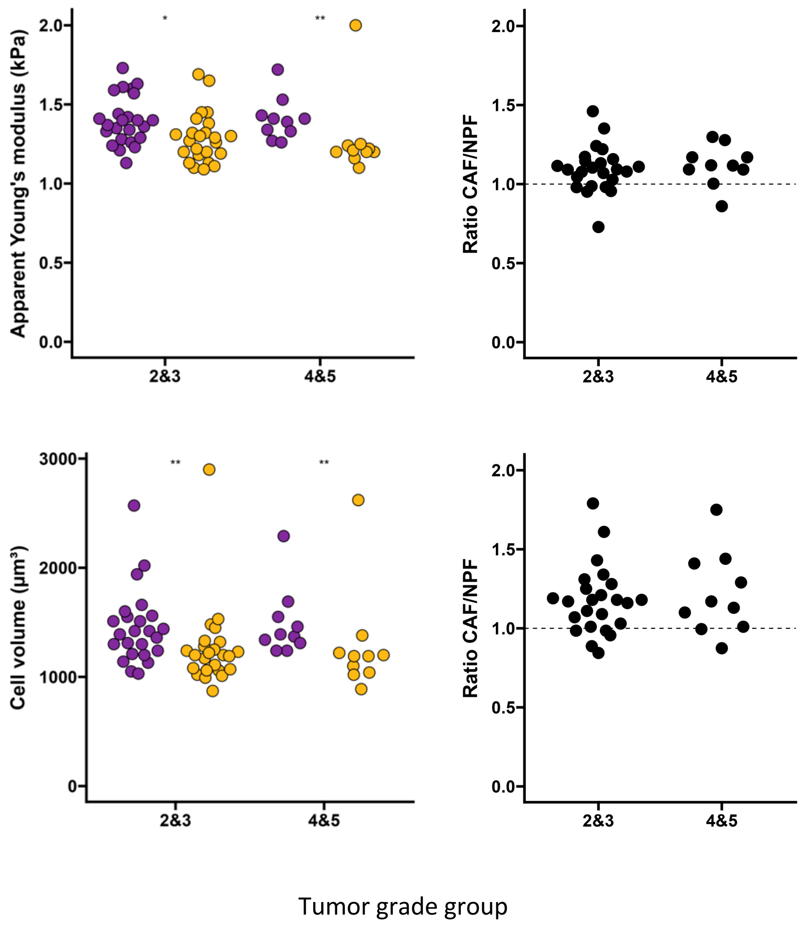


**Supplementary Figure 9. RT-DC measurements by tumour grade group.** Ratios of apparent Young's moduli and volumes are shown on the right. Dots represent median value per patient or ratio of median values per patient (n = 34 pairs/ratios, medians). Results of a Mann Whitney test comparing CAFs and NPFs are shown for the data per patient and cell type (apparent Young's modulus: $P$(2&3) = 0.0133; $P$(4&5) = 0.0028; volume: $P$(2&3) = 0.0047; $P$(4&5) = 0.0091), * $P<0.05$, ** $P<0.01$. Ratios were tested with a Mann Whitney test (non-significant, ratio apparent Young's modulus: $P=0.36$; ratio volume $P=0.81$).

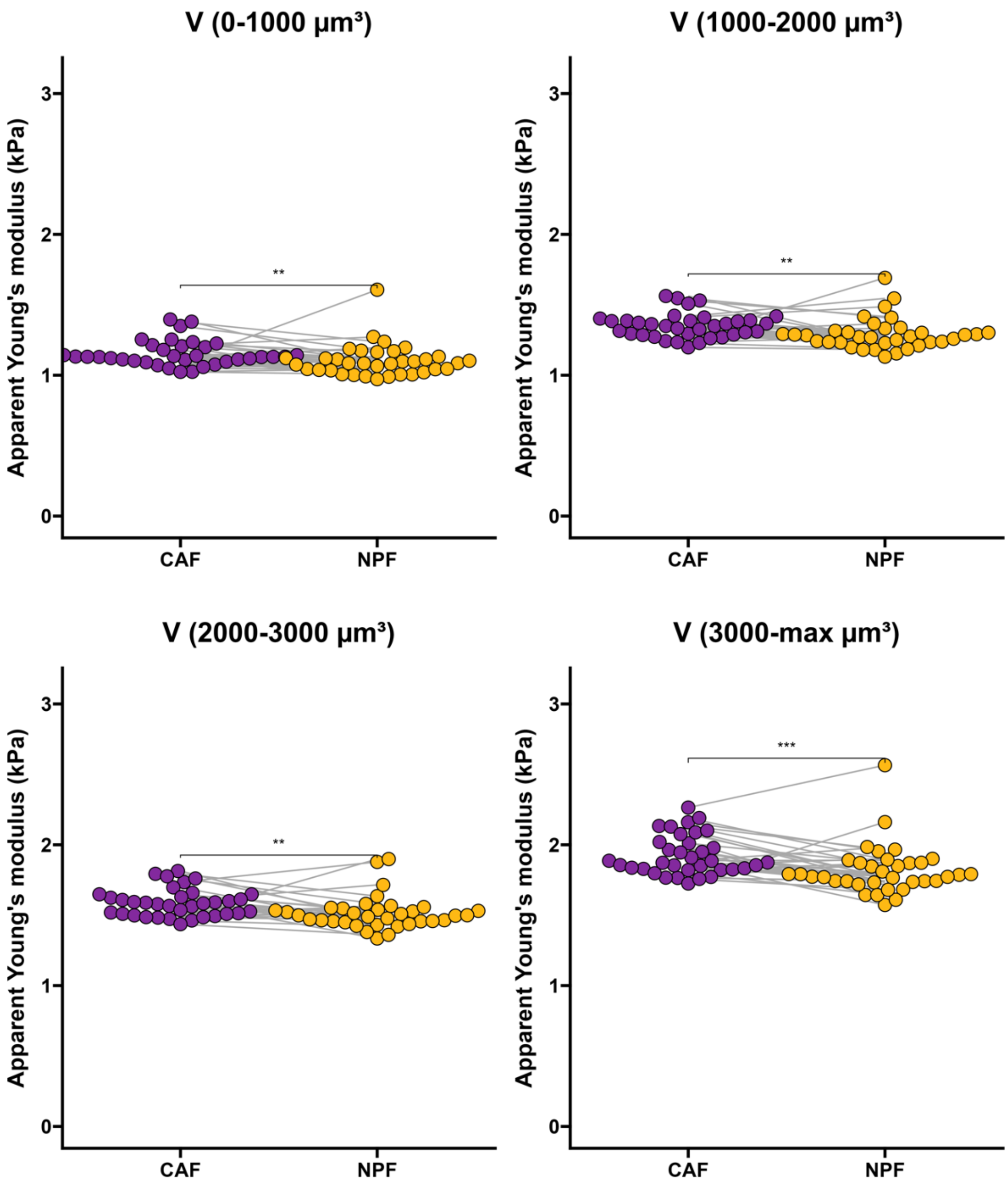


**Supplementary Figure 10. Apparent Young's moduli for comparable volume ranges.** Individual dots represent median apparent Young's modulus per patient binned by volume. Results of a Wilcoxon Signed-Rank test are shown. Volumes 0-1000 µm$^3$: *P* = 0.0013; 1000-2000 µm$^3$: *P*=0.0014; 2000-3000µm$^3$: *P*=0.0028; 3000-max µm$^3$: *P*= 0.00048, ** *P*<0.01, *** *P*<0.001. n=35 (donor pairs, medians).

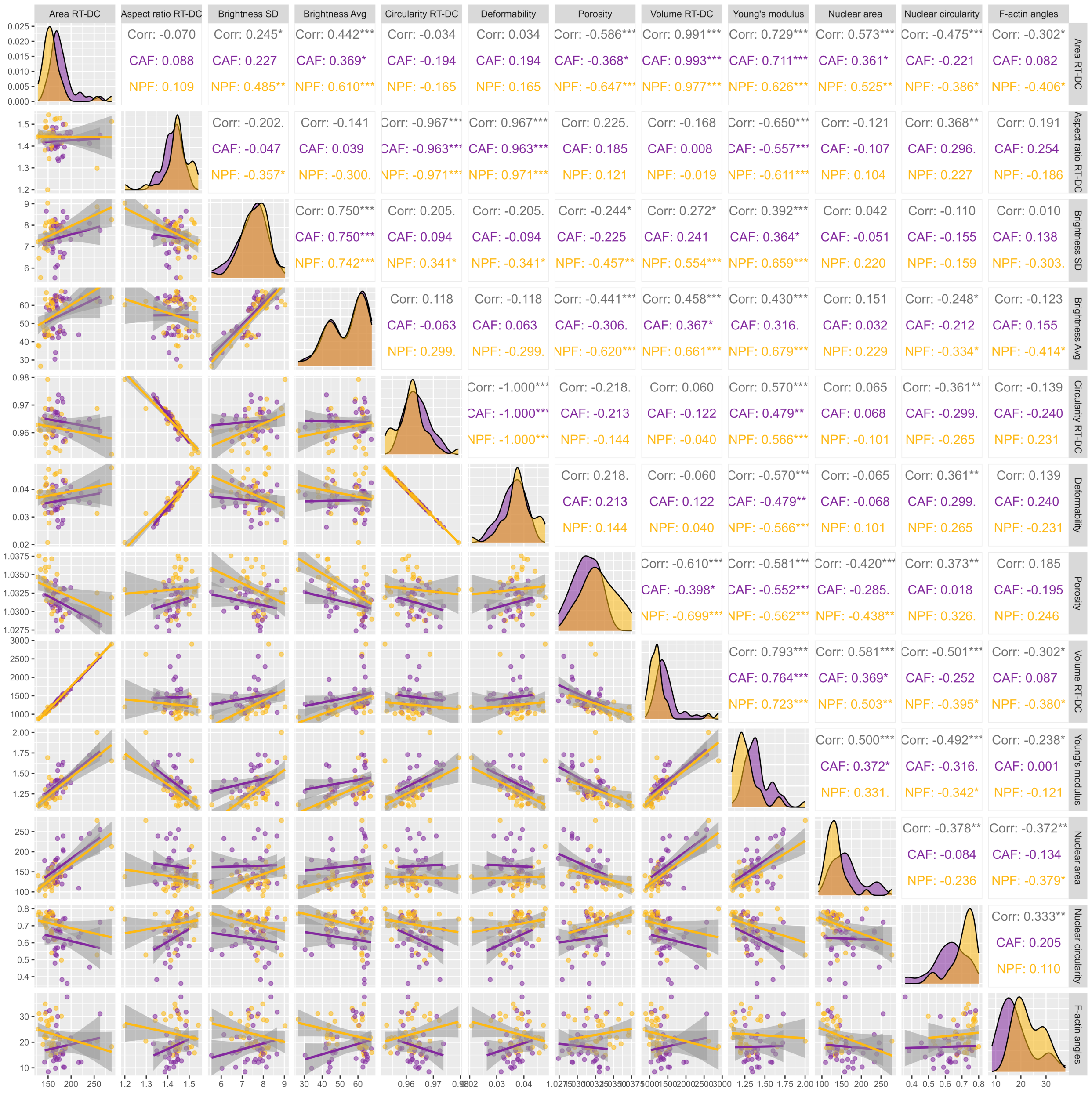

**Supplementary Figure 11. Correlation of additional RT-DC parameters and morphological parameters.** Correlation matrix with scatter plots and histograms of features extracted from RT-DC measurement and morphological parameters. Spearman correlation coefficients with associated p-values are shown (Corr. – all data, both cell types), * *P*<0.05. ** *P*<0.01, *** *P*<0.001. n = 35 donor pairs (medians).

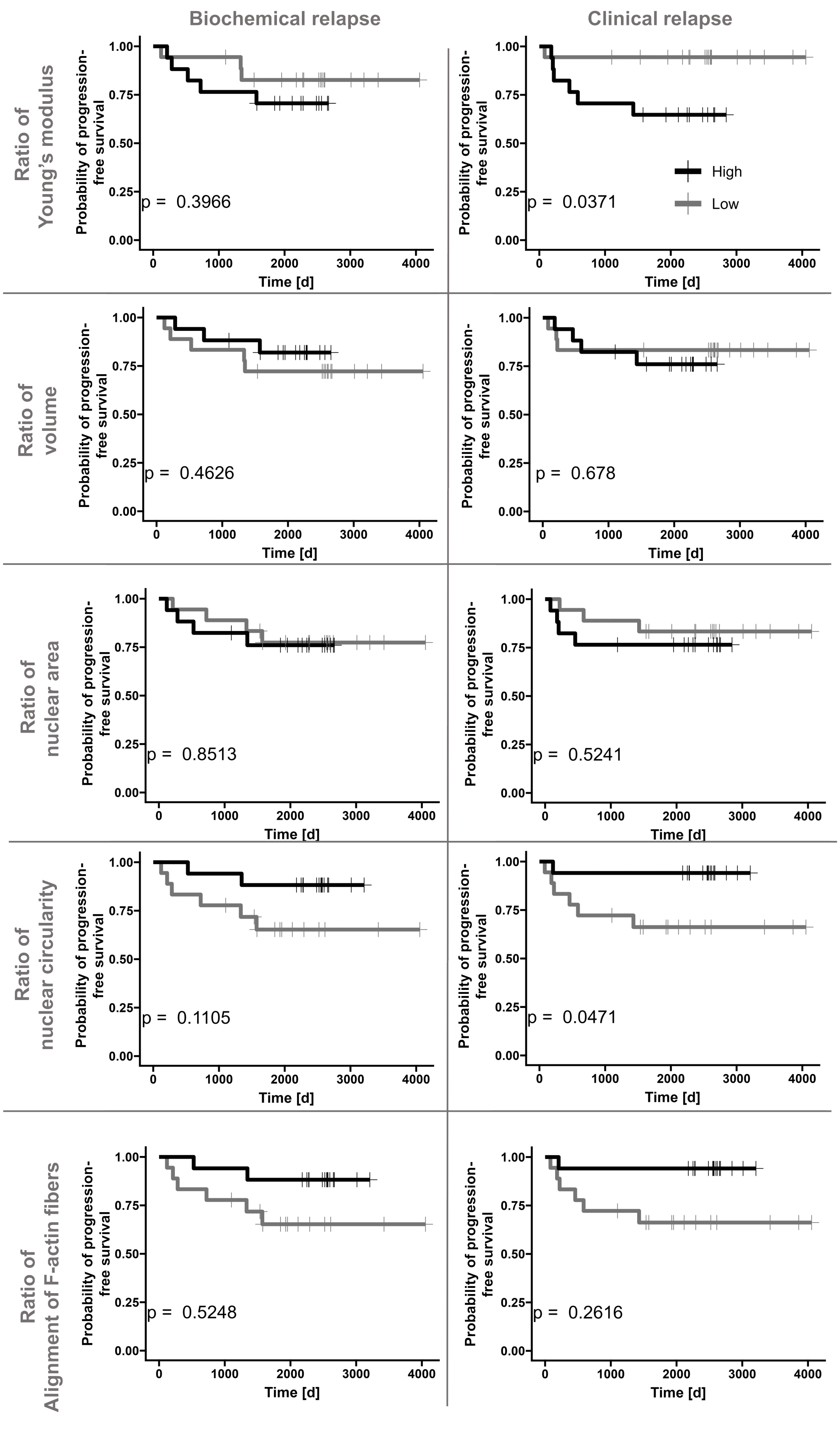


**Supplementary Figure 12. Kaplan-Meier plots of morphological and mechanical parameters.** Kaplan-Meier plots for biochemical (PSA) and clinical relapse for patients with ratios of parameter in the lower (grey, n = 18) and upper 50 percentile (black, n=17). The between-group significance was tested using a log-rank test and is shown in each plot.

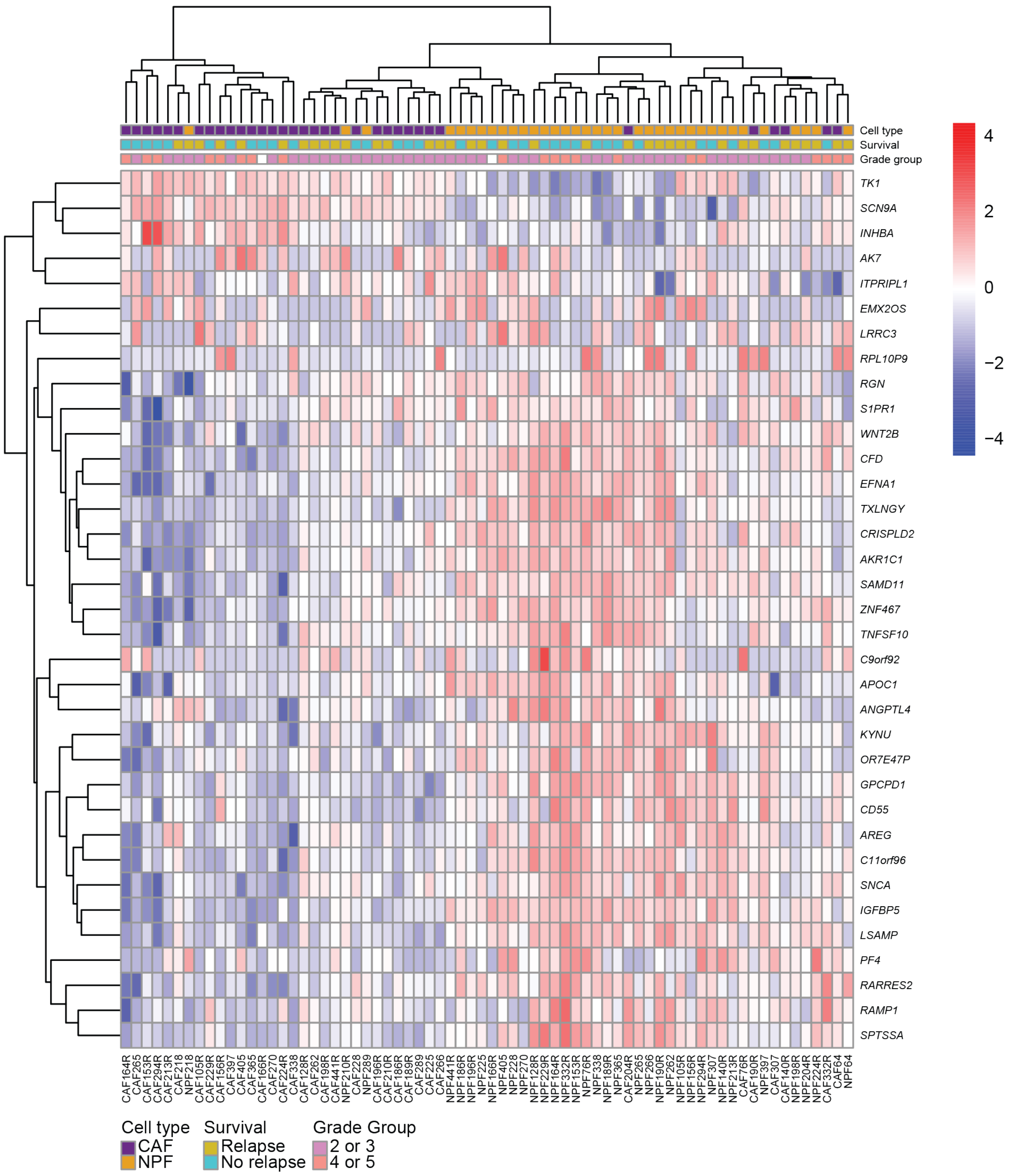


**Supplementary Figure 13. Genes associated with morphological and biomechanical features of CAFs and NPFs.** Heatmap of genes that are significantly correlated with the apparent Young's modulus ratios and/or volume.

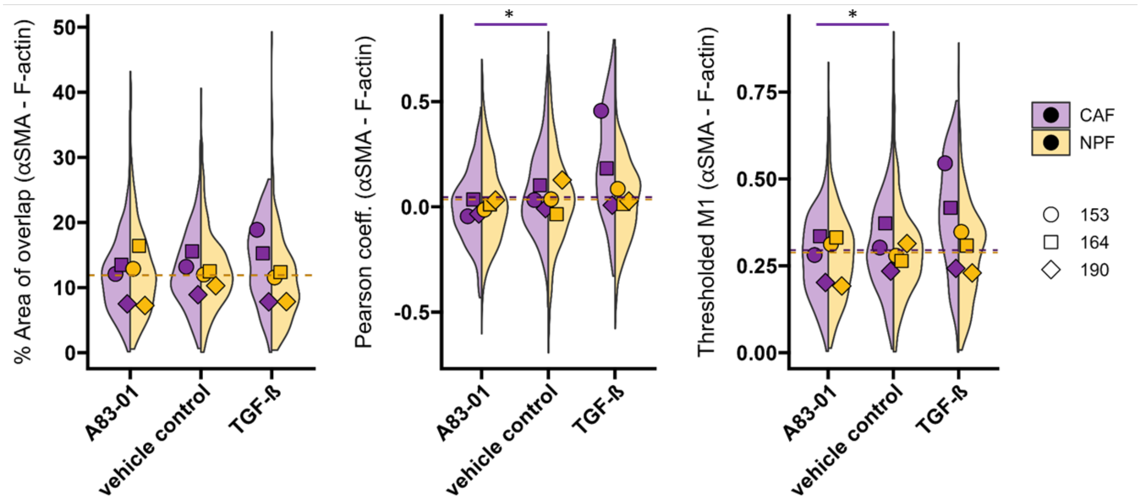


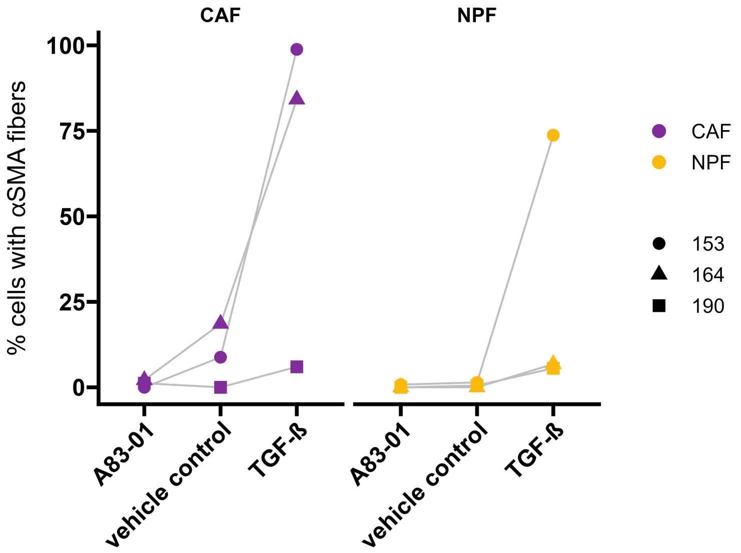


**Supplementary Figure 14. Analysis of αSMA staining after treatment with TGFβ.** Colocalization analysis of αSMA and F-actin was performed with the BIOP JACoP plugin for Fiji (https://github.com/BIOP/ijp-jacop-b). One perinuclear region of interest per cell was analysed with the automatic threshold method Otsu for both channels. Person coefficient, thresholded Manders' coefficient 1 and percentage of overlapping area between αSMA and F-actin are shown for three patients. Percentage of cells with αSMA fibres was determined by counting cells with visible perinuclear fibres and the number of nuclei per image with 3 to 5 images analysed per condition. Medians per patient and cell type are shown. Top: Results of a likelihood-ratio test of linear mixed effects model is shown (% area: CAF, A83-01 vs ctrl: *P*=0.071; CAF TGF-β1 vs ctrl: *P*=0.55; NPF A83-01 vs ctrl: *P*=0.89; NPF TGF-β1 vs ctrl: *P*=0.13; Pearson: CAF: A83-01 vs ctrl: *P*=0.019; CAF TGF-β1 vs ctrl: *P*=0.18; NPF A83-01 vs ctrl: *P*=0.47; NPF TGF-β1 vs ctrl: *P*=0.59; tM1: CAF A83-01 vs ctrl: *P*=0.013; CAF TGF-β1 vs ctrl: *P*=0.14; NPF A83-01 vs ctrl: *P*=0.70; NPF TGF-β1 vs ctrl: *P*=0.70), * p<0.05. Bottom: Results of a Mann-Whitney test are shown (CAF A83-01 vs ctrl: *P*=0.507; CAF TGF-β1 vs ctrl *P*=0.4; CAF TGF-β1 vs A83-01: *P*=0.1; NPF A83-01 vs ctrl: *P*=0.643; NPF TGF-β1 vs ctrl: *P*=0.1; NPF TGF-β1 vs A83-01: *P*=0.076.

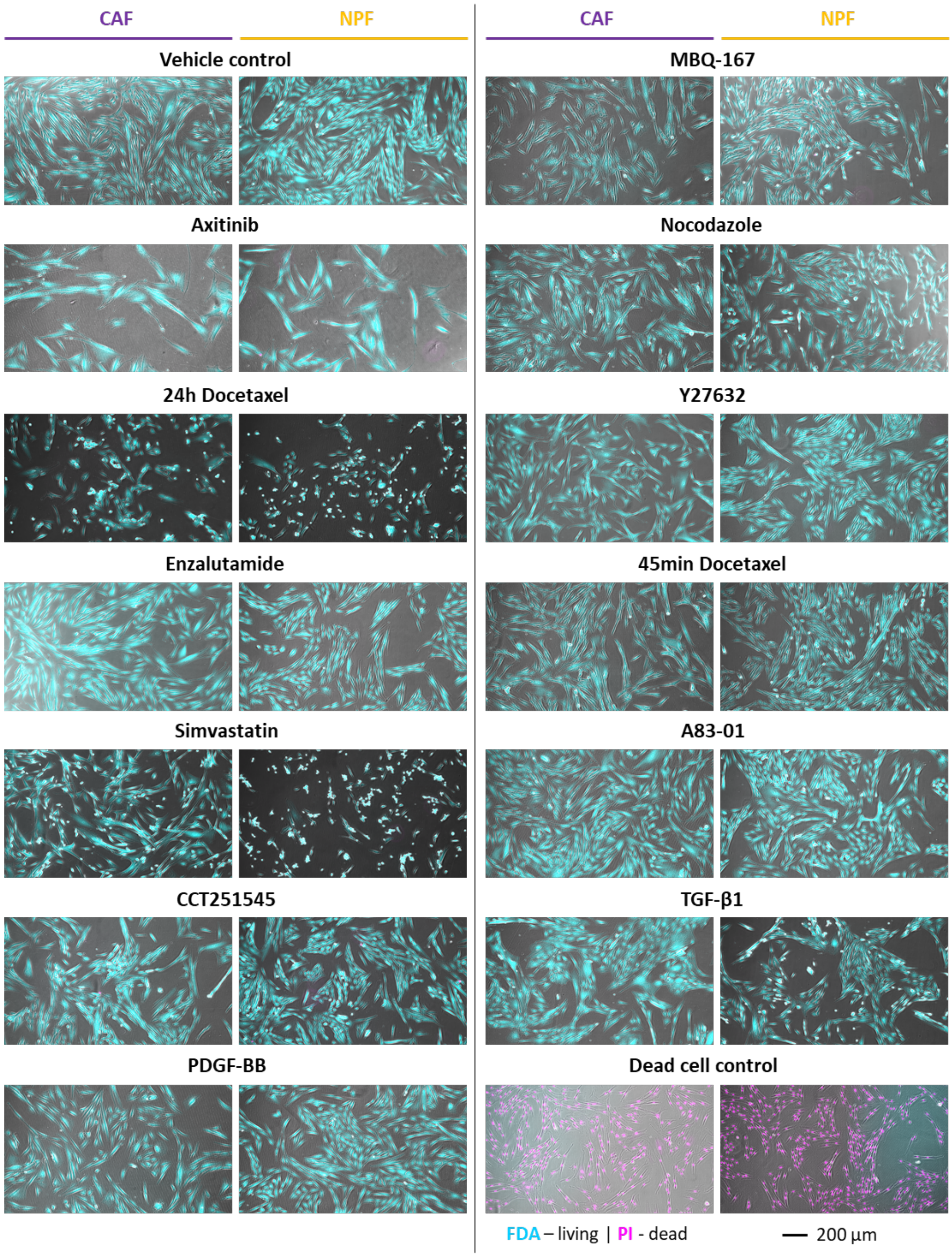


**Supplementary Figure 15. Live/dead staining of cells treated with different drugs.** Epifluorescence imaged of cells treated with different drugs. Cells were treated with the conditions mentioned in figure 6 and 7 (72h 1µM Axitinib; 24h 5nM Docetaxel; 72h 10 µM Enzalutamide; 24h 1µM Simvastatin; 72h 1µM CCT251545; 72h 25ng/mL PDFG-BB; 2h 0.5µM MBQ-167; 45min 5µM Nocodazole; 30min 10µM Y27632; 48h 5µM A83-01; 48h 10 ng/mL TGF-β1; 45min 5µM Docetaxel). After treatment, cells were stained with 8µg/mL fluorescein diacetate (FDA) and 20µg/mL propidium iodide in $CO_2$-independent medium (Gibco) for 5min at room temperature.

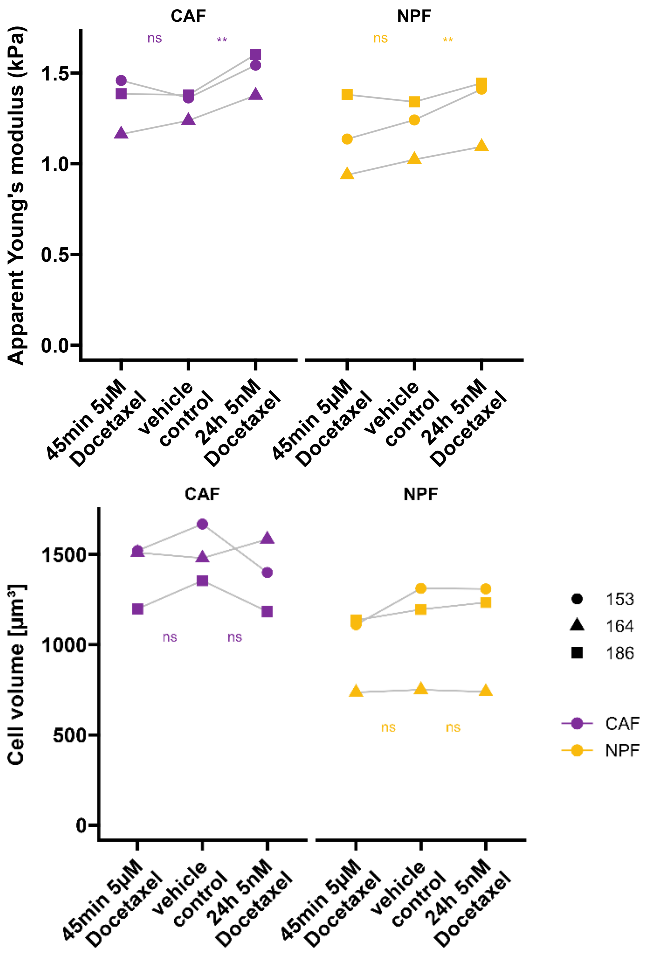


**Supplementary Figure 16. RT-DC measurement of cells treated with Docetaxel for longer time periods (24hrs) or acutely in suspension (45min).** Adherent cells were treated with 5nM Docetaxel or vehicle controls (DMSO 1:1000) for 24h; For acute treatment, cells were detached prior RT-DC measurements (as described in the methods) and treated in suspension with 5µM Docetaxel for 45min. Respective docetaxel concentrations were also present in the RT-DC assay buffer. Results of a likelihood-ratio test of linear mixed effects model is shown. (Volume: CAF 24h Docetaxel $P$=0.4749; CAF 45 min Docetaxel: $P$=0.1542; NPF 24h Docetaxel: $P$=0.2515; NPF 45min Docetaxel: $P$=0.0954; apparent Young's modulus: CAF 24h Docetaxel: $P$=0.0021; CAF 45min Docetaxel: $P$=0.9198; NPF 24h Docetaxel: $P$=0.0067; NPF 45min Docetaxel: $P$=0.2540, ns – non-significant, ** $P$<0.01. n = 3 donors, >9500 cells for each measurement were measured. Medians of one experiment are presented.

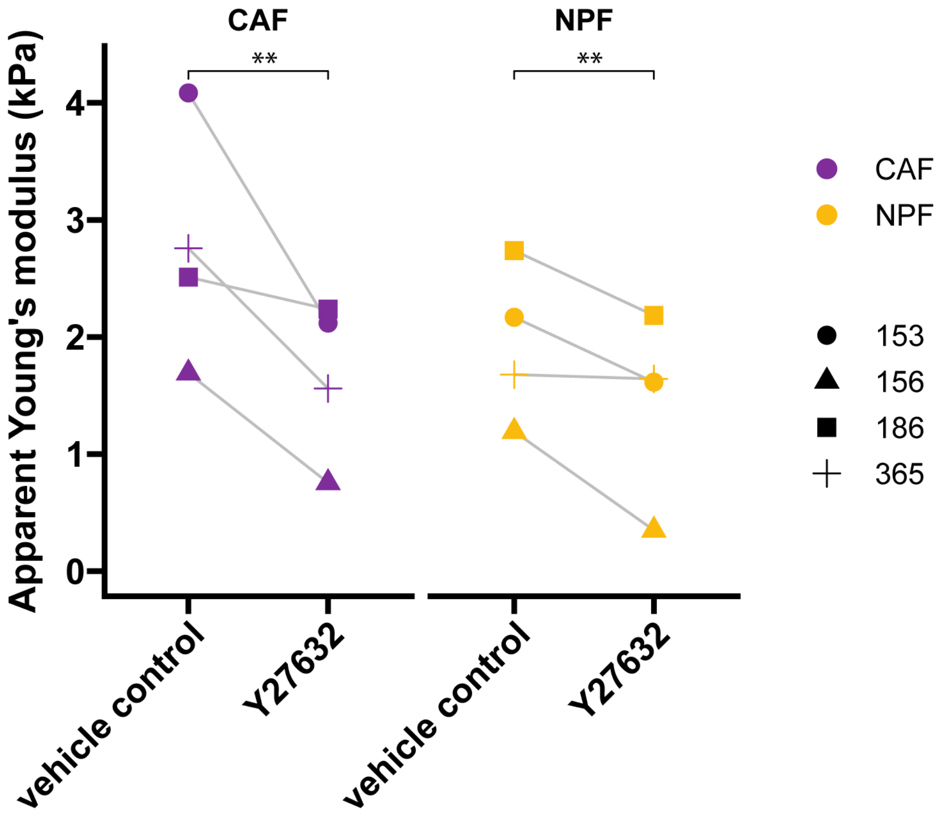


**Supplementary Figure 17. AFM probing of adherent cells after treatment with ROCK inhibitor Y27632.** Cells were seeded into cell culture dishes 24h before measurement. 10µM Y27632 or vehicle control ($H_2O$) were added to the measurement medium ($CO_2$-independent medium) 30 min prior measurements. Cells were probed using a spherical indenter of 5µm diameter (see methods). Results of a likelihood-ratio test of linear mixed effects model is shown (CAF: *P*=0.007; NPF: *P*=0.0024, n = 4 donors, >50 cells for each. Medians are presented.
